# Membrane Environment Sets the Functional pK_a_ of Ionizable Lipids

**DOI:** 10.64898/2026.02.18.706567

**Authors:** Marius F.W. Trollmann, Paolo Rossetti, Rainer A. Böckmann

**Author notes:** Contributing authors.

## Abstract

Ionizable aminolipids enable lipid nanoparticles (LNPs) to encapsulate nucleic acids at neutral pH and to release their cargo upon endosomal acidification. The discrepancy between this effective, acidic LNP pK_a_ and the basic intrinsic pK_a_ of aminolipids, however, remains poorly understood.

Here, we performed microsecond constant-pH molecular dynamics simulations of five widely used aminolipids (DODAP, DLin-MC3-DMA, DLin-KC2-DMA, ALC-0315, and SM-102) embedded in different LNP-relevant ternary DOPC/D-SPC–cholesterol membranes to quantify how aminolipid structure and membrane composition jointly govern aminolipid protonation and the associated pH-dependent membrane remodeling.

Across all systems, membrane embedding lowers the apparent aminolipid pK_a_, yielding physiologically relevant values of 6–7.5 corresponding to shifts by up to 3.5 pK_a_ units or approx. 20 kJ mol***^−^*^1^** with respect to the intrinsic pK_a_. Strikingly, the magnitude of the pK_a_ shift correlates with pH-driven membrane remodeling upon deprotonation: polyunsaturated aminolipids undergo surface-to-core translocation, branched aminolipids preferentially form laterally segregated surface domains, and DODAP remains interfacially anchored through sustained hydration and hydrogen bonding. Saturated helper lipids (DSPC) systematically enhance segregation and amplify pK_a_ shifts relative to DOPC.

Together, these results identify membrane phase behavior as a primary regulator of aminolipid protonation equilibria and establish quantitative design principles for tuning LNP composition toward desired pK_a_, membrane remodeling, and delivery performance.

## Introduction

The delivery of nucleic acids using lipid nanoparticle (LNP) formulations has opened new avenues for the development of innovative therapies and vaccines targeting a wide spectrum of diseases [1]. Structurally, LNPs consist of a hydrophobic core surrounded by a lipid monolayer [1, 2]. The negatively charged nucleic acids can be efficiently encapsulated and stabilized within LNPs through the incorporation of cationic lipids [3].

LNPs typically comprise four different lipid components: Aminolipids, helper lipids, cholesterol, and PEG-lipids, each fulfilling a distinct functional role. These components are subject to diverse optimization strategies, resulting in a broad and highly tunable LNP compositional space. Aminolipids are the predominant component (usually 40-50%) of LNPs [4] and their pH-responsiveness is central to loading or encapsulation efficiency, their bio-distribution, safety, and intracellular payload delivery [5].

Cholesterol is the second most abundant component (≈ 40% in currently reported LNP formulations) and provides support to surface-core organization of LNPs, conferring fluidity while keeping a tight lipid packing preventing payload escape. The substitution of cholesterol with alternative sterols may affect delivery efficiency and favor different tissue tropism, although the precise mechanism is not clear [6]. Notably, chimeric molecules joining the two major LNP components (aminolipid and sterol) achieved satisfactory safety and efficacy profiles both *in-vitro* and *in-vivo* [7] highlighting the untapped potential for exploring novel LNP components.

Helper lipids are usually zwitterionic and bilayer-forming phospholipids. While DSPC is included in all three FDA-approved LNP compositions, sphingomyelin (SM) and ceramides have been proposed as alternative helper lipids in several patents [8, 9]. Importantly, SM lipids have been shown to confer peculiar bio-distribution properties [10], highlighting helper lipid type and molar ratio as critical parameters for further LNP optimization.

PEG-lipids are typically included at low molar fractions (1-2%) to prevent LNP aggregation and prolong their half-life *in vivo* [11]. The lengths of the PEG chain and of the hydrophobic anchor govern PEG desorption kinetics, thereby influencing LNP biodistribution [12]: Shorter hydrophobic anchors favor faster desorption and early formation of the protein corona promoting liver tropism, while bulkier anchors prolong LNP circulation time and tumor accumulation [13] but may cause anti-PEG immunogenicity [14]. Recent developments in the design of novel LNP formulations have removed PEG-lipids entirely from the composition [15].

Early aminolipids were based on quaternary amine headgroups, rendering them permanently cationic. This inherent cationicity facilitates strong electrostatic interactions with nucleic acids, yielding overall positively charged LNP formulations. While such formulations are generally considered beneficial for cellular uptake by promoting interactions with the negatively charged cell surface and glycocalyx [3, 16], permanently cationic lipids have been shown to induce hepatotoxicity, lung toxicity, and result in rapid clearance of LNPs from the bloodstream due to pronounced first-pass effects [17].

These limitations were largely overcome by the development of cationic *ionizable* aminolipids (CILs), which contain secondary or tertiary amine groups capable of undergoing dynamic (de)protonation [3]. CILs are characterized by the following pH-dependent behavior:

- At acidic pH (≈ 4), CILs are protonated, enabling efficient nucleic acid encapsulation [3].
- At physiological pH (7.4), CILs adopt a deprotonated, neutral state, resulting in the formation of LNPs with a lipidic core composed of deprotonated aminolipids and cholesterol, surrounded by a lipid monolayer. In mRNA-loaded LNPs, the negative charge of the nucleic acids is shielded by locally protonated aminolipids [2, 18].
- Upon endosomal acidification (≈ 6), CILs re-protonate at the LNP surface [18], enhancing electrostatic interactions with anionic endosomal lipids [19]. In addition, the bilayer-destabilizing properties of CILs likely promote membrane fusion and thereby nucleic acid release [1].

Accordingly, the protonation behavior of aminolipids is central to LNP function. It is determined by the *intrinsic* pK_a_ of the aminolipid defined as the pK_a_ of the functional ionizable group in infinite dilution conditions (pK_a_*^AL^*). Experimentally, pK_a_*^AL^* values are typically determined by potentiometric titration of water-soluble aminolipid analogs and can be tuned by modifying the distance between the ionizable nitrogen and the electron-withdrawing groups (Fig. 1) [5].

**Fig. 1.**
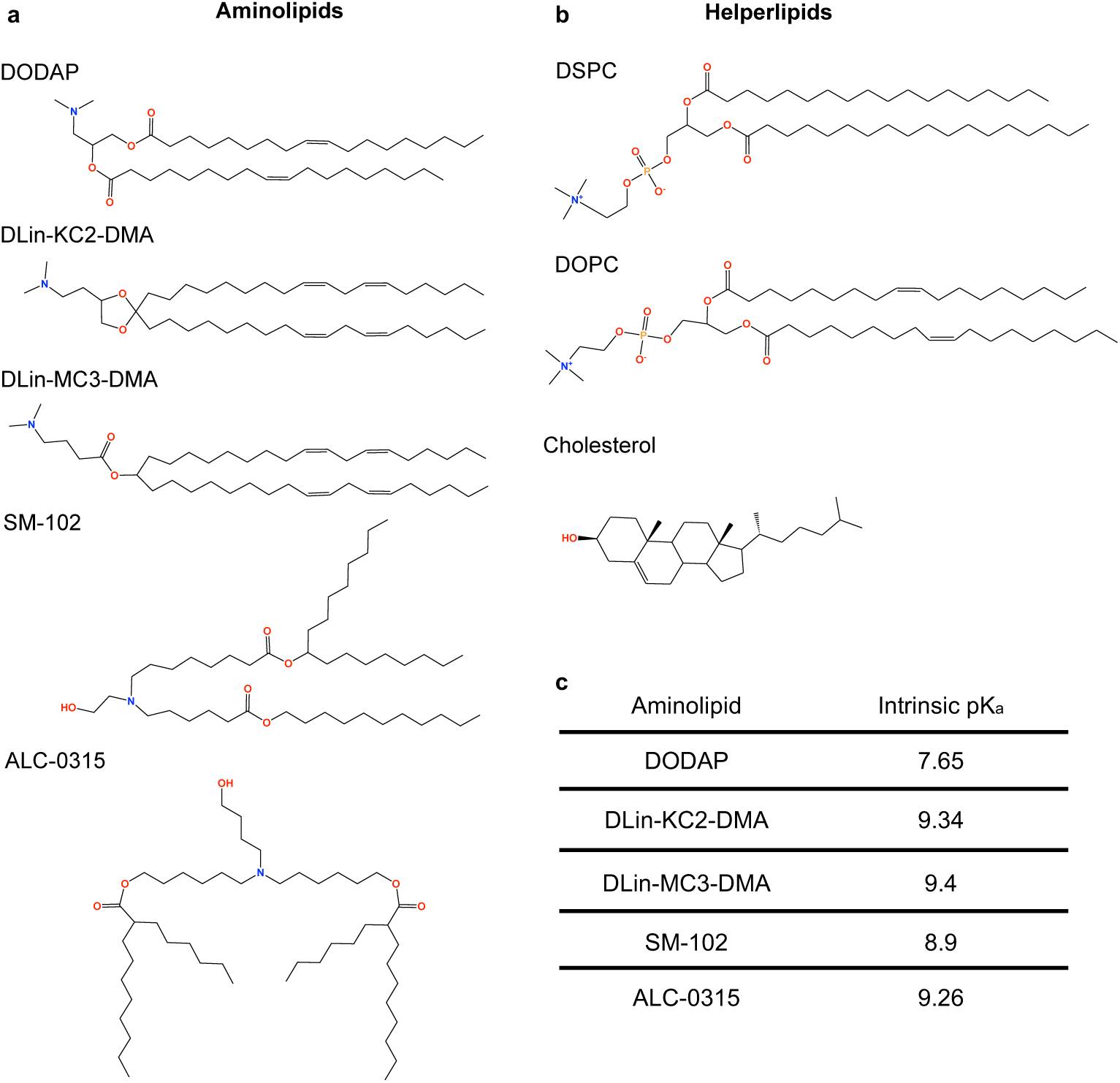
**a** Chemical structures of the aminolipids investigated in this study. **b** Helper phospholipids used in model membranes. **c** Intrinsic pK_a_ values of DODAP, DLin-KC2-DMA, DLin-MC3-DMA [^21^], ALC-0315 [4, 31], and SM-102 [4, 31].

In contrast, the *apparent* pK_a_ of full LNPs (pK_a_*^LNP^*) depends not only on the intrinsic pK_a_*^AL^*, but additionally on the local LNP composition, structure, and molecular organization, and on the particular method employed to measure the pK_a_*^LNP^*. While reported intrinsic pK_a_*^AL^* values range from 7 to 10 (Fig. 1c), effective LNP formulations consistently exhibit pK_a_*^LNP^* values between 6 and 7, i.e., consistently slightly below physiological pH.

A widely used approach to determine pK_a_*^LNP^* is the TNS binding assay [20], which monitors fluorescence changes upon TNS partitioning into LNP membranes. Because TNS binding increases with membrane positive charge – and thus aminolipid protonation – this method provides an indirect measure of membrane charge. We previously demonstrated that TNS localizes predominantly within the LNP monolaye shell; consequently, pK*^LNP^*values obtained from TNS assays primarily reflect aminolipid protonation in the LNP shell rather than in the core [18]. Accordingly, TNS assays do not allow to sense the protonation states of aminolipids residing in the LNP core [5, 21]. This limitation has recently been addressed using DNA-based fluorescent probes, which revealed that the LNP core is highly permeable to protons [22].

Beyond intrinsic and LNP-level pK_a_ values, one may further define the *apparent* aminolipid pK_a_ as the pH at which the probability of aminolipid protonation equals that of deprotonation within a given environment. This apparent pK_a_*^AL^*depends on the local lipid environment, including surrounding lipid species and aminolipid density. Classical molecular dynamics (MD) simulations have been successfully employed to estimate apparent pK*^AL^* values in defined environments [4]; however, such approaches neglect cooperative protonation effects and typically rely on predefined protonation states. In contrast, constant-pH molecular dynamics (CpHMD) simulations [23] enable unbiased sampling of dynamic protonation equilibria of aminolipids as a function of the local environment [18, 24].

The function of aminolipids is strongly influenced by both their intrinsic pK_a_ and their molecular shape [25]. While the headgroup chemistry is decisive for the intrinsic pK_a_, the number and saturation of fatty acyl chains govern membrane embedding and aminolipid shape, which in turn modulate the pK_a_ shift, fusogenicity, and membrane remodeling.

The aminolipids most frequently employed in LNP formulations include (Fig. 1):

- **DODAP** (1,2-dioleoyl-3-dimethylammonium propane), which features two oleyl chains linked by a 1,3-propanediol backbone and a tertiary amine headgroup.
- **MC3** (DLin-MC3-DMA) possesses two linoleyl tails and just one ester group, conferring enhanced hydrophobicity and bilayer-destabilizing properties. MC3 belongs to the *Patisiran* (*Onpattro*) LNP composition, the first FDA-approved siRNA and LNP-based therapeutic, used to treat hereditary transthyretin-mediated amyloidosis [26].
- **KC2**(DLin-KC2-DMA) shares the acyl chains of MC3 but has a different headgroup domain, characterized by a dioxolane ring.
- **ALC-0315**, a next-generation, four-tailed aminolipid introduced following the pioneering work of Akinc *et al.* [27]. ALC-0315 is a component of the Pfizer SARS-CoV-2 mRNA vaccine and has demonstrated improved siRNA delivery relative to MC3 in preclinical studies [28].
- **SM-102**, the aminolipid used in the Moderna SARS-CoV-2 vaccine, is characterized by a branched three-tail architecture and a slightly lower intrinsic pK_a_ due to the shorter distance between the hydroxyl group and the tertiary amine.

Contemporary aminolipid design increasingly combines multi-tail architectures with unsaturations, cyclohexyl moieties, and anteiso-branched chains [8, 9, 29, 30], reflecting continued efforts to expand the aminolipid chemical space. In this work, we investigate how aminolipid structure and lipid environment jointly influence protonation equilibria and pH-dependent membrane remodeling. Specifically, five well-characterized aminolipids (see above) spanning different stages of structural evolution are studied in membranes with varying helper phospholipid composition and cholesterol content using microsecond-scale all-atom CpHMD simulations. Our analysis focuses on aminolipid pK_a_ shifts within ternary membranes of differing acyl chain saturation and cholesterol concentration, as well as on the pH-dependent membrane structure and organization.

## Results

This study examines how lipid membrane composition modulates the protonation equilibria of embedded aminolipids and, conversely, how pH-dependent aminolipid protonation affects membrane organization. Symmetric lipid bilayers were composed of 20% aminolipids, 60% (or 40%) of either fully saturated DSPC or diunsaturated DOPC phospholipids, and 20% (or 40%) cholesterol. These compositions were selected to ensure membrane stability across the investigated pH range (3 – 11).

Each system contained 240 lipid and cholesterol molecules at a hydration level of approximately 50 water molecules per lipid (or cholesterol molecule). System neutrality was maintained using a pH-dependent number of chloride ions and buffer particles. The total aggregate simulation time amounted to approximately 0.26 ms. Details of aminolipid parameterization, system setup, and analysis are provided in the Methods section.

### pK_a_ shifts of aminolipids in ternary membranes

The pH dependence of aminolipid protonation was quantified by computing the mean fraction of deprotonated molecules, *S^deprot^* (Fig. 2), over a pH range from 3 to 11. For all aminolipids and membrane compositions, the resulting titration curves were well described by a generalized Henderson-Hasselbalch equation. Fitting the averaged *S^deprot^* values yielded the apparent aminolipid pK_a_ values, as well as the associated cooperativity parameters *n* (Tab. 1).

**Fig. 2.**
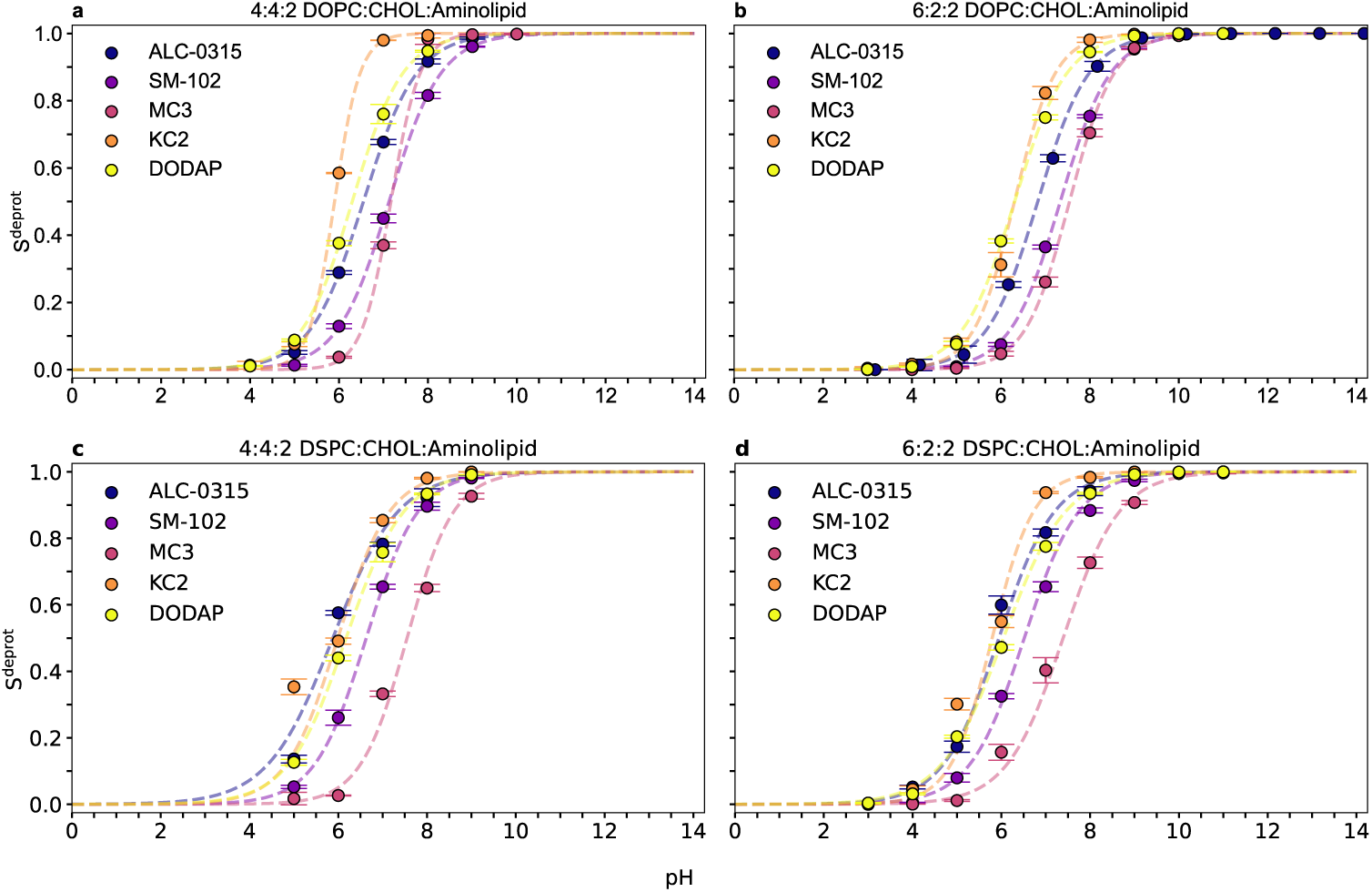
Titration curves of aminolipids in different membrane compositions. Titration curves are shown for DOPC-based (*top*) and DSPC-based (*bottom*) membranes containing **a**, **c** 40 mol% cholesterol and **b**, **d** 20 mol% cholesterol. The aminolipid content was fixed at 20 mol% in all systems. Error bars denote the standard error of the mean, obtained by block averaging (see Methods). The data were fitted using the generalized Henderson-Hasselbalch equation, and the resulting fit parameters are summarized in Tab. 1. The values of *S^deprot^* as a function of time are shown in Supplementary Figures 49 – 52.

**Table 1.**
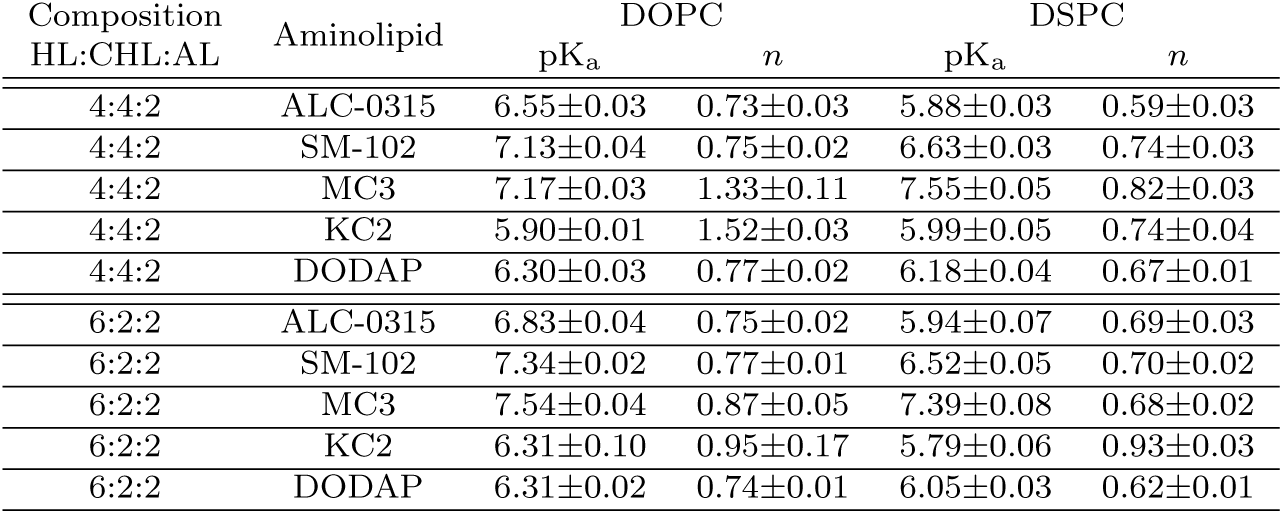
Apparent pK_a_ and cooperativity (*n*) values estimated from fits to the averaged degree of aminolipid deprotonation as a function of pH (Fig. 2). Errors denote the 95% confidence interval calculated via parametric bootstrapping.

| Composition<br>HL:CHL:AL | Aminolipid | DOPC |  | DSPC |  |
| --- | --- | --- | --- | --- | --- |
|  |  | pK<sub>a</sub> | *n* | pK<sub>a</sub> | *n* |
| 4:4:2 | ALC-0315 | 6.55±0.03 | 0.73±0.03 | 5.88±0.03 | 0.59±0.03 |
| 4:4:2 | SM-102 | 7.13±0.04 | 0.75±0.02 | 6.63±0.03 | 0.74±0.03 |
| 4:4:2 | MC3 | 7.17±0.03 | 1.33±0.11 | 7.55±0.05 | 0.82±0.03 |
| 4:4:2 | KC2 | 5.90±0.01 | 1.52±0.03 | 5.99±0.05 | 0.74±0.04 |
| 4:4:2 | DODAP | 6.30±0.03 | 0.77±0.02 | 6.18±0.04 | 0.67±0.01 |
| 6:2:2 | ALC-0315 | 6.83±0.04 | 0.75±0.02 | 5.94±0.07 | 0.69±0.03 |
| 6:2:2 | SM-102 | 7.34±0.02 | 0.77±0.01 | 6.52±0.05 | 0.70±0.02 |
| 6:2:2 | MC3 | 7.54±0.04 | 0.87±0.05 | 7.39±0.08 | 0.68±0.02 |
| 6:2:2 | KC2 | 6.31±0.10 | 0.95±0.17 | 5.79±0.06 | 0.93±0.03 |
| 6:2:2 | DODAP | 6.31±0.02 | 0.74±0.01 | 6.05±0.03 | 0.62±0.01 |

**Table 2.** All-atom molecular dynamics simulations using the constant-pH method to parametrize and validate *V ^MM^* (*λ*).

| System | Aminolipid:H<sub>2</sub>O:Na<sup>+</sup>:Cl<sup>-</sup>:Buf | Sim. Time (μs) | pH | Objective |
| --- | --- | --- | --- | --- |
| DODAP<sub>A</sub> | 1:6945:19:20:1 | 0.6 | - | Thermodynamic Integration |
| DLin-MC3-DMA<sub>A</sub><sup>a</sup> | 1:6944:19:20:1 | 0.6 | - | Thermodynamic Integration |
| DLin-KC2-DMA<sub>A</sub> | 1:6948:19:20:1 | 0.6 | - | Thermodynamic Integration |
| SM-102<sub>A</sub> | 1:6941:19:20:1 | 1.7 | - | Thermodynamic Integration |
| ALC-0315<sub>A</sub><sup>b</sup> | 1:6938:19:20:1 | 1.7 | - | Thermodynamic Integration |
| DODAP<sub>B</sub> | 1:6944:19:20:3 | 0.6 | - | Goodness of polynomial coefficients |
| DLin-MC3-DMA<sub>B</sub><sup>a</sup> | 1:6943:19:20:3 | 0.3 | - | Goodness of polynomial coefficients |
| DLin-KC2-DMA<sub>B</sub> | 1:6947:19:20:3 | 0.6 | - | Goodness of polynomial coefficients |
| SM-102<sub>B</sub> | 1:6940:19:20:3 | 1.0 | - | Goodness of polynomial coefficients |
| ALC-0315<sub>B</sub><sup>b</sup> | 1:6937:19:20:3 | 1.7 | - | Goodness of polynomial coefficients |
| DODAP<sub>C</sub> | 1:6944:19:20:3 | 10 × 0.6 | 5–14 | Intrinsic pK<sub>a</sub> |
| DLin-MC3-DMA<sub>C</sub><sup>a</sup> | 1:6943:19:20:3 | 10 × 0.3 | 5–14 | Intrinsic pK<sub>a</sub> |
| DLin-KC2-DMA<sub>C</sub> | 1:6947:19:20:3 | 10 × 0.6 | 5–14 | Intrinsic pK<sub>a</sub> |
| SM-102<sub>C</sub> | 1:6940:19:20:3 | 10 × 1.0 | 5–14 | Intrinsic pK<sub>a</sub> |
| ALC-0315<sub>C</sub><sup>b</sup> | 1:6937:19:20:3 | 10 × 1.0 | 4.17–14.17 | Intrinsic pK<sub>a</sub> |

<sup>a</sup>Equivalent to [24].
<sup>b</sup>Equivalent to [18]

**Table 3.**
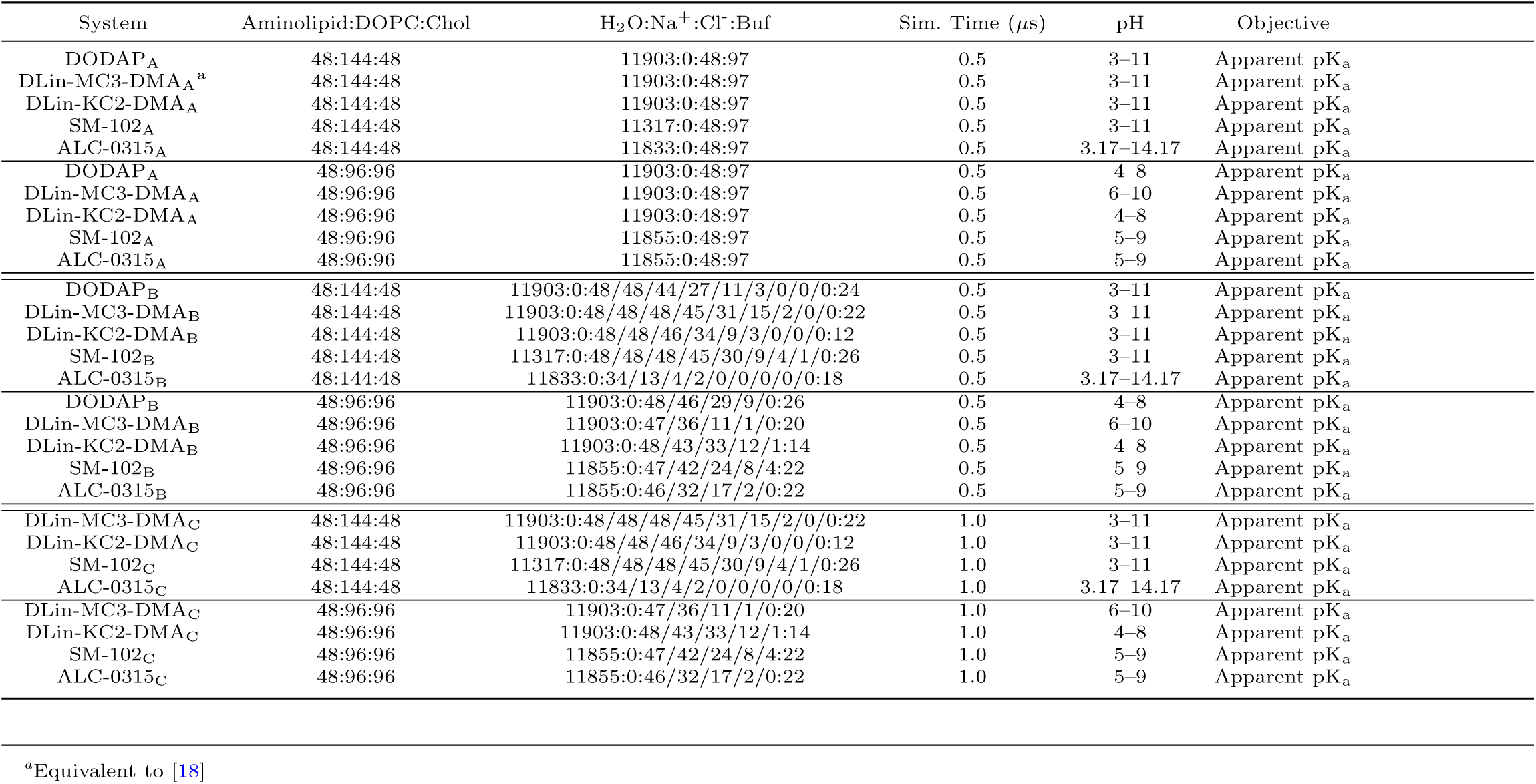
All-atom molecular dynamics simulations of ternary membranes including the helperlipid DOPC using the constant-pH method.

Across all systems, the apparent pK_a_ values were consistently lower than the intrinsic pK_a_ values of the corresponding aminolipids at infinite dilution (Fig. 3a,b), indicating a destabilization of the protonated state within membrane environments. The magnitude of the pK_a_ shift depended on both membrane composition and aminolipid structure. The largest shift was observed for KC2 in DOPC-based membranes at a 40:40:20 DOPC:cholesterol:KC2 composition, with Δ*pK*_a_ = 3.55, corresponding to a protonation free-energy change of Δ*G_a_* = ln 10 · Δ*pK*_a_ · *RT* ≈ 21.1 kJ mol*^−^*^1^. In contrast, the smallest shift was found for DODAP (Δ*pK*_a_ = 1.35) in a 40:40:20 DOPC:cholesterol:DODAP membrane (Fig. 3a,b).

**Fig. 3.**
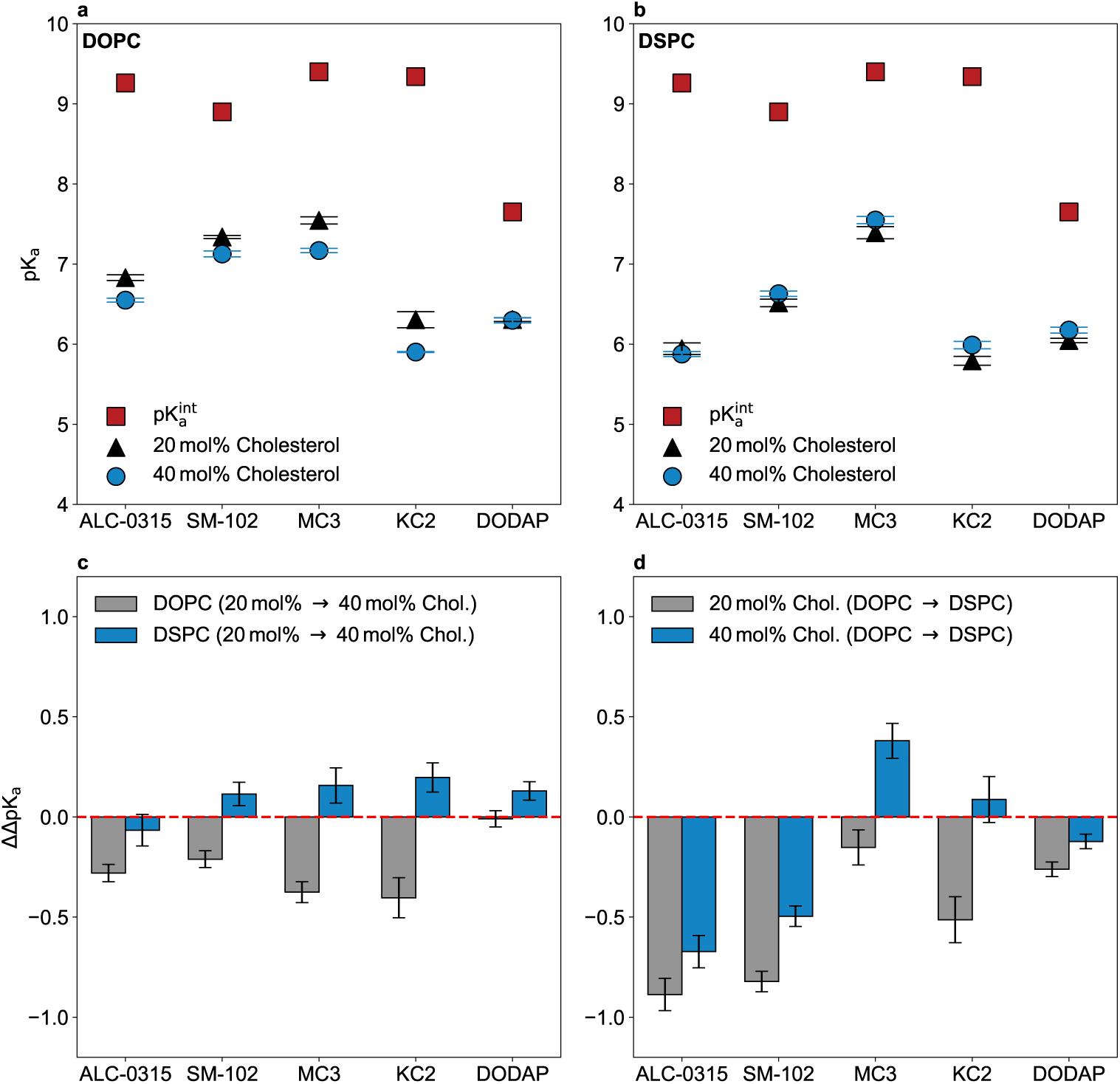
Influence of membrane composition on aminolipid pK_a_. Comparison of apparent pK_a_ values for **a** DOPC- and **b** DSPC-based membranes containing 20 mol% or 40 mol% cholesterol with the intrinsic pK_a_ values at infinite dilution. **c** and **d** show differential pK_a_ shifts (ΔΔ*pK*a), where Δ*pK*a = *pK*a^app^ *−pK*a^int^ and ΔΔ*pK*a denotes the difference between two Δ*pK*a values. Specifically, **c** quantifies the effect of cholesterol content (40 mol% *−* 20 mol%) for the different aminolipids, shown separately for DOPC- and DSPC-based membranes, whereas **d** reports the effect of helper lipid identity (DSPC – DOPC) at fixed cholesterol fractions (20 mol% and 40 mol%). Error bars indicate 95% confidence intervals: for **a** and **b**, intervals were obtained by bootstrap resampling of the titration fits (see Methods); for **c** and **d**, confidence intervals were calculated by propagation of the bootstrap-derived standard errors, assuming independent estimates.

The influence of cholesterol on the pK_a_ shift differed between unsaturated (DOPC) and saturated (DSPC) membranes. In DOPC systems, increasing cholesterol content led to larger pK_a_ shifts, whereas in DSPC systems the effect was weaker and, for all aminolipids except ALC-0315, reversed (Fig. 3c). For most compositions, replacing DOPC with DSPC resulted in an increased pK_a_ shift (more negative), an effect that was particularly pronounced for branched aminolipids, with increases of approximately 0.5-1 pK_a_unit. Only for the polyunsaturated aminolipids KC2 and MC3 at high cholesterol content was a slight increase in the apparent pK_a_ observed upon DOPC→DSPC substitution. Notably, these aminolipids also exhibited the strongest sensitivity to cholesterol concentration (Fig. 3c).

### pH-driven (de-)mixing of ternary membranes containing aminolipids

The response of the ternary membranes to changes in environmental pH depends strongly on both membrane composition and aminolipid structure. pH-induced changes in membrane organization were quantified using the normalized average conditional mixing entropy, *S^mix^* (Fig. 4) [32]. For all compositions, *S^mix^* reached a maximum at acidic pH (pH 3), corresponding to homogeneously mixed bilayers containing phospholipids, aminolipids, and cholesterol. With increasing pH, several membrane compositions exhibited pronounced de-mixing (*S^mix^ <* 1), resulting in the formation of compositionally distinct phases.

**Fig. 4.**
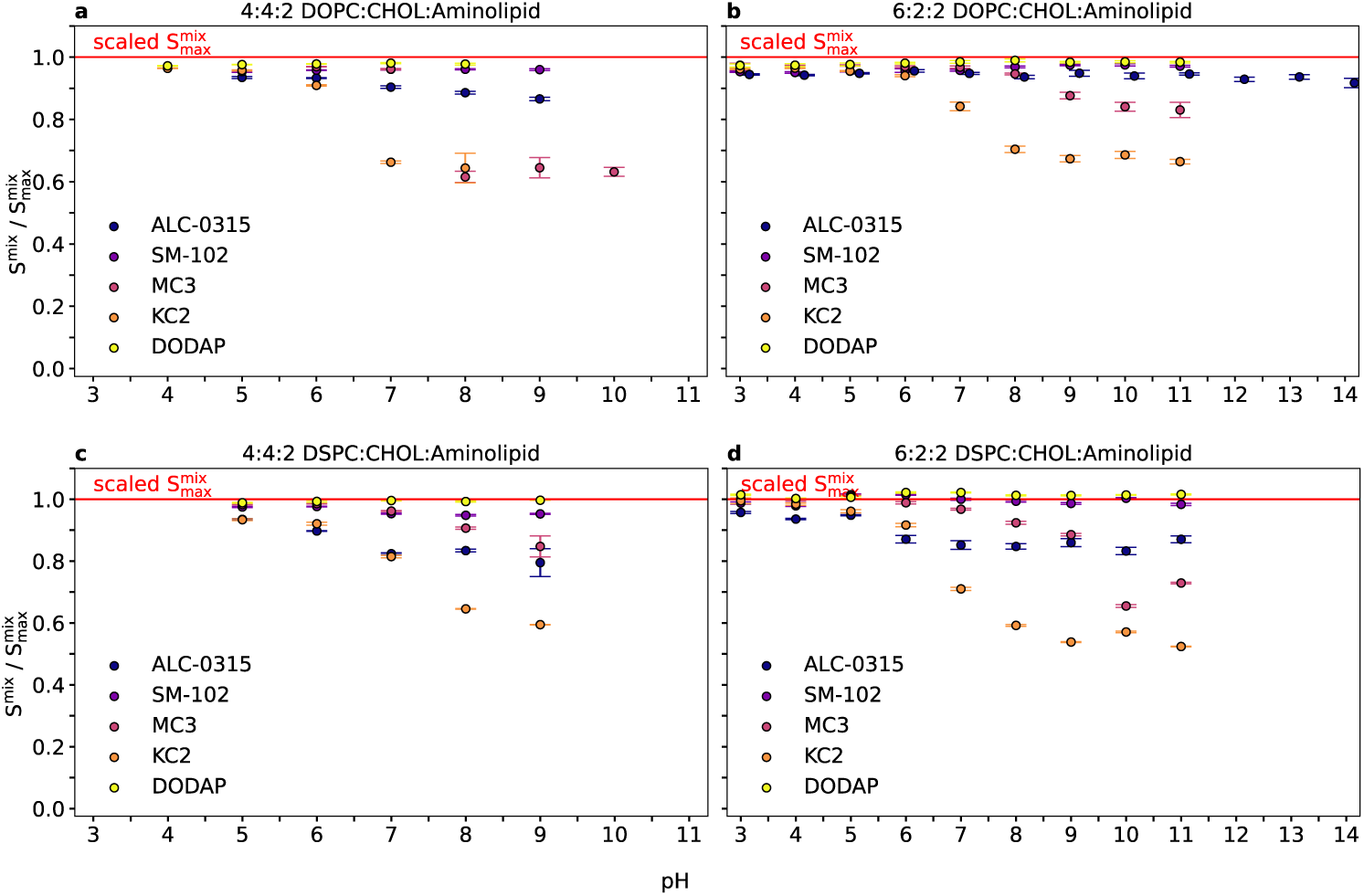
Standardized mixing entropy (*S^mix^*) as a function of pH. Mean values of *S^mix^*are shown for DOPC-based (top) and DSPC-based (bottom) membranes containing **a**, **c** 40 mol% cholesterol and **b**, **d** 20 mol% cholesterol. Conditional mixing entropies were calculated following Brandani *et al.* [32], using a 1 nm cutoff to define neighboring lipids. Contacts were defined based on the distance between reference atoms in the headgroups of the target lipids: the titratable nitrogen for aminolipids, the oxygen atom for cholesterol, and the sn-2 carbon atom for phospholipids. Error bars indicate the standard error of the mean, obtained by block averaging (see Methods). The values of *S^mix^* as a function of time are shown in Supplementary Figures 53 – 56.

Neighbor analysis using a 1 nm cutoff (Fig. 5) revealed that this de-mixing originates from preferential segregation of aminolipids from phospholipids and cholesterol. The overall tendency toward segregation followed the order KC2 (highest segregation) *>* MC3 *>* ALC-0315 *>* SM-102 *>* DODAP (lowest segregation) and was most pronounced in membranes containing higher fractions of DSPC. For the polyunsaturated aminolipids KC2 and MC3, increasing pH induced aminolipid migration *into* the hydrophobic membrane core, where they localized between the phospholipid leaflets together with a fraction of the cholesterol population (Fig. 6), consistent with previous LNP-mimetic studies [2, 33, 34]. In contrast, branched aminolipids (ALC-0315 and SM-102) predominantly underwent *lateral* segregation from surrounding phospholipids, an effect that was strongest for the four-tailed ALC-0315 in DSPC-containing membranes (Fig. 7). Migration of aminolipids toward the membrane interior has also been reported for ALC-0315 and SM-102 in standard LNP formulations [2, 18, 35]. Over the investigated timescale of 1 *µ*s, DODAP showed little to no segregation under any conditions, consistent with its high mixing entropy (Fig. 4). Representative snapshots of all systems after 2 *µ*s (or 1 *µ*s) are shown in Supplementary Figures 7 – 44.

**Fig. 5.**
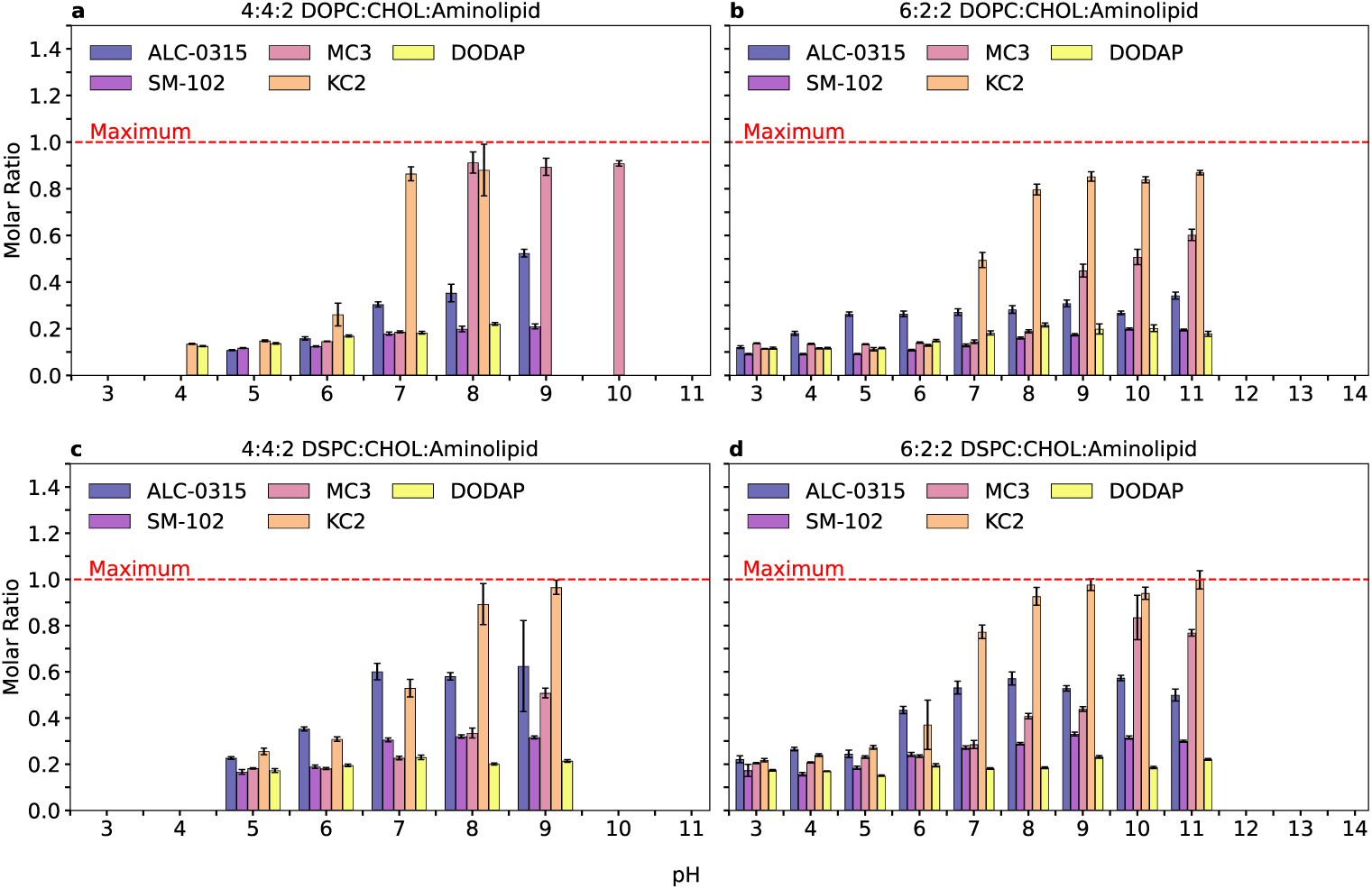
Local pH-dependent lipid environment of aminolipids. Fraction of aminolipid–aminolipid contacts within 1 nm, averaged over all aminolipids and normalized by the total number of contacts around aminolipids, as a function of pH for membranes containing **a**, **b** DOPC or **c**, **d** DSPC. Fraction of contacts between aminolipids-phospholipids and aminolipids-cholesterol are omitted for clarity. Contacts were defined based on the distance between reference atoms in the headgroups of the target lipids: the titratable nitrogen for aminolipids, the oxygen atom for cholesterol, and the sn-2 carbon atom for phospholipids. Error bars indicate the standard error of the mean, obtained by block averaging (see Methods). The average number of neighboring lipids per aminolipid as a function of time are shown in Supplementary Figures 61 – 66.

**Fig. 6.**
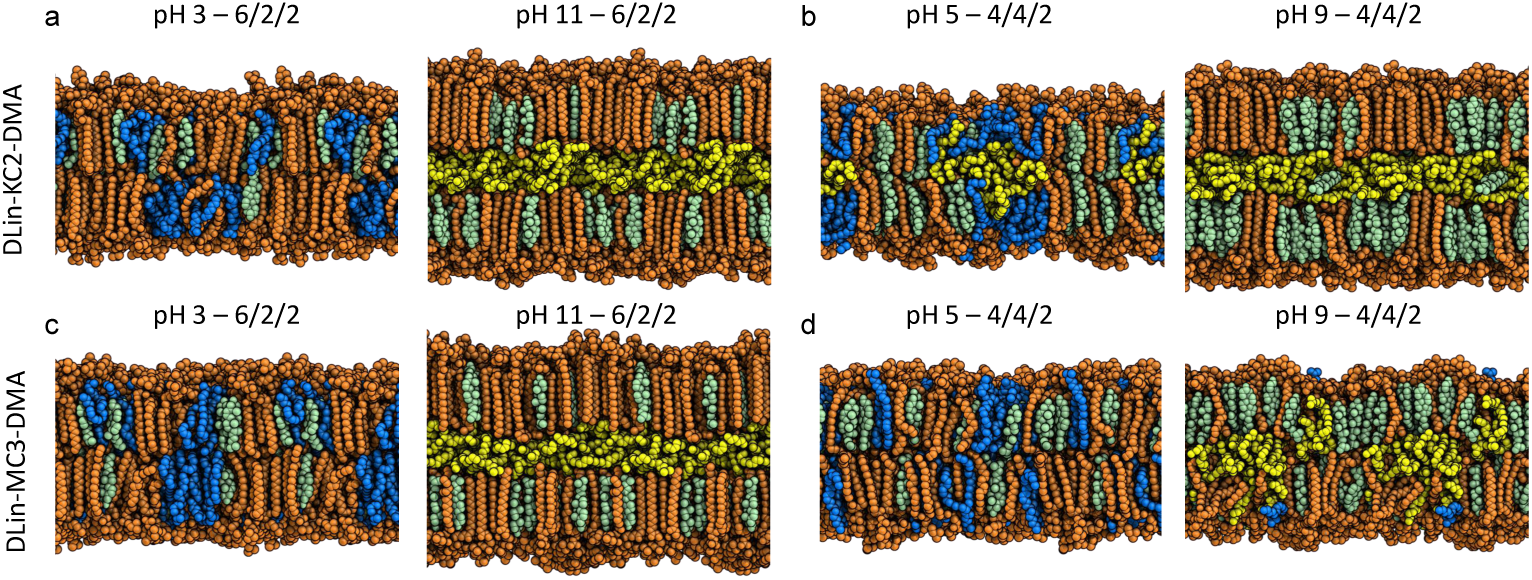
Side views of DSPC:cholesterol:MC3/KC2 membranes. Representative snapshots of DSPC-based membranes after 2 *µ*s containing DLin-KC2-DMA (*top row*) or DLin-MC3-DMA (*bottom row*) at cholesterol fractions of 20 mol% (**a**, **c**) and 40 mol% (**b**, **d**). DSPC lipids are shown in *orange*, cholesterol in *green*. At low pH, protonated aminolipids (*blue*) reside at the membrane surface, whereas at higher pH the deprotonated aminolipids (*yellow*) migrate into the membrane core. Figures were rendered with PyMOL [36].

**Fig. 7.**
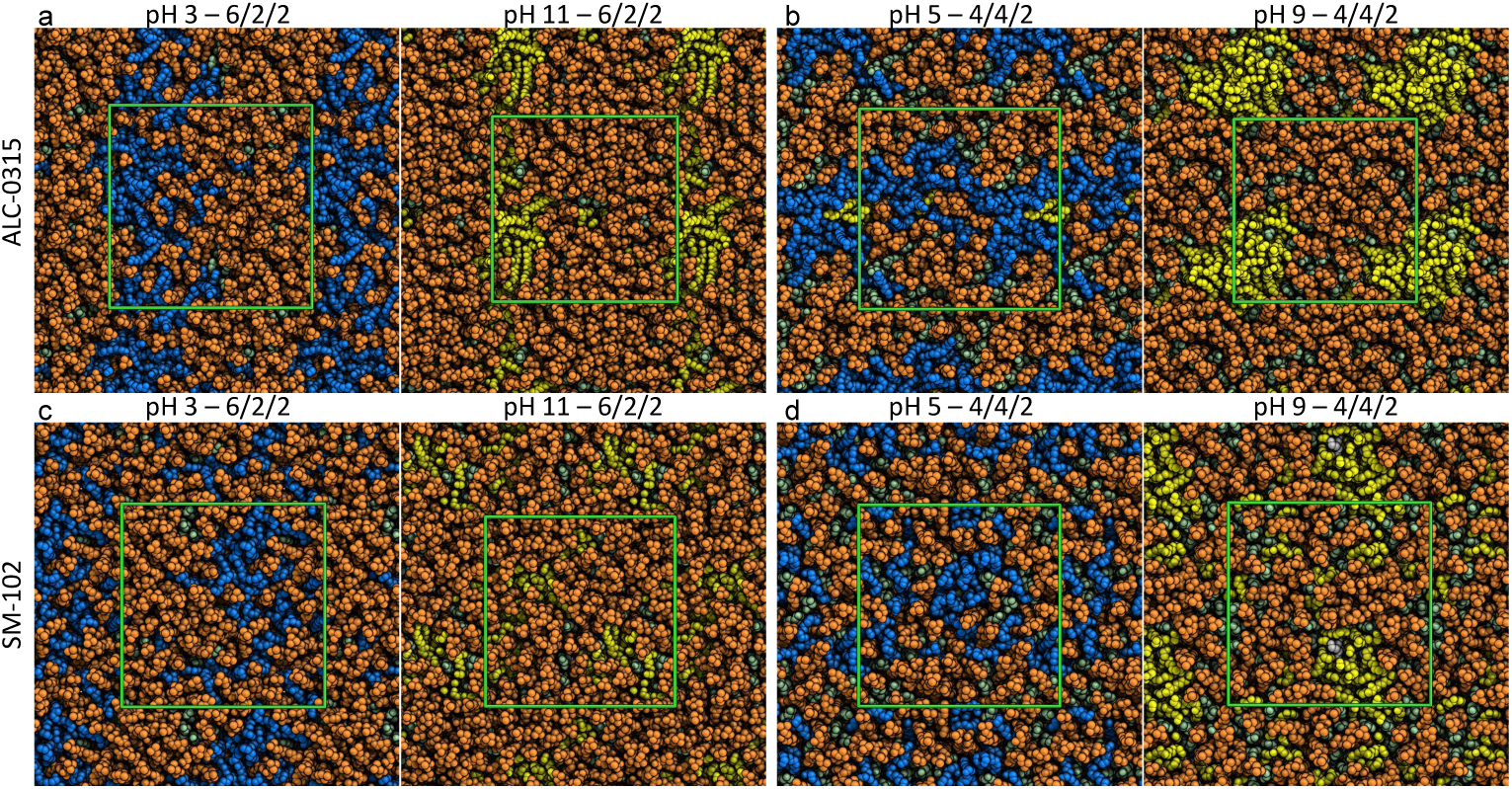
Top views of DSPC:cholesterol:ALC-0315/SM-102 membranes. Representative snapshots of DSPC-based membranes after 2 *µ*s containing ALC-0315 (*top row*) or SM-102 (*bottom row*) at cholesterol fractions of 20 mol% (**a**, **c**) and 40 mol% (**b**, **d**). DSPC lipids are shown in *orange* and cholesterol in *green*. At low pH, protonated aminolipids (*blue*) are homogeneously mixed with other membrane components, whereas at higher pH deprotonated aminolipids (*yellow*) undergo pronounced lateral segregation. Figures were rendered with PyMOL [36].

Phase separation was accompanied by pronounced changes in membrane structure. Average bilayer thickness profiles (Fig. 8) first confirm that DOPC-containing membranes (Fig. 8a,b) are generally thinner than their DSPC counterparts (Fig. 8c,d), reflecting the presence of di-saturated acyl chains. More importantly, distinct trends emerged between acidic and basic pH regimes. At low pH (*<* 6), bilayer thickness followed the order DODAP (thickest) *>* KC2/MC3 *>* SM-102 *>* ALC-0315 (thinnest), whereas at high pH the ranking shifted to KC2 (thickest) *>* MC3 *>* DODAP *>* ALC-0315/SM-102 (thinnest). These changes reflect pH-dependent aminolipid relocation within the membrane, which directly modulates bilayer thickness.

**Fig. 8.**
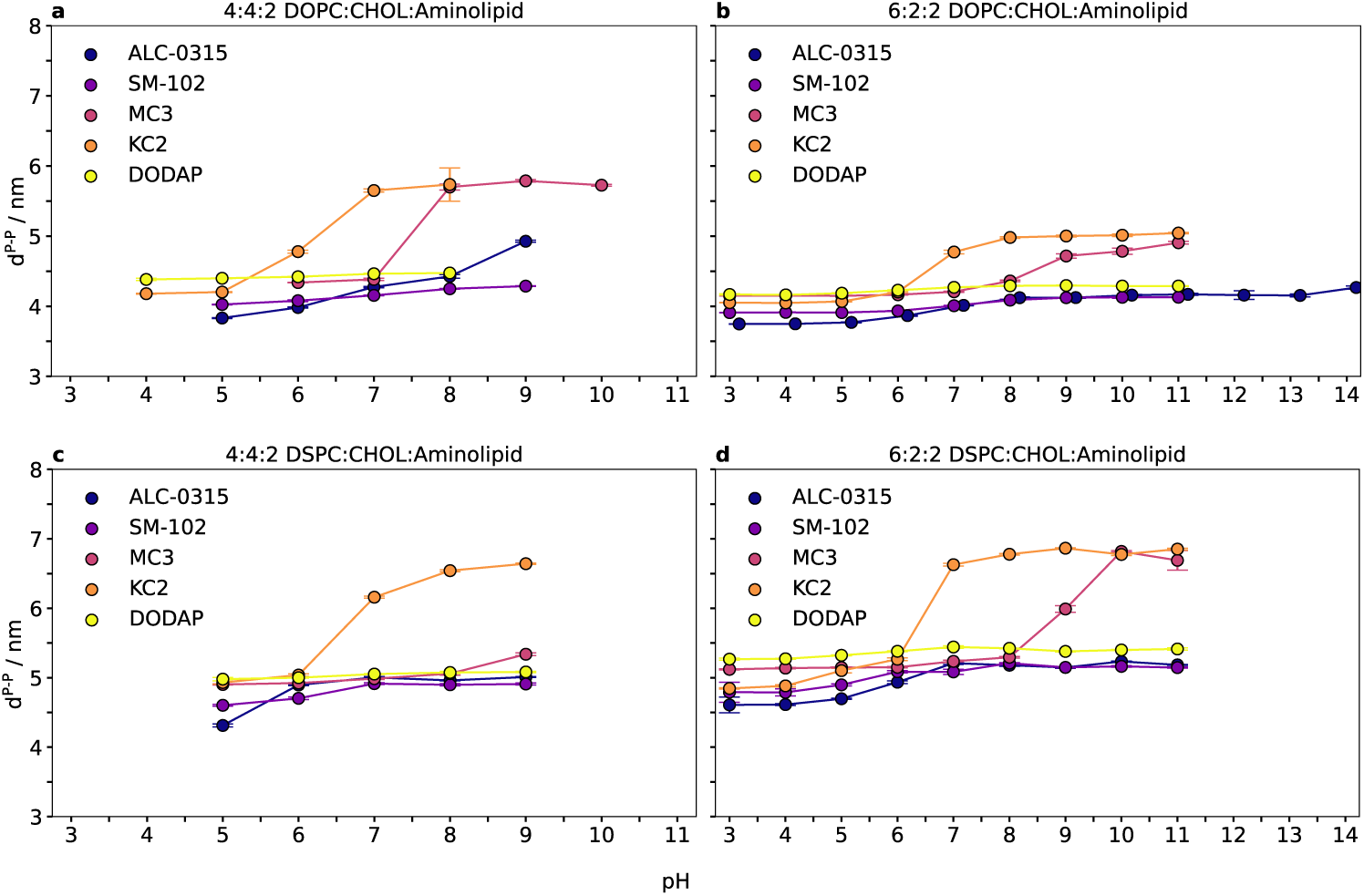
pH-dependent membrane thickness of aminolipid-containing membranes. Membrane thickness as a function of pH for systems containing **a**, **b** DOPC or **c**, **d** DSPC. Thickness was defined as the distance between the average z-positions of the phosphorus atoms in the upper and lower membrane leaflets. Average z-positions were computed using a periodic trigonometric approach (see Methods). Error bars indicate the standard error of the mean, obtained by block averaging (see Methods). The values for the thickness as a function of time are shown in Supplementary Figures 67 – 70.

Notably, KC2- and MC3-containing membranes thickened by approximately 1.5 nm upon aminolipid migration toward the membrane core (Fig. 8a). This transition occurred at lower pH for KC2 than for MC3 and was observed across all tested compositions, except for the 4:4:2 DSPC:cholesterol:MC3 system, where membrane thickening was minimal. In contrast, systems containing ALC-0315 or SM-102 exhibited only minor thickness changes, while DODAP-containing membranes remained essentially unaffected across the entire pH range.

Variations in pH also reshaped the hydrogen-bonding networks of aminolipids at the membrane interface. At low pH, aminolipids formed numerous hydrogen bonds with surrounding phospholipids (Fig. 9), driven by electrostatic attraction between protonated aminolipid headgroups and anionic phosphate groups of the helper lipids. Upon deprotonation at higher pH, these interactions were either redirected toward cholesterol and neighboring aminolipids (ALC-0315, SM-102, DODAP) or almost completely lost (KC2, MC3).

**Fig. 9.**
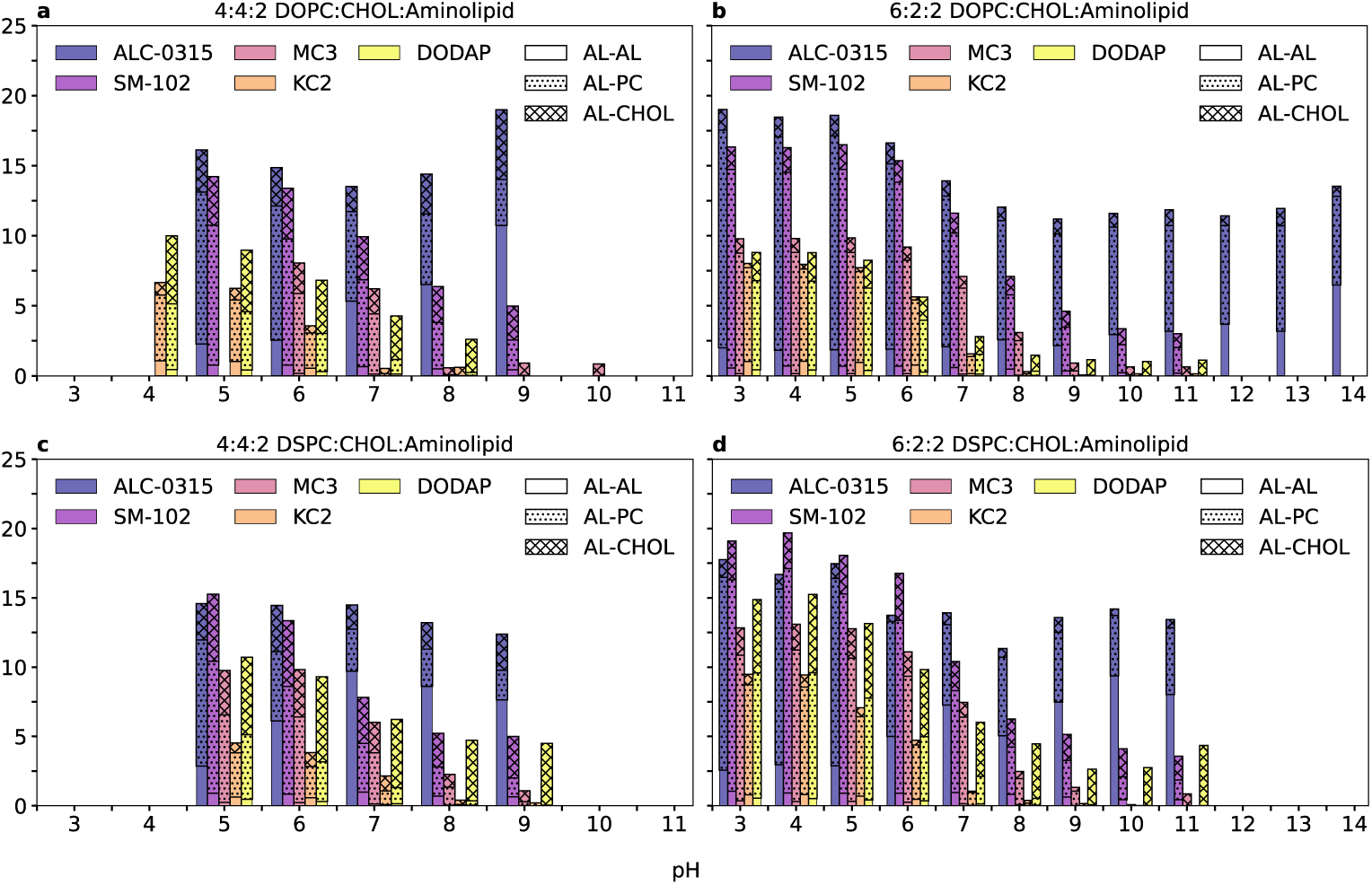
pH-dependent hydrogen bonding of aminolipids. Average number of hydrogen bonds between aminolipids and the surrounding membrane components as a function of pH for systems containing **a**, **b** DOPC or **c**, **d** DSPC lipids. Hydrogen bonds were identified using the MDAnalysis library (v2.7.0) with a distance cutoff of 3.0 Å and an angle cutoff of 150°.

Branched aminolipids ALC-0315 and SM-102 exhibited enhanced hydrogen-bonding capacity due to their two ester linkages and hydroxyl-containing headgroups. For ALC-0315, the total number of intermolecular hydrogen bonds remained nearly constant across the studied pH range; however, the fraction of ALC-0315–ALC-0315 contacts increased, consistent with its pronounced lateral segregation. DODAP maintained hydrogen bonding with cholesterol via its ester moieties even at elevated pH. In contrast, KC2 and MC3 formed hydrogen bonds with DOPC or DSPC only at low pH and lost nearly all such interactions upon migration into the membrane core. Overall, KC2 exhibited the lowest hydrogen-bonding propensity, consistent with the low polarity of its 1,3-dioxolane headgroup (Fig. 1).

Increasing pH further induced dehydration of the aminolipid titratable sites, quantified as the average number of water molecules within 0.5 nm of the N–H proton (Fig. 10). MC3, KC2, and DODAP displayed high hydration at low pH, consistent with solvent-exposed titratable sites. Upon inward migration of KC2 and MC3, their hydration shells were almost completely lost, whereas DODAP remained at the membrane surface and exhibited only a modest reduction in solvation. In contrast, the titratable sites of branched aminolipids (ALC-0315 and SM-102) were less solvent-exposed even at low pH, resulting in less pronounced dehydration at higher pH. Nonetheless, a clear dehydration trend was observed at neutral and basic pH, indicating a condensing effect associated with lateral segregation of these aminolipids.

**Fig. 10.**
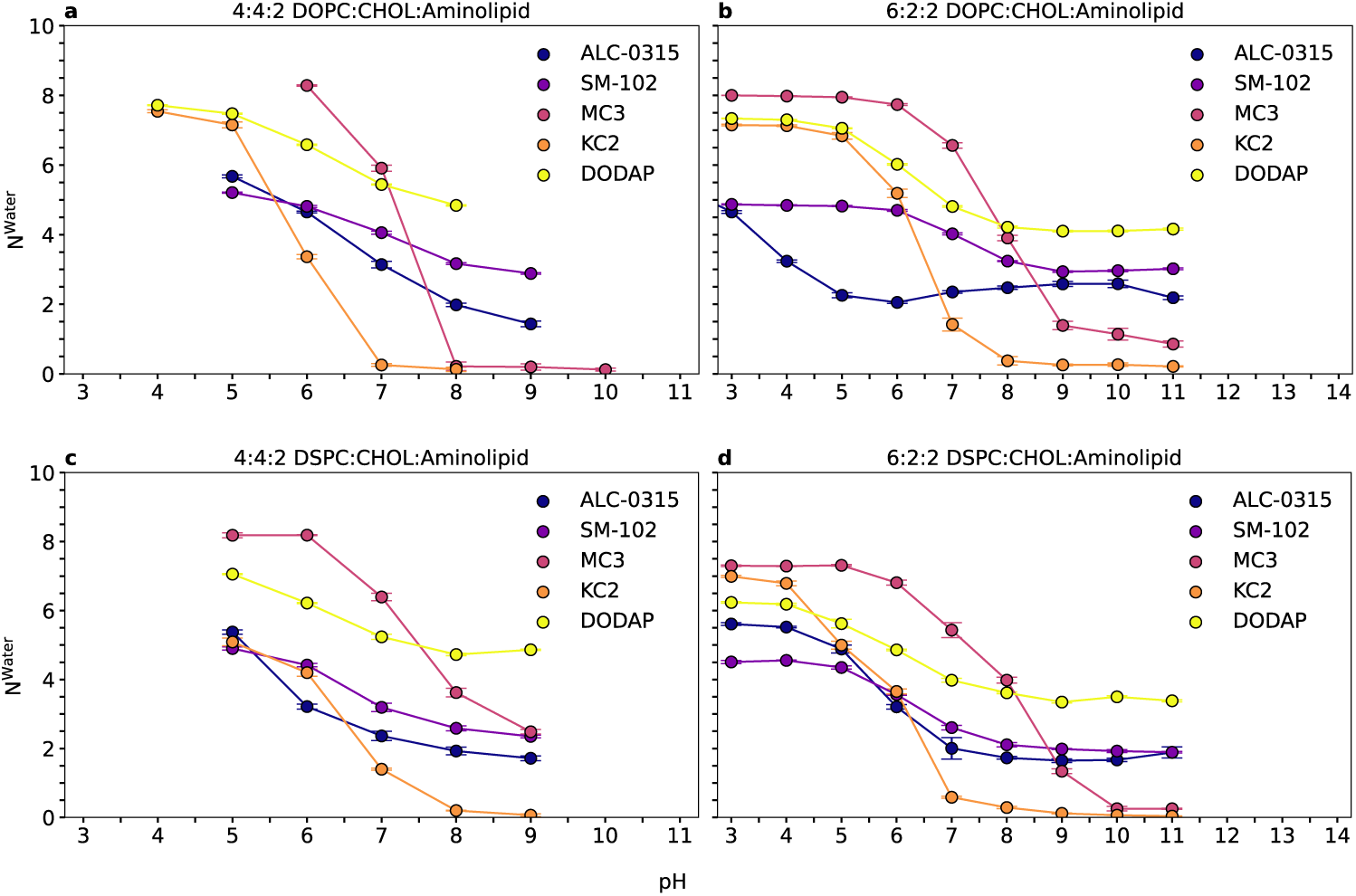
pH-dependent hydration of aminolipid titratable sites. Average number of water molecules within 0.5 nm of the aminolipid titratable site as a function of pH for systems containing **a**, **b** DOPC or **c**, **d** DSPC phospholipids. The values for the number of water molecules as a function of time are shown in Supplementary Figures 71 – 74.

## Discussion

CpHMD simulations of aminolipid–membrane systems enabled an unbiased investigation of pH-driven membrane remodeling associated with aminolipid (de)protonation and revealed a nonlinear coupling between membrane composition, membrane organization, and aminolipid protonation equilibria. Importantly, and in contrast to previous work [37], we demonstrate that CpHMD simulations of lipid membranes yield well-defined, unbiased titration curves.

Despite substantial differences among the investigated aminolipids in headgroup chemistry, acyl-chain number, saturation, and intrinsic pK_a_ values ranging from 7.6 to 9.4, the apparent aminolipid pK_a_ values converged to a narrow, physiologically relevant range (6 – 7.5). These values are only slightly higher than pK_a_ estimates reported for full lipid nanoparticles based on TNS binding assays or *ζ*-potential measurements [21, 38–40]. This agreement is notable given that the ternary membranes studied here differ substantially from typical LNP formulations (50:38.5:10:1.5 mol% aminolipid:cholesterol:helper lipid:PEG-lipid [21, 41]). Comparison with previous *in silico* studies [2, 18] and related experimental work [42, 43] suggests that the present model membranes more closely mimic the surface-exposed lipid environment of LNPs than their bulk composition, likely explaining the close correspondence between apparent aminolipid pK_a_ values and experimentally determined LNP pK_a_ values. A detailed discussion of the relationship between surface protonation and LNP pK_a_ is provided elsewhere [18].

For the polyunsaturated aminolipids DLin-KC2-DMA and DLin-MC3-DMA, increasing pH induces aminolipid deprotonation and drives migration into the hydrophobic membrane core, as reflected by a pronounced decrease in mixing entropy (Fig. 4) and their density along the z-axis (Supplementary Figures 45 a–d & 47 a–d). Despite similar intrinsic pK_a_ values and closely related molecular architectures, DLin-KC2-DMA consistently exhibits a lower apparent pK_a_ than DLin-MC3-DMA across all membrane compositions (Fig. 3a,b). This difference can be attributed to the presence of a polar ester group in the DLin-MC3-DMA headgroup, compared to the apolar 1,3-dioxolane moiety in DLin-KC2-DMA. Consequently, DLin-MC3-DMA retains a denser hydration shell at low pH (Fig. 10) and participates more extensively in intermolecular hydrogen bonding (Fig. 9), stabilizing its protonated state by approximately 7.5 – 9.2 kJ mol*^−^*^1^ relative to DLin-KC2-DMA.

The smallest pK_a_ shift was observed for DODAP. This aminolipid is anchored at the membrane interface by its highly polar, strongly hydrated headgroup (Fig. 10) and two ester linkages connecting the headgroup to the oleyl tails. DODAP preferentially forms hydrogen bonds with both cholesterol and phospholipids (Fig. 9), which stabilizes its interfacial localization (Supplementary Figures 45 e–f & 47 e–f) and limits the pK_a_ shifts with respect to the intrinsic pK_a_.

In contrast to the polyunsaturated, two-tailed aminolipids, the branched aminolipids SM-102 and ALC-0315 primarily undergo lateral, rather than transmembrane, segregation at pH ≥ 7 (Supplementary Figures 46 & 48). For ALC-0315, this behavior leads to enhanced aminolipid–aminolipid hydrogen bonding (Fig. 9). The observed lateral segregation arises from acyl-chain mismatch: the branched tails of ALC-0315 promote a conical molecular shape, in contrast to the more cylindrical geometry of DOPC and DSPC, favoring phospholipid self-packing and driving aminolipid clustering. SM-102 exhibits weaker segregation at basic pH (Fig. 4), likely because its non-branched, fully saturated acyl chain enables more favorable packing with both phospholipids and cholesterol. At aminolipid concentrations typical of LNPs (40 mol%), lateral segregation is concomitant with migration into the membrane core, as previously demonstrated for ALC-0315 [18]. At acidic pH, long-range electrostatic repulsion between protonated aminolipid headgroups counteracts segregation; upon deprotonation, this repulsion is lost and acyl-chain mismatch becomes dominant. For both branched aminolipids, lateral clustering is amplified in DSPC-containing membranes relative to DOPC, resulting in significantly larger pK_a_ shifts (Fig. 3d).

Although the simplified membrane compositions employed here do not fully recapitulate realistic LNP formulations, meaningful qualitative comparisons with experimental data can be drawn. Small-angle X-ray scattering measurements by He *et al.* [40] on LNPs formulated at 50:10:38.5:1.5 mol% aminolipid:DSPC:cholesterol:DMG-PEG2000 reported headgroup *d*-spacings for DODMA (a DODAP analog), MC3, ALC-0315, and SM-102 that agree well with both the absolute values and relative trends in bilayer thickness observed here (Fig. 8). Lipoplexes containing DOPC and DODAP or MC3 show slightly larger *d*-spacings and increased thickness at higher pH [44], consistent with our observations.

Cryo-TEM studies consistently report electron-dense, internally amorphous morphologies for LNPs containing DLin-KC2-DMA [21, 45, 46] and DLin-MC3-DMA [21, 42, 47, 48], in agreement with the rapid and complete translocation of these aminolipids into the membrane core observed in our simulations. In contrast, Cryo-TEM and MD studies report homogeneous, electron-dense interiors for ALC-0315 [2, 24, 35, 38, 39] and SM-102 [35, 47, 48]. Two factors may explain the absence of core migration for branched aminolipids in the present simulations: (i) the simulated timescale may be insufficient to capture this transition, or (ii) higher aminolipid concentrations may be required. The latter interpretation is consistent with the findings of Dehghani-Ghahnaviyeh *et al.* [49], who observed pronounced membrane reorganization only at aminolipid concentrations exceeding 20 mol%. This aligns with our observations, as Lipid-5, that is structurally similar to SM-102, was simulated as a racemic mixture of protonated and deprotonated species corresponding to pH ≈ 7.

## Conclusion

Using all-atom constant-pH molecular dynamics (CpHMD) simulations, this study elucidates how aminolipid protonation equilibria and membrane reorganization are jointly governed by aminolipid structure and lipid membrane composition. CpHMD parameters for each aminolipid were derived using thermodynamic integration [50] and subsequently applied to simplified ternary membrane systems, enabling an unbiased description of pH-dependent protonation and membrane remodeling. Although these model membranes do not aim to reproduce the full structural complexity of LNPs, they provide controlled access to the local interfacial lipid environment that governs aminolipid protonation behavior. At the same time, this approach avoids artificial constraints on lateral degrees of freedom (e.g., fixed monolayer area), which can suppress intrinsic pH-driven reorganization and potentially lead to non-physical charge distributions in LNP-mimetic systems [51].

Across all investigated compositions, the apparent – or functional – aminolipid pK_a_ values were lower compared to the intrinsic values at infinite dilution, in agreement with experimental observations on lipid nanoparticles.

Distinct reorganization mechanisms were observed depending on aminolipid architecture. The polyunsaturated aminolipids DLin-KC2-DMA and DLin-MC3-DMA rapidly and fully segregated from helper lipids upon deprotonation, migrating towards the hydrophobic membrane core. This behavior is driven by the fluidity of the diunsaturated acyl chains and a limited capacity to form stabilizing hydrogen bonds at the membrane interface. In contrast, DODAP populated the membrane surface across the full pH range, owing to its more polar headgroup conferring stronger retention of the hydration shell, finally resulting in the lowest observed ΔpK_a_.

In contrast, the branched aminolipids ALC-0315 and SM-102 undergo pronounced lateral segregation, intermediate headgroup dehydration, and exhibit ΔpK_a_ values that are highly sensitive to helper lipid type. Upon deprotonation, these aminolipids remain surface-localized and form in-plane clusters rather than migrating into the membrane core, as observed for polyunsaturated aminolipids. Likely, this distinct reorganization pathway directly shifts the surface-to-core distribution of aminolipids within LNPs. Enrichment of branched aminolipids with inverted cone-like molecular geometry at the particle surface increases interfacial curvature stress and thereby promotes larger LNP radii.

Collectively, our results demonstrate that aminolipid phase separation — whether vertical (surface-to-core migration) or lateral (in-plane clustering) — destabilizes the protonated state and promotes a larger shift between intrinsic and apparent pK_a_ values. The extent and nature of this shift depend critically on both membrane composition and aminolipid molecular geometry. Saturated helper lipids (DSPC) consistently enhanced segregation relative to unsaturated DOPC, amplifying pK_a_ shifts across all aminolipid classes. These findings establish a direct mechanistic link between membrane-driven lipid organization and aminolipid protonation equilibria, providing design principles for tuning LNP composition toward optimized pH responsiveness, membrane remodeling, and intracellular delivery efficiency.

## Methods

### System preparation

Systems containing ALC-0315 and SM-102 were assembled in a bilayer configuration together with the corresponding helper lipid and cholesterol using the MemPackGen tool [52] from the AmberTools suite [53]. Systems containing MC3, KC2, and DODAP were prepared using CHARMM-GUI [54], employing the protonated form of the respective aminolipid in combination with the helper lipid and cholesterol.

In all cases, the membrane systems were solvated with approximately 12,000 TIPS3P [55] water molecules, corresponding to a hydration level of roughly 50 water molecules per lipid, and neutralized by the addition of chloride counterions.

### Force field parameters

All molecular mechanics parameters for the aminolipids were derived using the CHARMM General Force Field (CGenFF) [56] in combination with the CHARMM36 lipid force field [57]. Parameters for the protonated and deprotonated states of MC3, KC2, and DODAP were adopted from the work of Park *et al.* [58]. Force-field parameters for ALC-0315 were taken from Trollmann and Böckmann [18]. SM-102 was parameterized following the same workflow previously established for ALC-0315 [18].

An initial topology for SM-102 was generated using the CGenFF program (v4.0) [59–61], compatible with CGenFF v4.6. Partial atomic charges were assigned based on standard CHARMM chemical moieties [56, 57]. All bonded parameters for the protonated form of SM-102 exhibited penalty scores below 1.5 and were therefore accepted without further refinement.

Parameters for cholesterol, DSPC, and DOPC were taken from the CHARMM36 lipid force field [57, 62, 63]. All simulations employed the July 2022 release of the CHARMM36 force-field port for GROMACS.

### General MD parameters

All MD simulations were performed using the classical leap-frog integrator with a time step of 2 fs. The minimum update frequency of the neighbor list was set to 20 steps using the Verlet cutoff scheme, with the verlet-buffer-tolerance set to the default value of 0.005 kJ mol*^−^*^1^ ps*^−^*^1^. Consequently, the short-range neighbor-list cutoff (rlist) was automatically adjusted during the simulation.

Electrostatic interactions were computed using the smooth Particle-Mesh Ewald (PME) method [64], with a real-space Coulomb cutoff of 1.2 nm and a Fourier grid spacing of 0.14 nm. Van der Waals interactions were evaluated with a cutoff of 1.2 nm and smoothly switched off between 1.0 and 1.2 nm using the force-switch modifier.

All simulations were conducted at a reference temperature of 310 K, maintained using the velocity-rescaling (v-rescale) thermostat with a coupling time constant of 1 ps. Two independent temperature-coupling groups were employed: one for all membrane components (lipids) and one for the solvent components (water, counterions, and buffer particles). Pressure was maintained at 1 bar using the c-rescale barostat with semi-isotropic pressure coupling, a coupling time constant of 5 ps, and the standard isothermal compressibility of water (4.5 × 10*^−^*^5^ bar*^−^*^1^).

All bonds involving hydrogen atoms were constrained using the LINCS algorithm, employing default settings (fourth-order expansion and one iteration).

### Constant-pH parameters

All constant-pH molecular dynamics (CpHMD) simulations were performed using the scalable *λ*-dynamics implementation implemented by Aho *et al.* [23] within the GROMACS MD engine [65, 66]. For each aminolipid, a single titratable site was defined, comprising a subset of atoms whose partial charges were scaled according to the instantaneous protonation state. An overview of the selected titratable atoms and their partial charges is provided in Fig. 11.

**Fig. 11.**
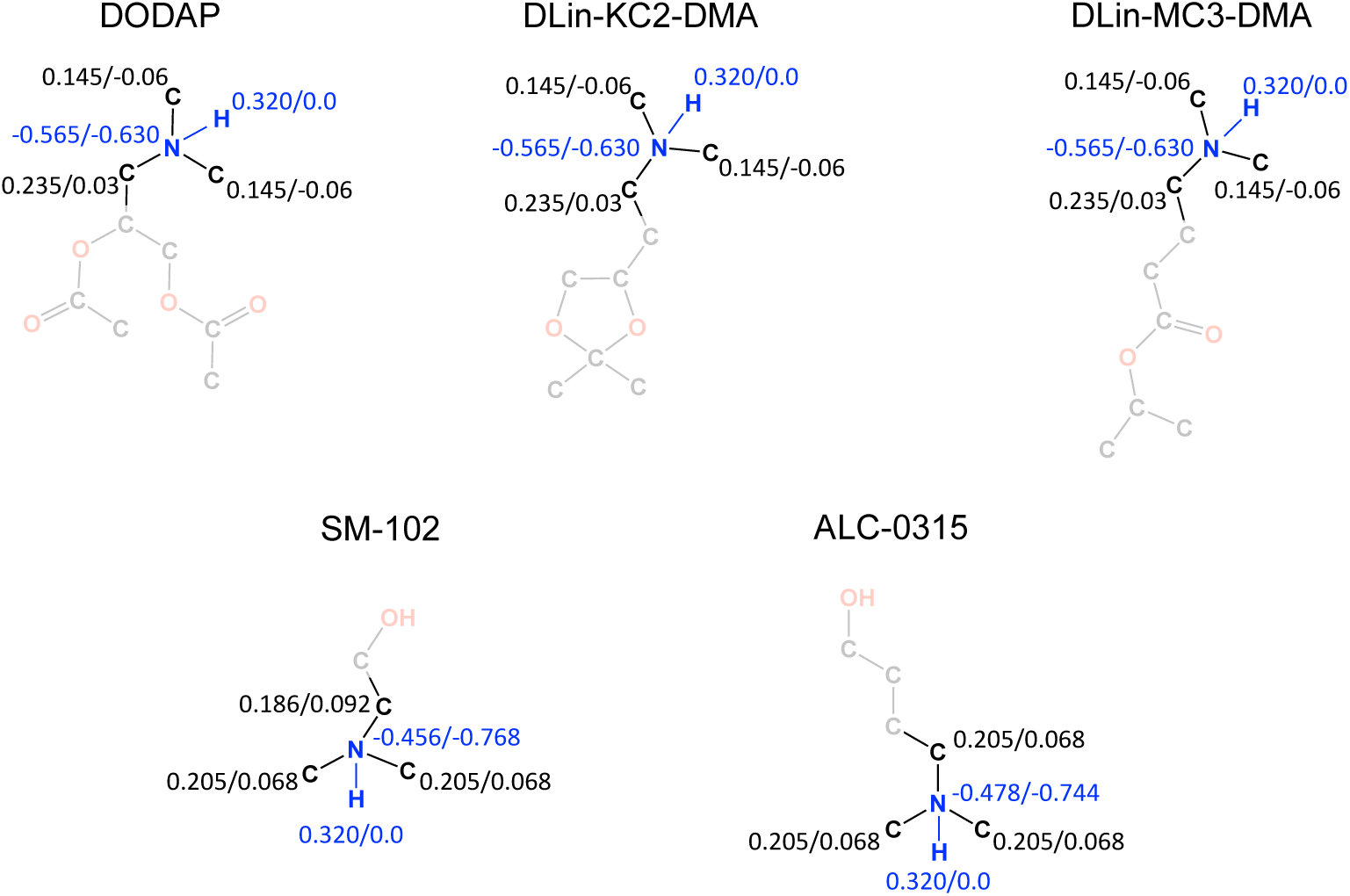
Titratable sites of the studied aminolipids. Headgroup structures of each aminolipid with the corresponding partial charges in the protonated and deprotonated states. Carbons are shown in black, oxygens in red, nitrogens in blue, and polar hydrogens in blue/red. Aliphatic hydrogens are omitted for clarity. Transparent atoms indicate non-titratable groups whose partial charges remain fixed.

In the current CpHMD implementation, *λ*-dynamics is used to interpolate between the protonated (*λ_i_* = 0) and deprotonated (*λ_i_* = 1) states of titratable site *i* by scaling the atomic partial charges. The resulting *λ*-dependent electrostatic potential Φ(**R***_i_, λ*) acting on atom *i* is given by

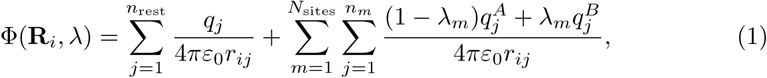

where *n*_rest_ denotes the number of atoms not belonging to any titratable site, *N*_sites_ is the total number of titratable sites, *n_m_* is the number of atoms in titratable site *m*, and *q_j_^A^* and *q_j_^B^* are the partial charges of atom *j* in the protonated and deprotonated states, respectively. Through this coupling, the protonation state of each amino-lipid becomes dependent on its local environment, with the *λ*-coordinate evolving dynamically according to Newton’s equations of motion.

Accurate CpHMD simulations require estimation of the protonation free-energy term *V* ^MM^(*λ*) for each titratable site [23, 50, 67]. Parameterization followed the workflow established in our previous work [18, 24]. Briefly, a single aminolipid and one buffer particle were placed in a 6 × 6 × 6 nm^3^ simulation box containing water and 150 mM Na^+^Cl^-^. Thermodynamic integration (TI) was then performed to estimate *∂V* ^MM^*/∂λ* at fixed values of the *λ*-coordinate. For each aminolipid, 25 independent TI simulations were carried out with *λ* values spanning the interval [−0.10, 1.10] in increments of 0.05. Simulation lengths depended on aminolipid size, with larger aminolipids (ALC-0315 and SM-102) requiring longer trajectories to achieve convergence. A detailed overview of TI window lengths is provided in Tab. 4.

**Table 4.**
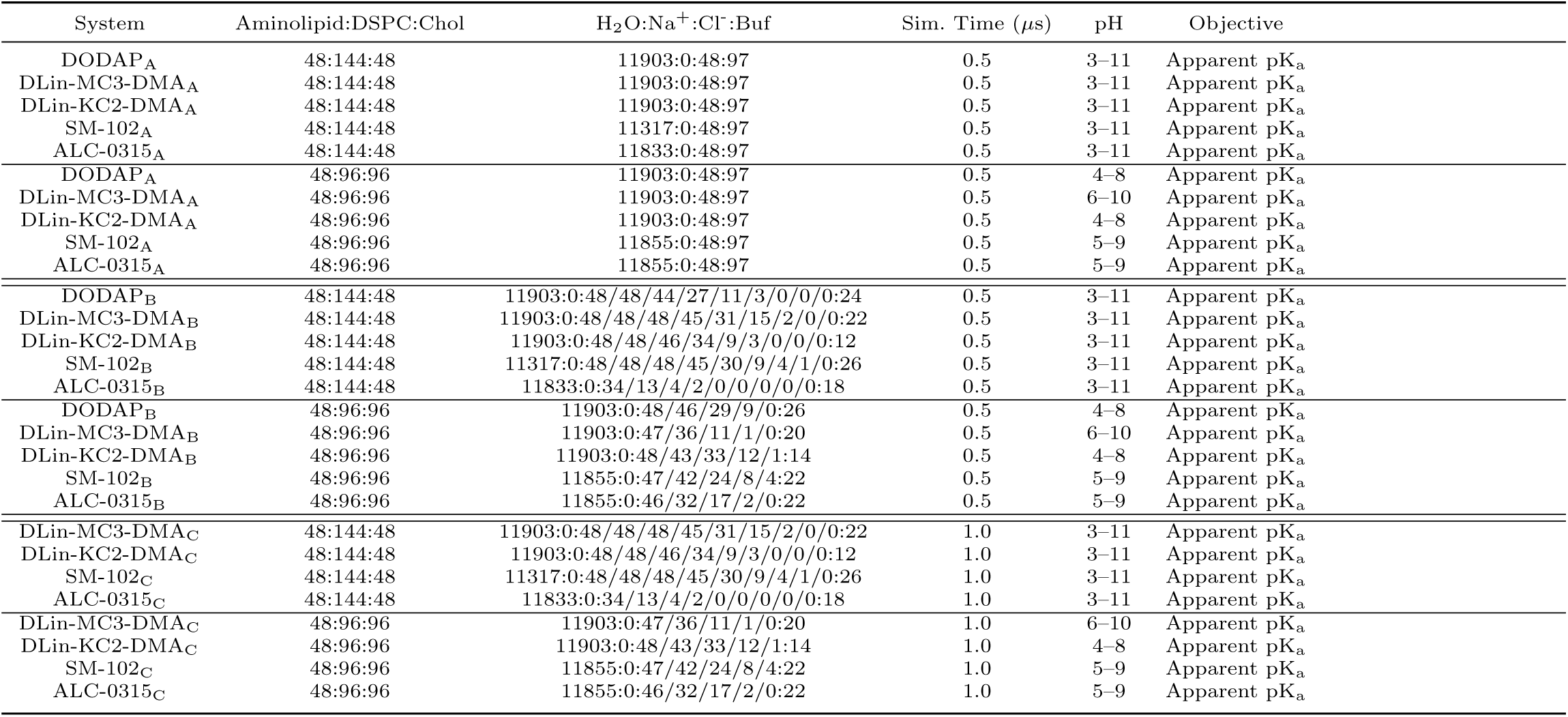
All-atom molecular dynamics simulations of ternary membranes including the helperlipid DSPC using the constant-pH method.

Following the recommendations of Buslaev *et al.* [50], the resulting *∂V* ^MM^*/∂λ* profiles were fitted using high-order polynomials. A ninth-order polynomial was employed for ALC-0315 and SM-102, while an eighth-order polynomial was sufficient for the remaining aminolipids. In addition to *V* ^MM^, two auxiliary potentials act on the *λ*-coordinates: *V* ^pH^, which accounts for the effect of the environmental pH, and *V* ^bias^, which prevents divergence of *λ* toward unphysical values and stabilizes sampling near the physical end states (*λ* ≈ 0 and *λ* ≈ 1). For all simulations, the barrier height of *V* ^bias^ was set to 7.5 kJ mol*^−^*^1^.

### Adjustment of buffer particles

The initial number of buffer particles was set to 97, corresponding to the maximum number of buffer particles allowed in the simulations (2*N* + 1, where *N* is the number of titratable sites). This choice ensures charge compensation for the largest possible fluctuation in membrane charge, corresponding to complete deprotonation of all aminolipids in the system. Although buffer particles are designed to minimize interference with the system [50], their number should be reduced to the required minimum.

After 500 ns of simulation, the dominant changes in membrane protonation had occurred, resulting in a substantial redistribution of charge to the buffer particles. At this stage, the simulations were restarted to reduce the number of buffer particles and to reset their partial charges to zero (i.e., *λ*^Buffer^ = 0.5). The required number of buffer particles was determined based on the charge fluctuations and drift observed during the preceding trajectory segment. The number of counterions was adjusted accordingly to maintain overall charge neutrality. Buffer particles located closest to the membrane and counterions farthest from the membrane were preferentially removed. Protonation states of all titratable sites in the final frame of the previous run were used to initialize the restarted simulations.

To ensure numerical stability after particle removal, a short energy minimization of up to 30 iterations was performed prior to resuming the simulations. This procedure was repeated once more after an additional 500 ns of simulation time.

### Validation

For each aminolipid, the fitted polynomial coefficients were validated using two complementary approaches. (i) A CpHMD simulation of a single aminolipid in solution was performed without the *V* ^pH^ contribution and with the barrier height set to 0.0 kJ mol*^−^*^1^. Under these conditions, the *λ*-coordinate is expected to sample the interval [0, 1] approximately uniformly. (ii) Titration simulations at infinite dilution were carried out, in which a single aminolipid was simulated over a series of pH values controlled by *V* ^pH^, using a barrier height of 7.5 kJ mol*^−^*^1^. From the fraction of simulation frames in which the aminolipid was deprotonated at each pH value, the intrinsic pK_a_ was determined. Agreement between the numerically obtained intrinsic pK_a_ and the value specified in the simulation input (.mdp file) confirmed that the fitted polynomial coefficients accurately reproduce the protonation free-energy profile.

### Equilibration and error estimation

Estimating observables from molecular dynamics trajectories as time averages without significant drift is a common challenge. Given the large number of simulations performed in this study, an automated and robust procedure was implemented to assess equilibration and convergence for any averaged observable of interest (e.g., *S^deprot^*, *S^mix^*, Thickness,. . .).

To quantify potential drift, a linear model was first fitted to the observable over the full trajectory interval [0*, T*]. Subsequently, progressively larger initial portions of the trajectory were discarded, and the linear fit was recomputed for each truncated interval [*τ_i_, T*]:

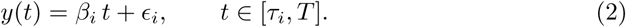

where *β_i_* and *ɛ_i_* denote the slope and intercept of the linear regression, respectively. For each truncated interval, the drift was defined as the difference between the fitted model values at the endpoints:

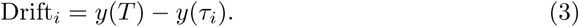

To obtain a dimensionless and objective measure of drift, this quantity was normalized by the empirical standard deviation *σ_i_* of the observable within the same interval:

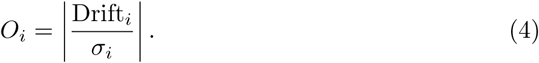

An observable was considered equilibrated once *O_i_ <* 1 for the first time, indicating that the residual drift was smaller than the intrinsic fluctuations of the signal. Standard errors of the mean were then estimated from the equilibrated portion of the trajectory using block averaging. The resulting error estimates were fitted to either a single- or double-exponential model following the approach of Hess [68]. In cases where the automated workflow yielded insufficient equilibration times, the starting points were adjusted manually. The final assigned equilibration times are shown as vertical lines in Supplementary Figures 49 – 56 and 61 – 78.

### Protonation state analysis

Protonation states of individual aminolipids were assigned following the recommendations of Aho *et al.* [23]. For lipid *i* in simulation frame *j*, the lipid was classified as protonated if *λ_j_^i^* < 0.2 and as deprotonated if *λ_j_^i^ >* 0.8. Frames in which *λ_j_^i^* fell within the intermediate range were excluded from protonation state analysis for that lipid. For each simulation frame, the fraction of deprotonated aminolipids was computed as

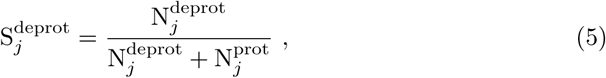

where N*_j_*^deprot^ and N*_j_*^prot^ denote the numbers of deprotonated and protonated amino-lipids, respectively. For simulations at infinite dilution containing a single aminolipid, *N* ^deprot^ and *N* ^prot^ correspond to the number of frames in which the aminolipid was classified as deprotonated or protonated, respectively.

Apparent pK_a_ values were obtained by fitting the pH-dependent mean fraction of deprotonated aminolipids to the generalized Henderson-Hasselbalch equation, *h*(pH; *n, pK*

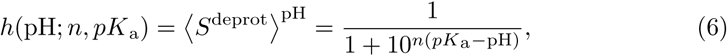

where ⟨*S*^deprot^⟩^pH^ denotes the time-averaged fraction of deprotonated aminolipids at a given pH. The parameter *n* describes the cooperativity of the transition, and pK_a_ corresponds to the pH at which ⟨*S_j_*^deprot^⟩^pH^ = 0.5.

For intrinsic pK_a_ estimation, *n* was fixed to 1.0 during least-squares fitting. For apparent pK_a_ values, an iterative reweighting least-squares scheme was employed to improve fit robustness. Initial weights were set to the inverse variance of the mean, *w*^pH^ = 1*/*(*σ*^pH^)^2^, and were iteratively updated by multiplying them with the derivative of Eq. 6,

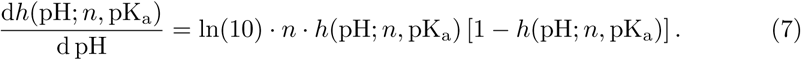

This procedure increases the relative weight of data points in the transition region, where statistical uncertainty is highest, while retaining contributions from well-sampled low- and high-pH regimes. Standard errors of the mean *σ*^pH^ were clipped to a minimum value of 10*^−^*^4^. The iterative scheme was terminated when the Euclidean norm of the difference between successive parameter vectors fell below 10*^−^*^8^.

Confidence intervals for fitted parameters were estimated using parametric bootstrapping [69, 70]. At each pH value, *S*^deprot^ was assumed to follow a normal distribution,

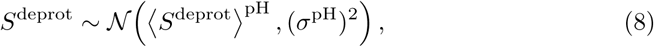

where 〈*S*^deprot^〉^pH^ denotes the mean over the equilibrated trajectory segment and *σ*^pH^ is the corresponding standard error of the mean obtained from block averaging. For each condition, 50,000 bootstrap samples were generated. Reported pK_a_ values correspond to the mean of the bootstrap distribution. The 95% confidence intervals were defined by the 2.5^th^ and 97.5^th^ percentiles of the bootstrap distribution.

### Periodic Center-of-Mass Definition

Each trajectory was post-processed by centering the membrane in the middle of the simulation box and reconstructing molecules fragmented across periodic boundary conditions (PBC). However, this workflow may occasionally result in the membrane being split across the periodic boundary in the *z*-direction. To avoid potential bias in the membrane thickness calculation and in density profiles along the *z*-axis, a periodic trigonometric approach similar to Bai *et al.* [71] was applied to determine the average position of the phosphorus atoms in the helper lipids. First, the *z*-coordinates *z_i_*were wrapped back into the primary unit cell. The coordinates were then mapped onto the unit circle according to

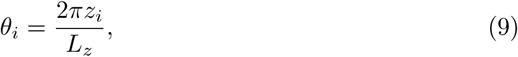

where *L_z_* denotes the box length in the *z*-direction. The averages of the sine and cosine components over all *N* phosphorus atoms were computed as

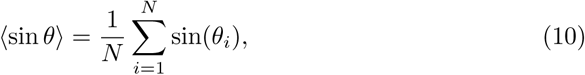

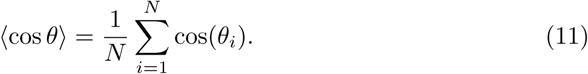

The mean *z*-position was then reconstructed using the two-argument arctangent function:

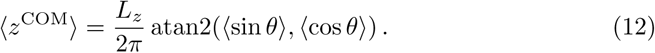

Finally, ⟨*z*^COM^⟩ was mapped back into the simulation box. At all stages, it was ensured that the average position of the upper leaflet remained greater than that of the lower leaflet. The average *z*-positions per frame for each leaflet are shown in Supplementary Figures 57 – 60.

## Data availability

All simulation models, input files and structures will be made available for download after acceptance of the manuscript.

## Acknowledgements

The authors gratefully acknowledge the scientific support and HPC resources provided by the Erlangen National High Performance Computing Center (NHR@FAU) of the Friedrich-Alexander-Universität Erlangen-Nürnberg (FAU). The hardware is funded by the German Research Foundation (DFG).

## Author contributions

M.F.W.T. and P.R. performed simulations and data analysis, R.A.B. designed the study, all wrote the manuscript.

## Additional information

The supplementary material includes

- Supplementary Figures 1 – 79

