## Supplementary Figures for "Membrane Environment Sets the Functional pK_a_ of Ionizable Lipids"

#### The PDF file includes

- Supplementary Figures 1 – 79,
- Supplementary References

### Supplementary Figures

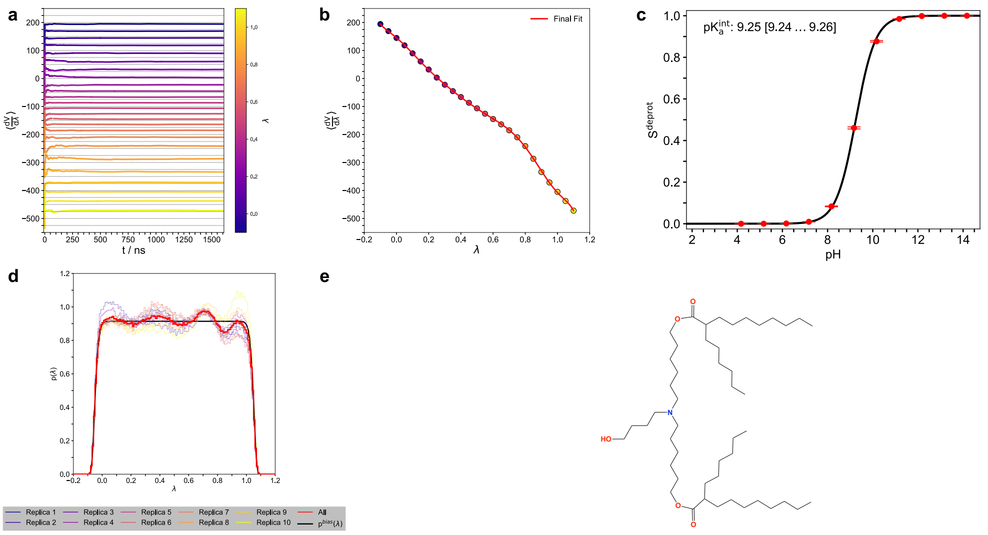


**Fig. 1** **a, b** Parameterization and **c, d** validation of polynomial coefficients used to model the correction potential for the protonation free energy of the **e** aminolipid ALC-0315. **a** Convergence of the estimated averages for $\frac{\partial V^{MM}}{\partial \lambda}$ obtained from thermodynamic integration simulations. **b** Fit of a ninth-order polynomial to the estimates of $\langle \frac{\partial V^{MM}}{\partial \lambda} \rangle$. **c** Titration curve of the aminolipid at infinite dilution, fitted with the Henderson-Hasselbalch equation. Error bars represent the standard error of the mean over ten replicas per pH level. The $\lambda$-dependent potentials $V^{bias}(\lambda)$ (barrier height 7.5 kJ·mol$^{-1}$), $V^{pH}(\lambda)$ (corresponding to the respective pH value), and the parameterized $V^{MM}(\lambda)$ were applied. **d** Distributions of the $\lambda$-coordinate obtained from ten replica simulations of the aminolipid at infinite dilution. In this case, only $V^{bias}(\lambda)$ (barrier height 0.0 kJ·mol$^{-1}$) and the parameterized $V^{MM}(\lambda)$ were applied. **e** Molecular structure of the aminolipid.

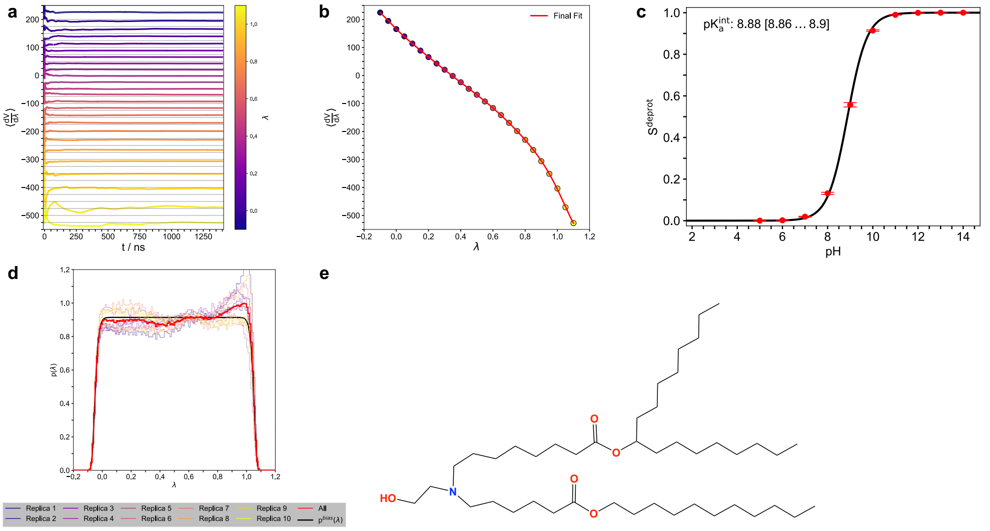


**Fig. 2** **a, b** Parameterization and **c, d** validation of polynomial coefficients used to model the correction potential for the protonation free energy of the **e** aminolipid SM-102. **a** Convergence of the estimated averages for $\frac{\partial V^{MM}}{\partial \lambda}$ obtained from thermodynamic integration simulations. **b** Fit of a ninth-order polynomial to the estimates of $\langle \frac{\partial V^{MM}}{\partial \lambda} \rangle$. **c** Titration curve of the aminolipid at infinite dilution, fitted with the Henderson-Hasselbalch equation. Error bars represent the standard error of the mean over ten replicas per pH level. The $\lambda$-dependent potentials $V^{bias}(\lambda)$ (barrier height 7.5 kJ·mol$^{-1}$), $V^{pH}(\lambda)$ (corresponding to the respective pH value), and the parameterized $V^{MM}(\lambda)$ were applied. **d** Distributions of the $\lambda$-coordinate obtained from ten replica simulations of the aminolipid at infinite dilution. In this case, only $V^{bias}(\lambda)$ (barrier height 0.0 kJ·mol$^{-1}$) and the parameterized $V^{MM}(\lambda)$ were applied. **e** Molecular structure of the aminolipid.

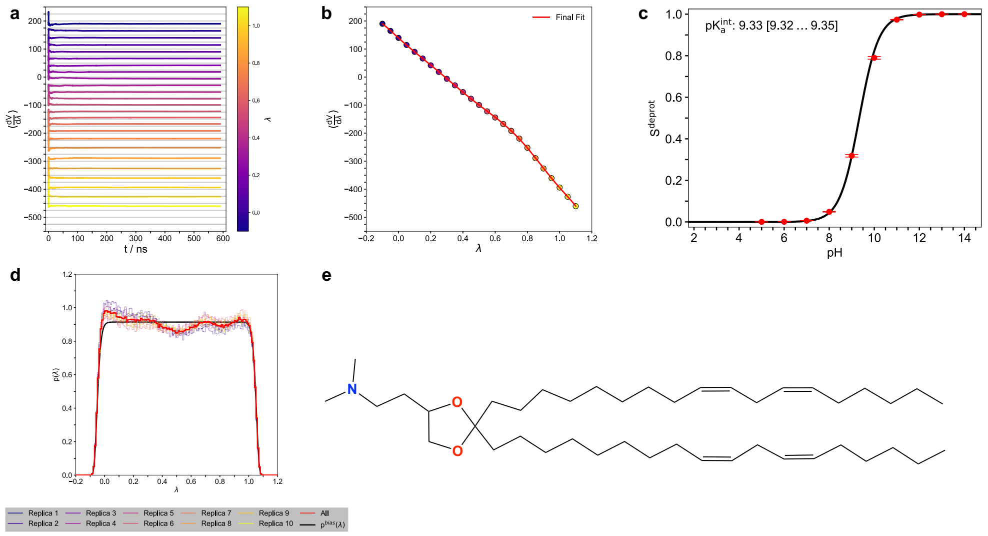


**Fig. 3** **a, b** Parameterization and **c, d** validation of polynomial coefficients used to model the correction potential for the protonation free energy of the **e** aminolipid DLin-KC2-DMA. **a** Convergence of the estimated averages for $\frac{\partial V^{MM}}{\partial \lambda}$ obtained from thermodynamic integration simulations. **b** Fit of an eighth-order polynomial to the estimates of $\langle \frac{\partial V^{MM}}{\partial \lambda} \rangle$. **c** Titration curve of the aminolipid at infinite dilution, fitted with the Henderson-Hasselbalch equation. Error bars represent the standard error of the mean over ten replicas per pH level. The $\lambda$-dependent potentials $V^{bias}(\lambda)$ (barrier height $7.5\,\mathrm{kJ{\cdot}mol^{-1}}$), $V^{pH}(\lambda)$ (corresponding to the respective pH value), and the parameterized $V^{MM}(\lambda)$ were applied. **d** Distributions of the $\lambda$-coordinate obtained from ten replica simulations of the aminolipid at infinite dilution. In this case, only $V^{bias}(\lambda)$ (barrier height $0.0\,\mathrm{kJ{\cdot}mol^{-1}}$) and the parameterized $V^{MM}(\lambda)$ were applied. **e** Molecular structure of the aminolipid.

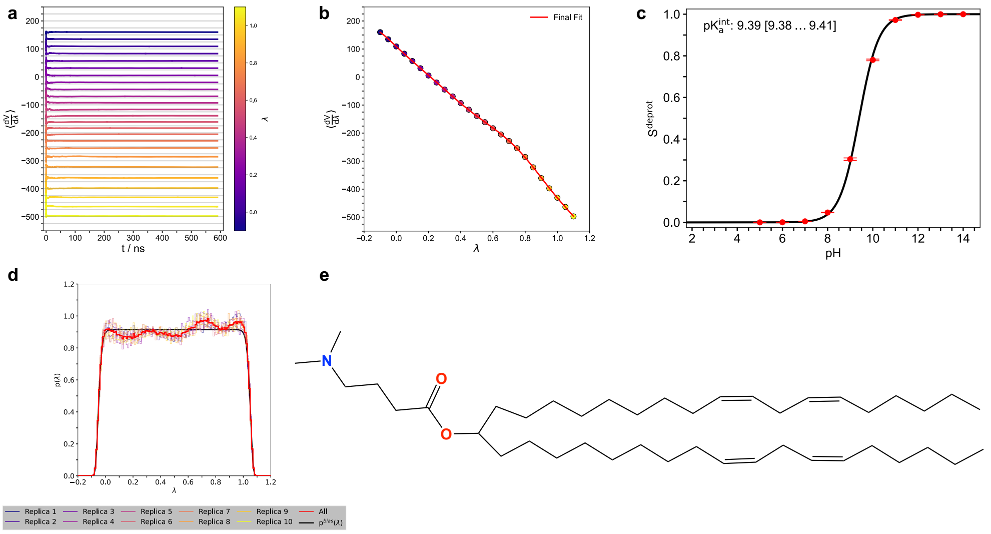


**Fig. 4** **a, b** Parameterization and **c, d** validation of polynomial coefficients used to model the correction potential for the protonation free energy of the **e** aminolipid DLin-MC3-DMA. **a** Convergence of the estimated averages for $\frac{\partial V^{MM}}{\partial \lambda}$ obtained from thermodynamic integration simulations. **b** Fit of an eighth-order polynomial to the estimates of $\langle\frac{\partial V^{MM}}{\partial \lambda}\rangle$. **c** Titration curve of the aminolipid at infinite dilution, fitted with the Henderson-Hasselbalch equation. Error bars represent the standard error of the mean over ten replicas per pH level. The $\lambda$-dependent potentials $V^{bias}(\lambda)$ (barrier height 7.5 kJ·mol$^{-1}$), $V^{pH}(\lambda)$ (corresponding to the respective pH value), and the parameterized $V^{MM}(\lambda)$ were applied. **d** Distributions of the $\lambda$-coordinate obtained from ten replica simulations of the aminolipid at infinite dilution. In this case, only $V^{bias}(\lambda)$ (barrier height 0.0 kJ·mol$^{-1}$) and the parameterized $V^{MM}(\lambda)$ were applied. **e** Molecular structure of the aminolipid.

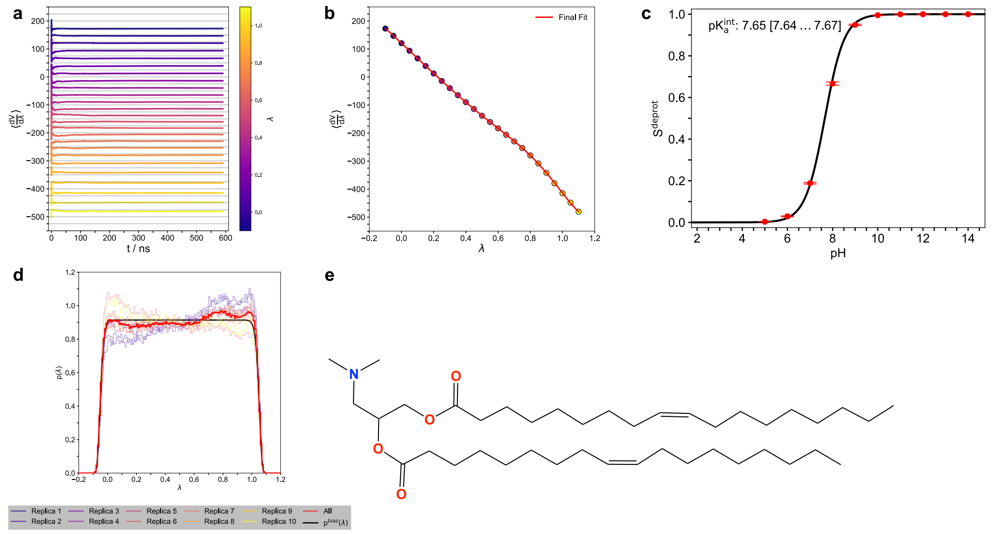


**Fig. 5** **a, b** Parameterization and **c, d** validation of polynomial coefficients used to model the correction potential for the protonation free energy of the **e** aminolipid DODAP. **a** Convergence of the estimated averages for $\frac{\partial V^{MM}}{\partial \lambda}$ obtained from thermodynamic integration simulations. **b** Fit of an eighth-order polynomial to the estimates of $\langle \frac{\partial V^{MM}}{\partial \lambda} \rangle$. **c** Titration curve of the aminolipid at infinite dilution, fitted with the Henderson-Hasselbalch equation. Error bars represent the standard error of the mean over ten replicas per pH level. The $\lambda$-dependent potentials $V^{bias}(\lambda)$ (barrier height $7.5\,\mathrm{kJ{\cdot}mol^{-1}}$), $V^{pH}(\lambda)$ (corresponding to the respective pH value), and the parameterized $V^{MM}(\lambda)$ were applied. **d** Distributions of the $\lambda$-coordinate obtained from ten replica simulations of the aminolipid at infinite dilution. In this case, only $V^{bias}(\lambda)$ (barrier height $0.0\,\mathrm{kJ{\cdot}mol^{-1}}$) and the parameterized $V^{MM}(\lambda)$ were applied. **e** Molecular structure of the aminolipid.

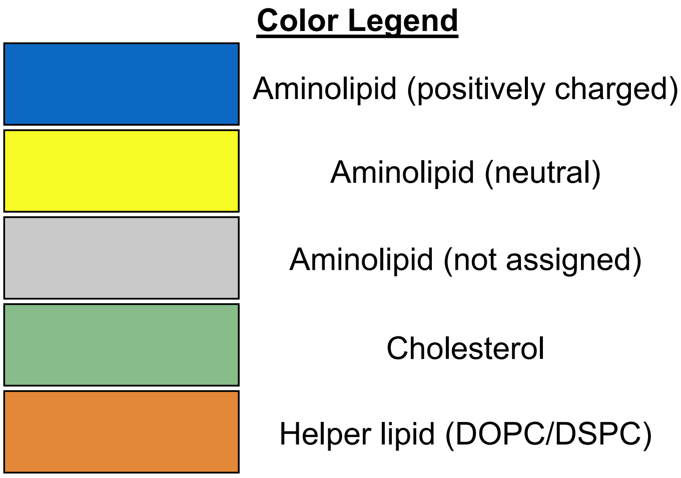


**Fig. 6** Color legend for all structural snapshots in the main text and Supporting Information. An aminolipid is classified as positively charged (blue) when its $\lambda$ coordinate is below 0.2 (protonated) and as neutral (yellow) when $\lambda$ exceeds 0.8 (deprotonated). For $0.2 \leq \lambda \leq 0.8$, the aminolipid is considered to be in a transition state (grey). Cholesterol is shown in green, and the helper lipids (DOPC/DSPC) in orange.

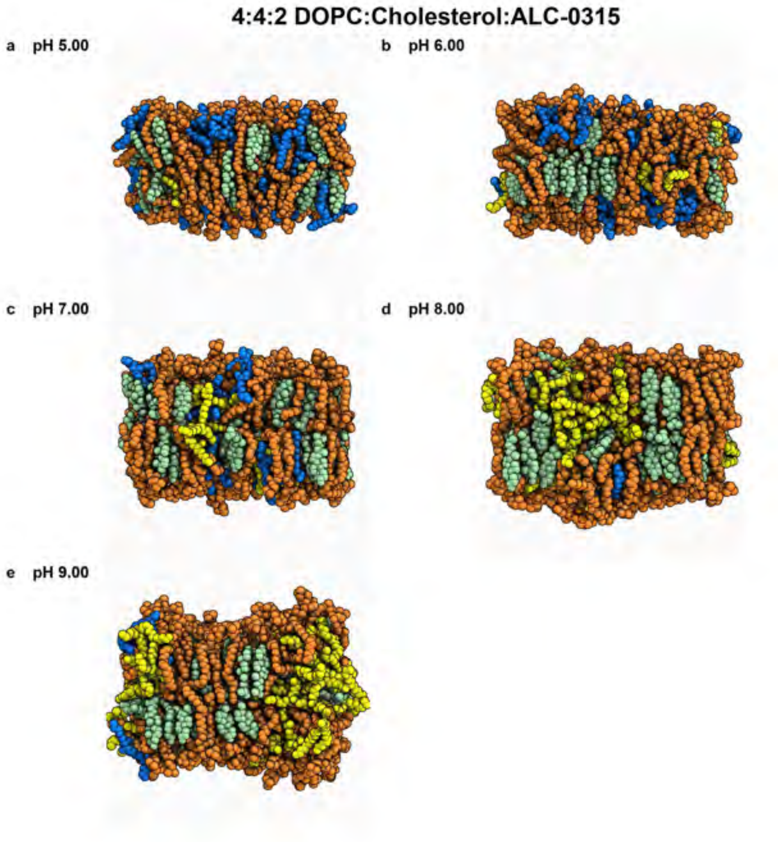


**Fig. 7** Snapshots of the membrane in the final frame of the simulation at the corresponding pH values. The helper lipid is shown in orange, cholesterol in green, the protonated aminolipid (net charge +1) in blue, and the deprotonated aminolipid (net charge 0) in yellow.

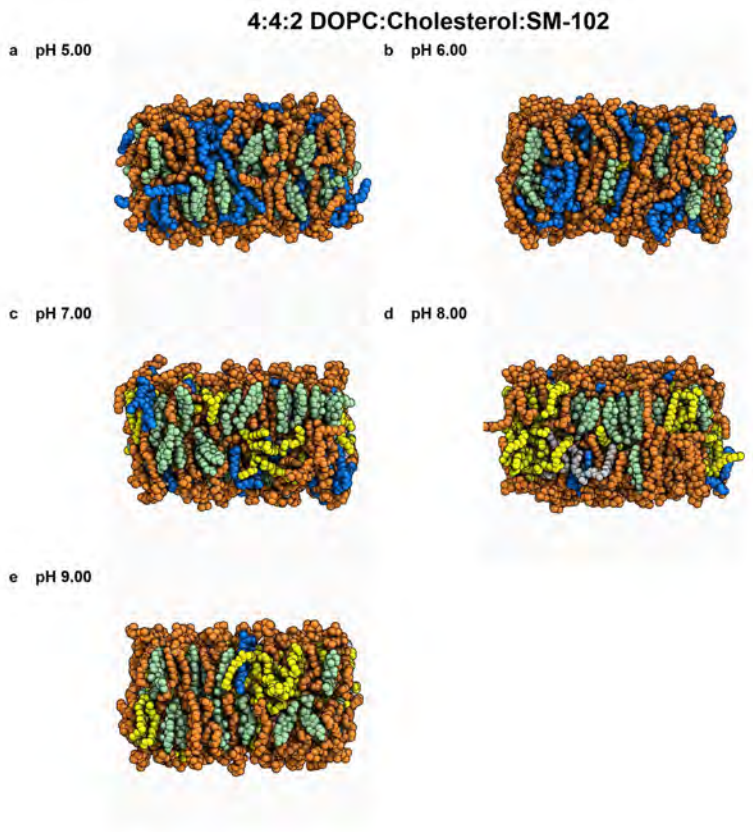


**Fig. 8** Snapshots of the membrane in the final frame of the simulation at the corresponding pH values. The helper lipid is shown in orange, cholesterol in green, the protonated aminolipid (net charge +1) in blue, and the deprotonated aminolipid (net charge 0) in yellow.

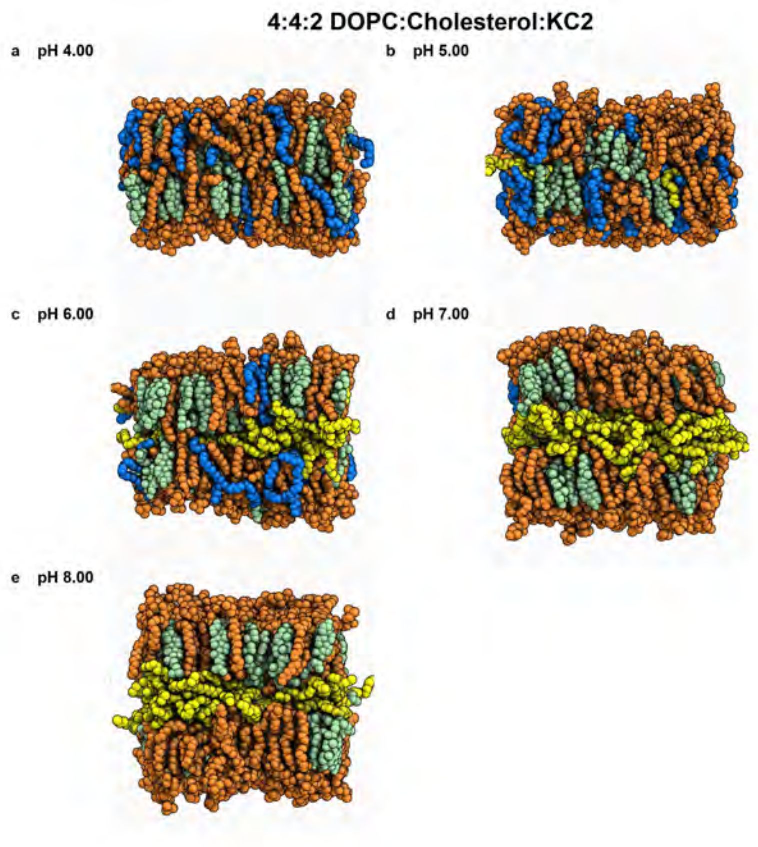


**Fig. 9** Snapshots of the membrane in the final frame of the simulation at the corresponding pH values. The helper lipid is shown in orange, cholesterol in green, the protonated aminolipid (net charge +1) in blue, and the deprotonated aminolipid (net charge 0) in yellow.

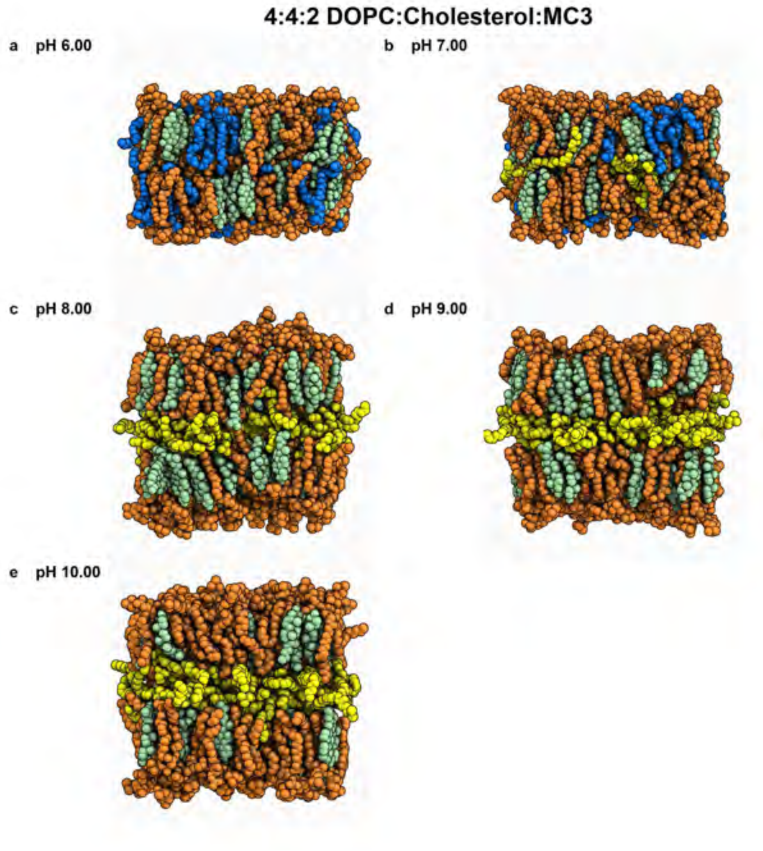


**Fig. 10** Snapshots of the membrane in the final frame of the simulation at the corresponding pH values. The helper lipid is shown in orange, cholesterol in green, the protonated aminolipid (net charge +1) in blue, and the deprotonated aminolipid (net charge 0) in yellow.

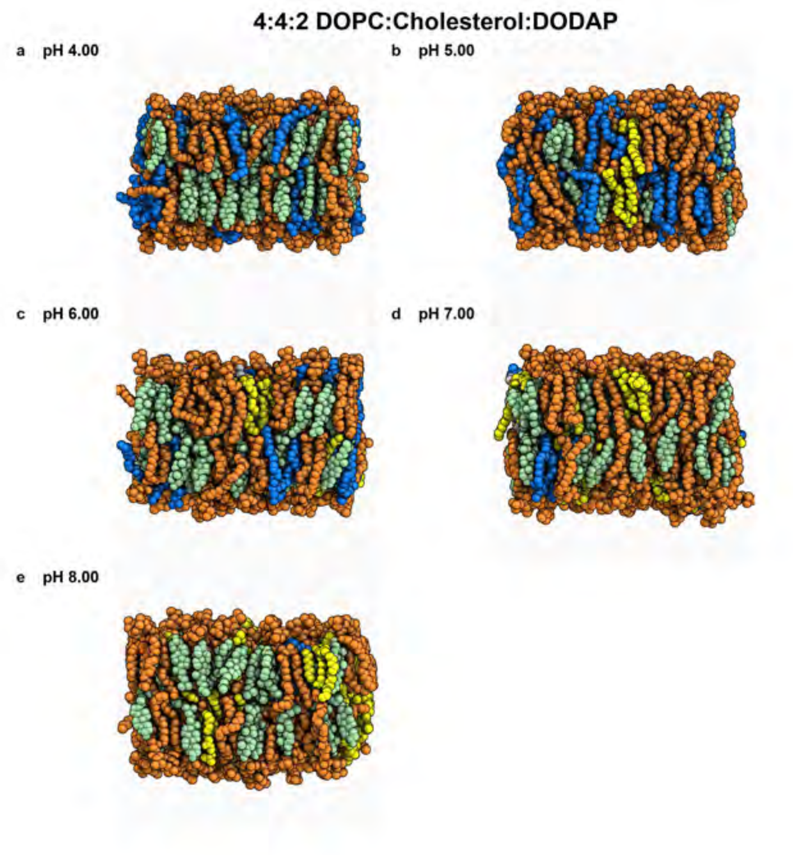


**Fig. 11** Snapshots of the membrane in the final frame of the simulation at the corresponding pH values. The helper lipid is shown in orange, cholesterol in green, the protonated aminolipid (net charge +1) in blue, and the deprotonated aminolipid (net charge 0) in yellow.

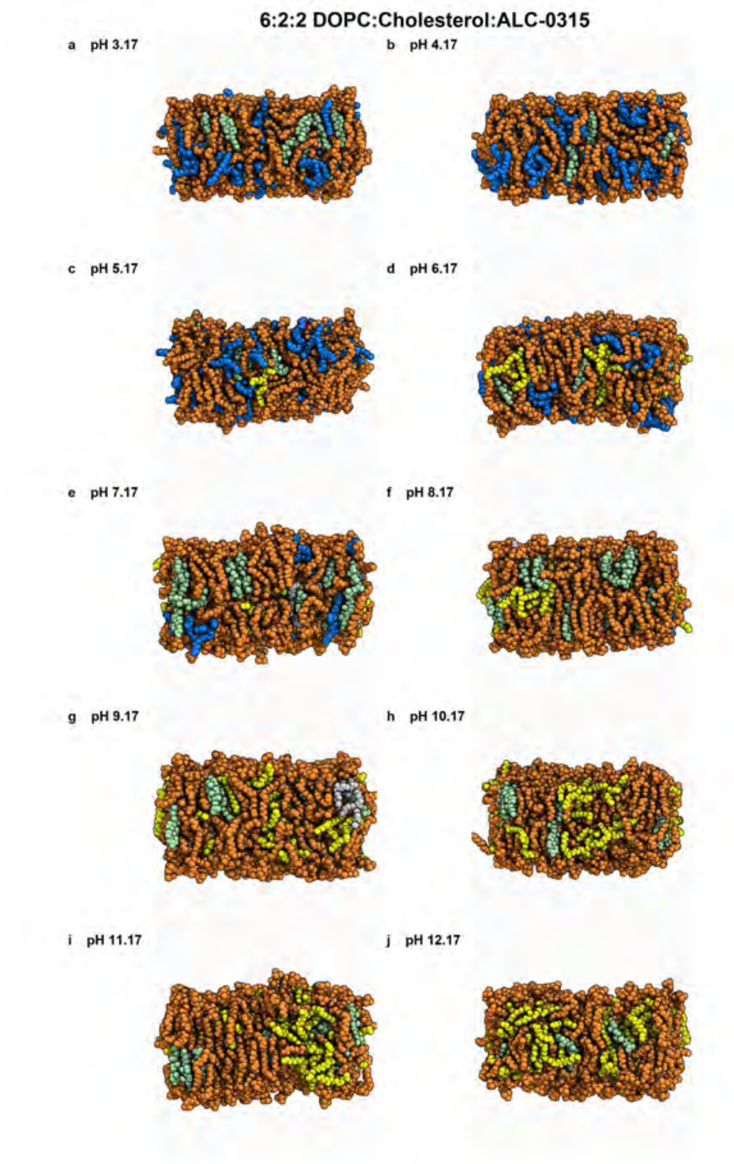


**Fig. 12** Snapshots of the membrane in the final frame of the simulation at the corresponding pH values. The helper lipid is shown in orange, cholesterol in green, the protonated aminolipid (net charge +1) in blue, and the deprotonated aminolipid (net charge 0) in yellow.

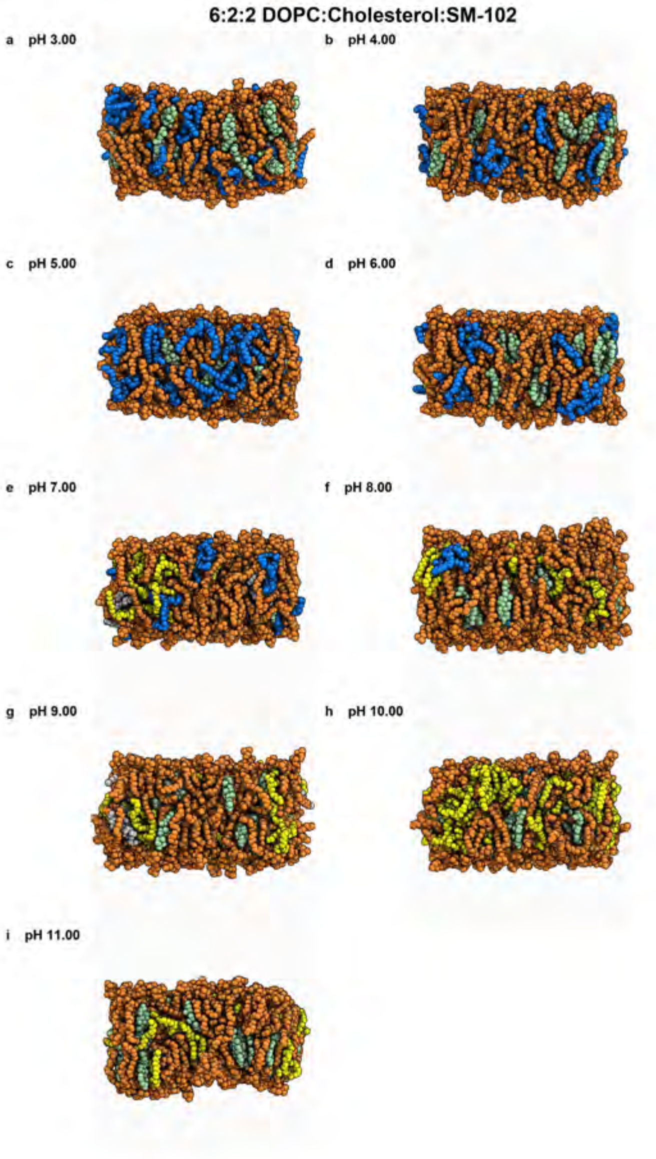


**Fig. 13** Snapshots of the membrane in the final frame of the simulation at the corresponding pH values. The helper lipid is shown in orange, cholesterol in green, the protonated aminolipid (net charge +1) in blue, and the deprotonated aminolipid (net charge 0) in yellow.

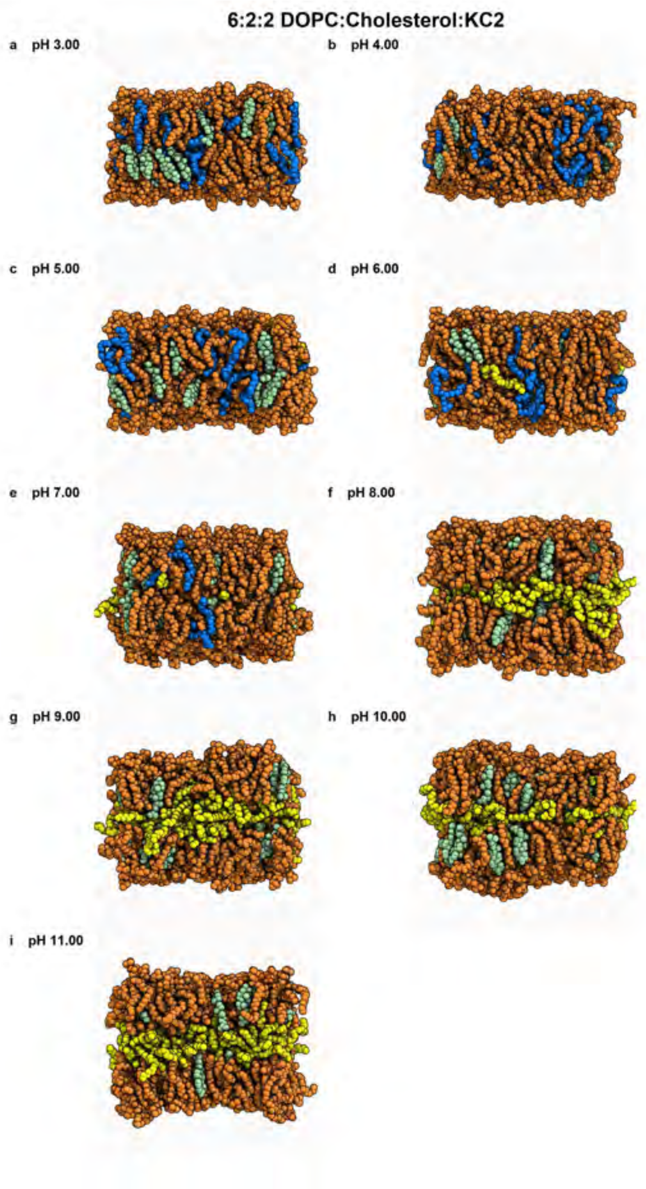


**Fig. 14** Snapshots of the membrane in the final frame of the simulation at the corresponding pH values. The helper lipid is shown in orange, cholesterol in green, the protonated aminolipid (net charge +1) in blue, and the deprotonated aminolipid (net charge 0) in yellow.

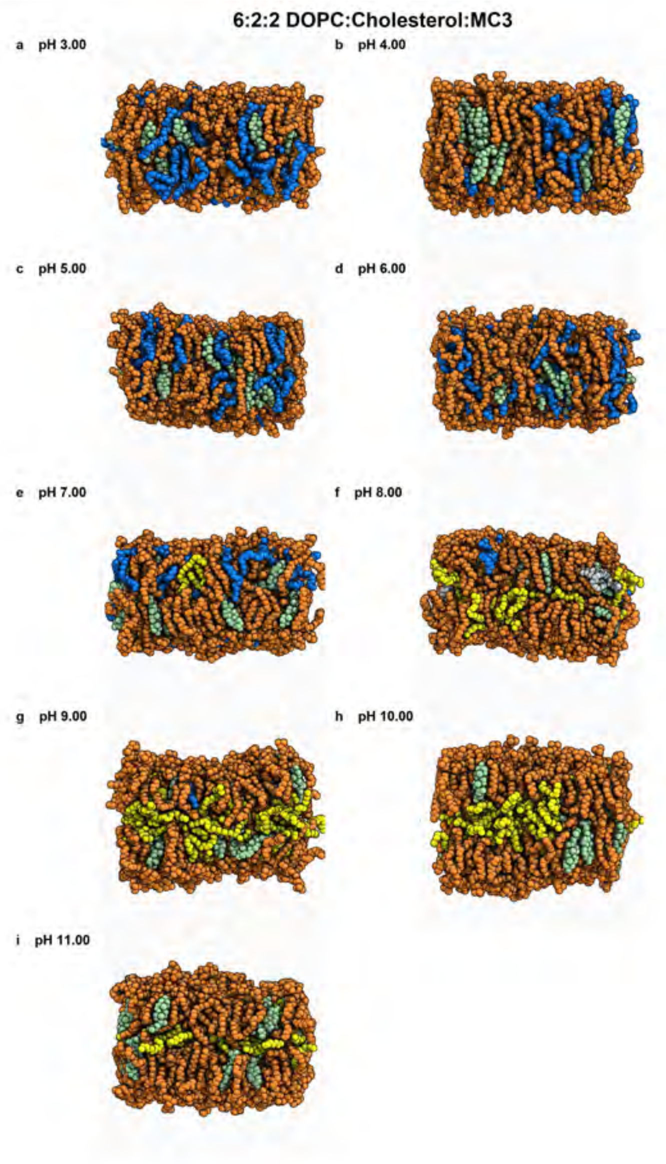


**Fig. 15** Snapshots of the membrane in the final frame of the simulation at the corresponding pH values. The helper lipid is shown in orange, cholesterol in green, the protonated aminolipid (net charge +1) in blue, and the deprotonated aminolipid (net charge 0) in yellow.

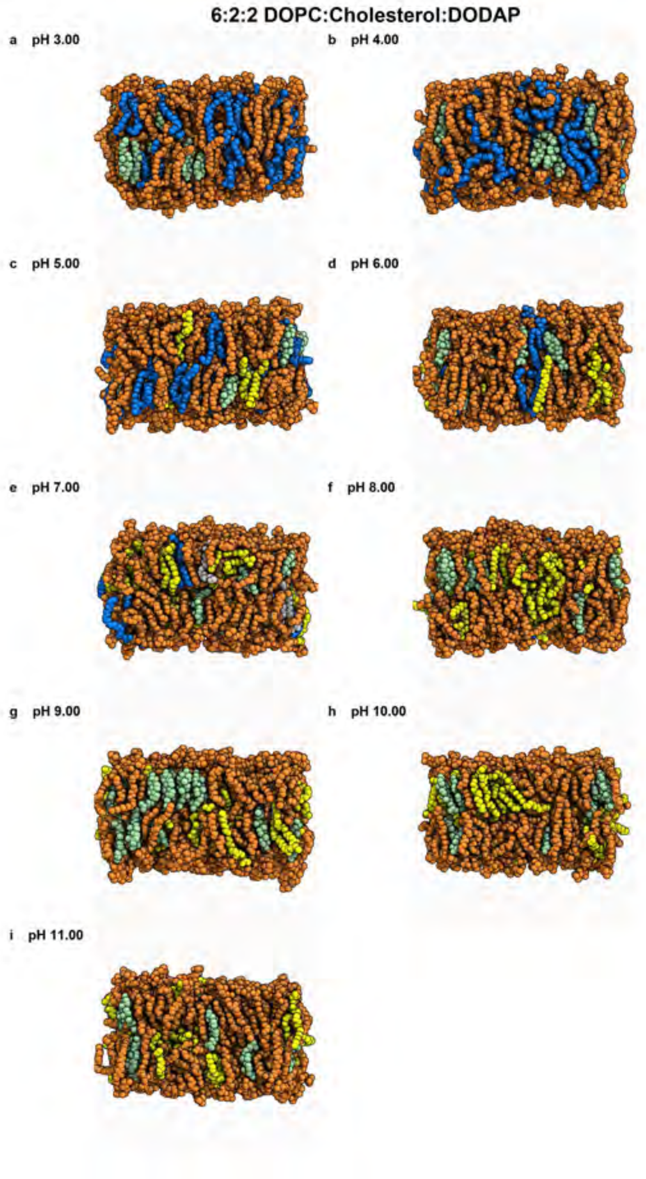


**Fig. 16** Snapshots of the membrane in the final frame of the simulation at the corresponding pH values. The helper lipid is shown in orange, cholesterol in green, the protonated aminolipid (net charge +1) in blue, and the deprotonated aminolipid (net charge 0) in yellow.

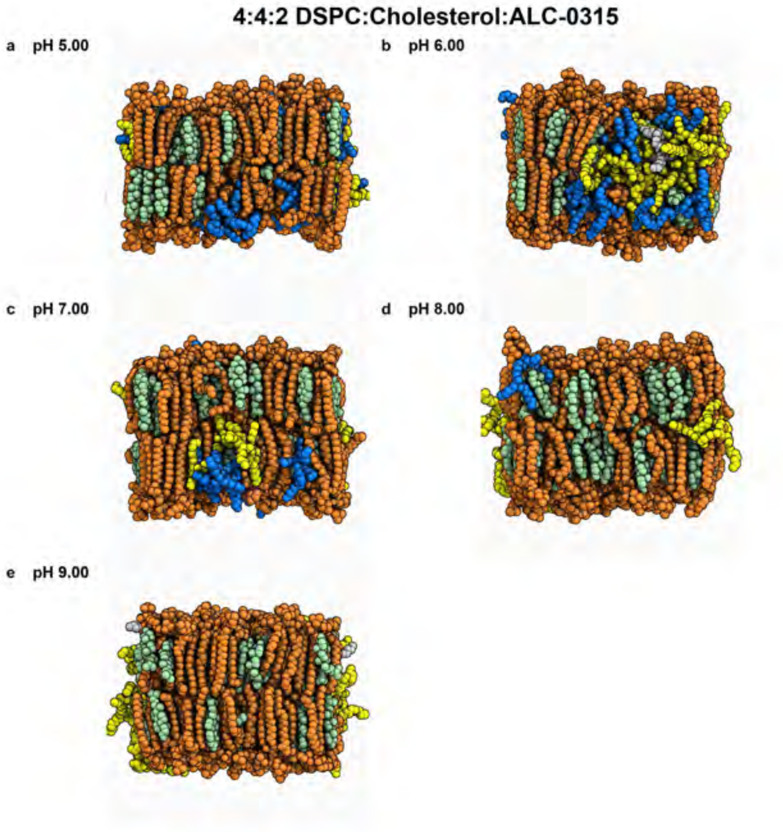


**Fig. 17** Snapshots of the membrane in the final frame of the simulation at the corresponding pH values. The helper lipid is shown in orange, cholesterol in green, the protonated aminolipid (net charge +1) in blue, and the deprotonated aminolipid (net charge 0) in yellow.

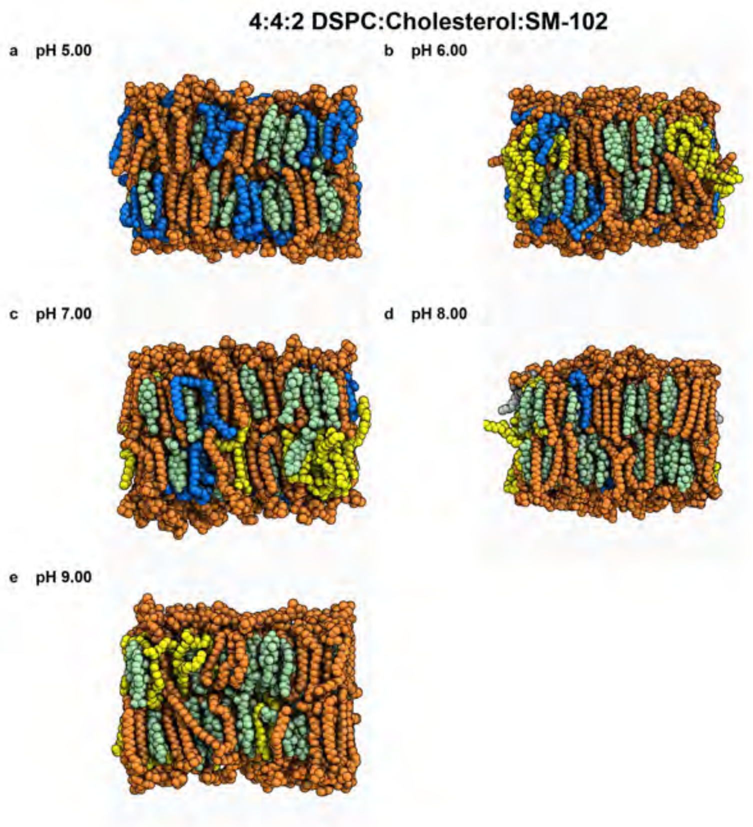


**Fig. 18** Snapshots of the membrane in the final frame of the simulation at the corresponding pH values. The helper lipid is shown in orange, cholesterol in green, the protonated aminolipid (net charge +1) in blue, and the deprotonated aminolipid (net charge 0) in yellow.

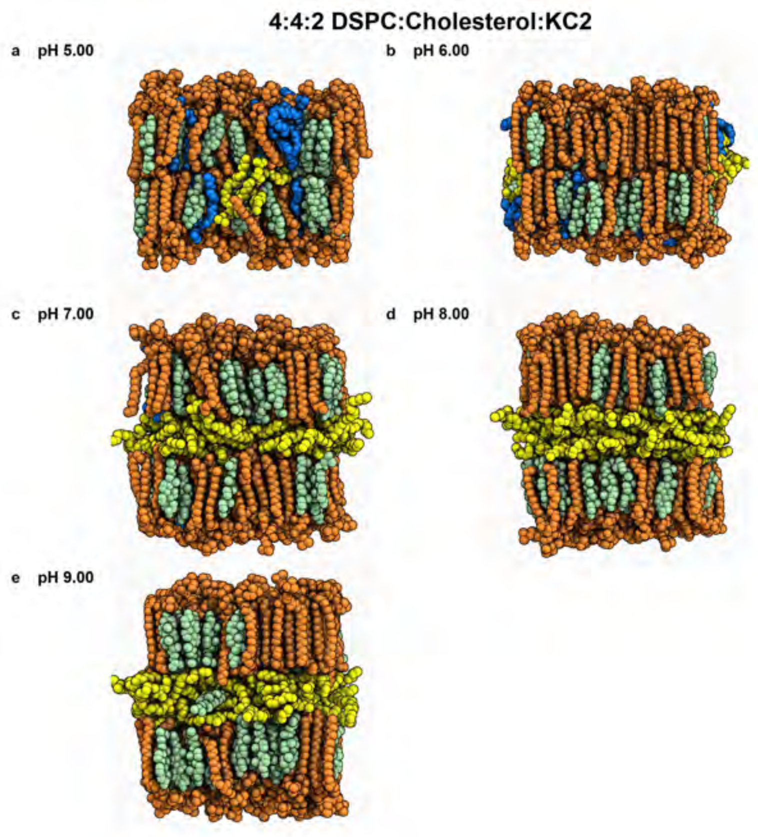


**Fig. 19** Snapshots of the membrane in the final frame of the simulation at the corresponding pH values. The helper lipid is shown in orange, cholesterol in green, the protonated aminolipid (net charge +1) in blue, and the deprotonated aminolipid (net charge 0) in yellow.

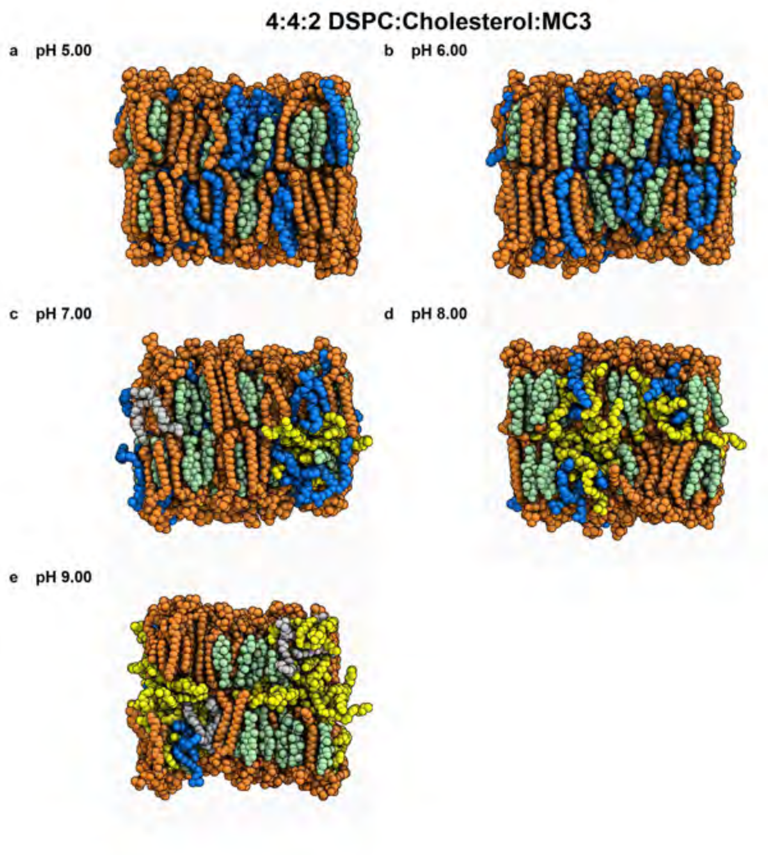


**Fig. 20** Snapshots of the membrane in the final frame of the simulation at the corresponding pH values. The helper lipid is shown in orange, cholesterol in green, the protonated aminolipid (net charge +1) in blue, and the deprotonated aminolipid (net charge 0) in yellow.

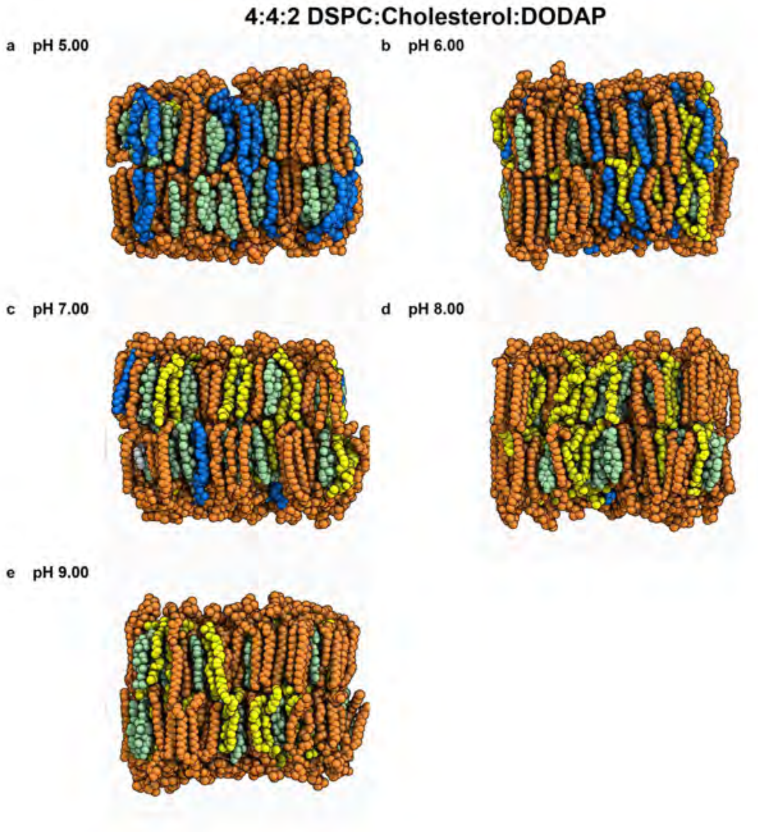


**Fig. 21** Snapshots of the membrane in the final frame of the simulation at the corresponding pH values. The helper lipid is shown in orange, cholesterol in green, the protonated aminolipid (net charge +1) in blue, and the deprotonated aminolipid (net charge 0) in yellow.

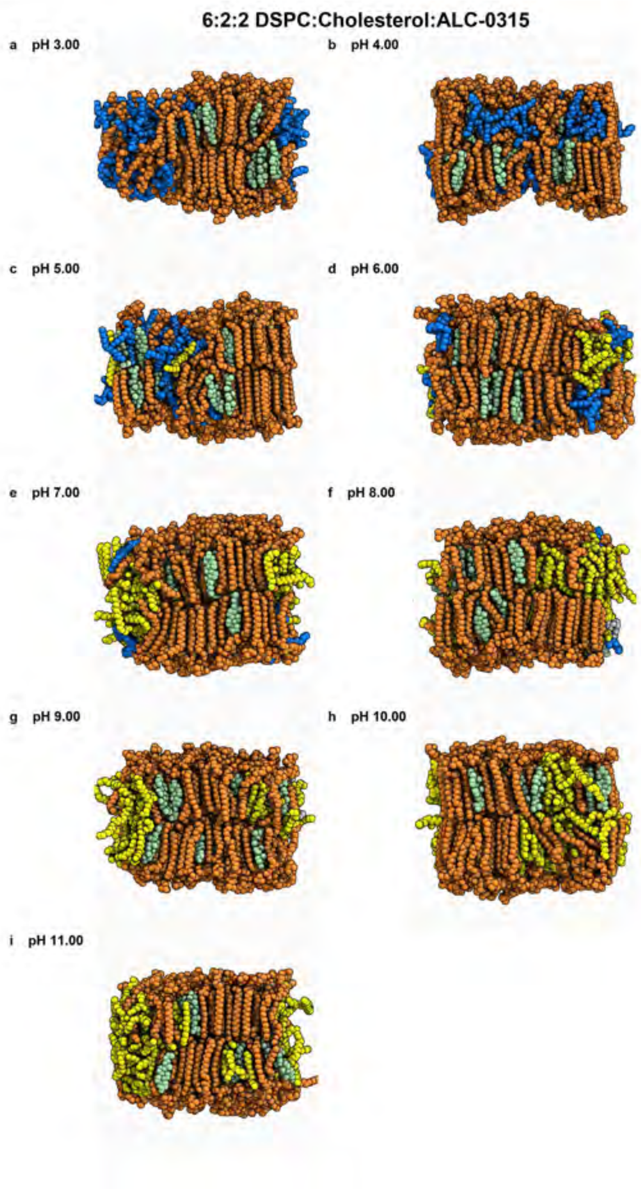


**Fig. 22** Snapshots of the membrane in the final frame of the simulation at the corresponding pH values. The helper lipid is shown in orange, cholesterol in green, the protonated aminolipid (net charge +1) in blue, and the deprotonated aminolipid (net charge 0) in yellow.

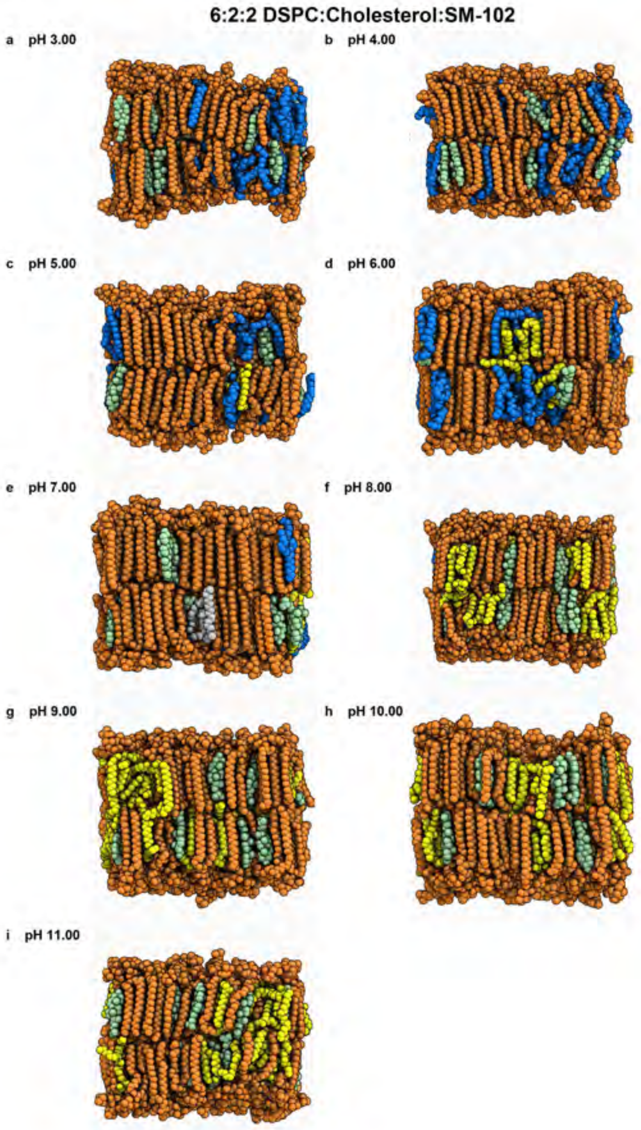


**Fig. 23** Snapshots of the membrane in the final frame of the simulation at the corresponding pH values. The helper lipid is shown in orange, cholesterol in green, the protonated aminolipid (net charge +1) in blue, and the deprotonated aminolipid (net charge 0) in yellow.

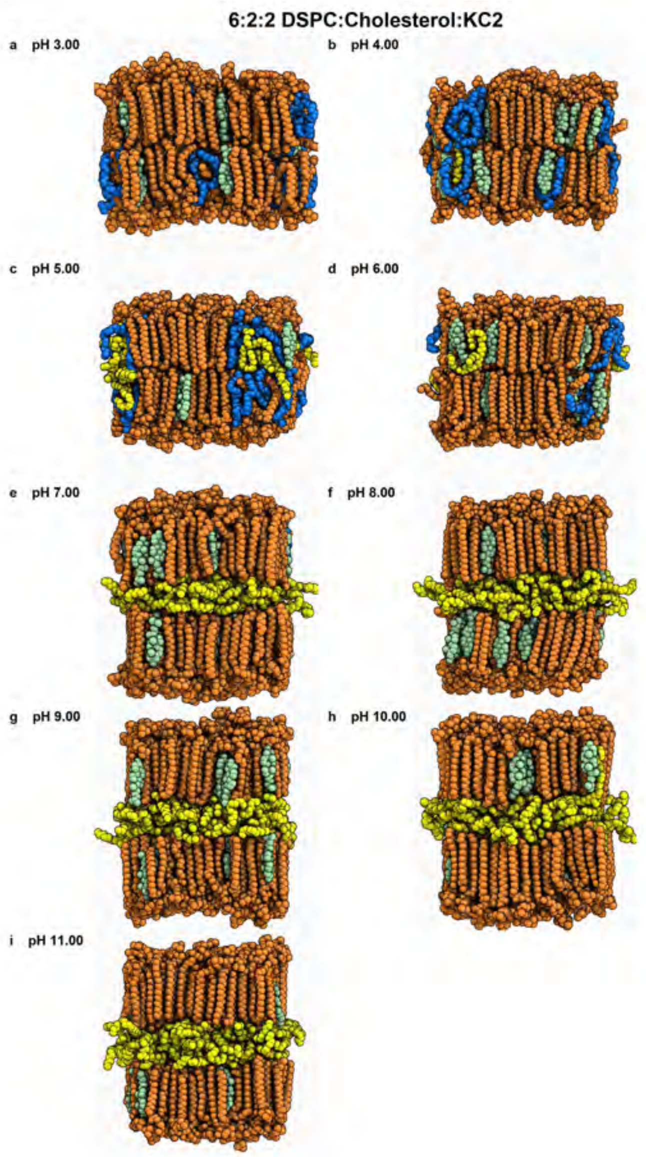


**Fig. 24** Snapshots of the membrane in the final frame of the simulation at the corresponding pH values. The helper lipid is shown in orange, cholesterol in green, the protonated aminolipid (net charge +1) in blue, and the deprotonated aminolipid (net charge 0) in yellow.

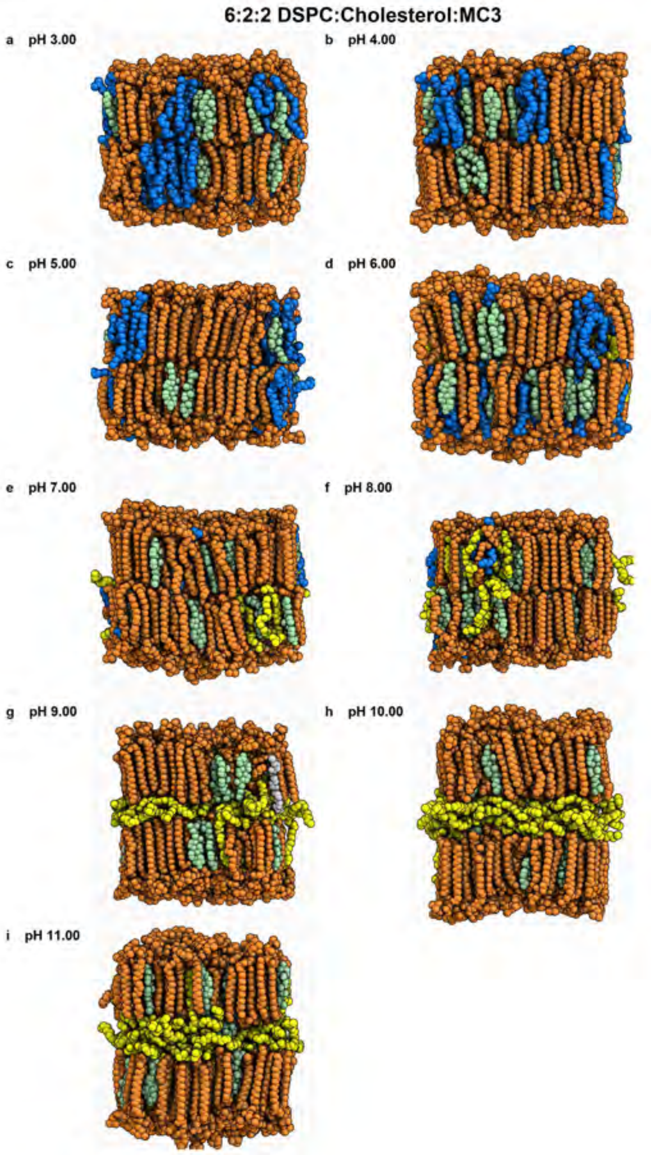


**Fig. 25** Snapshots of the membrane in the final frame of the simulation at the corresponding pH values. The helper lipid is shown in orange, cholesterol in green, the protonated aminolipid (net charge +1) in blue, and the deprotonated aminolipid (net charge 0) in yellow.

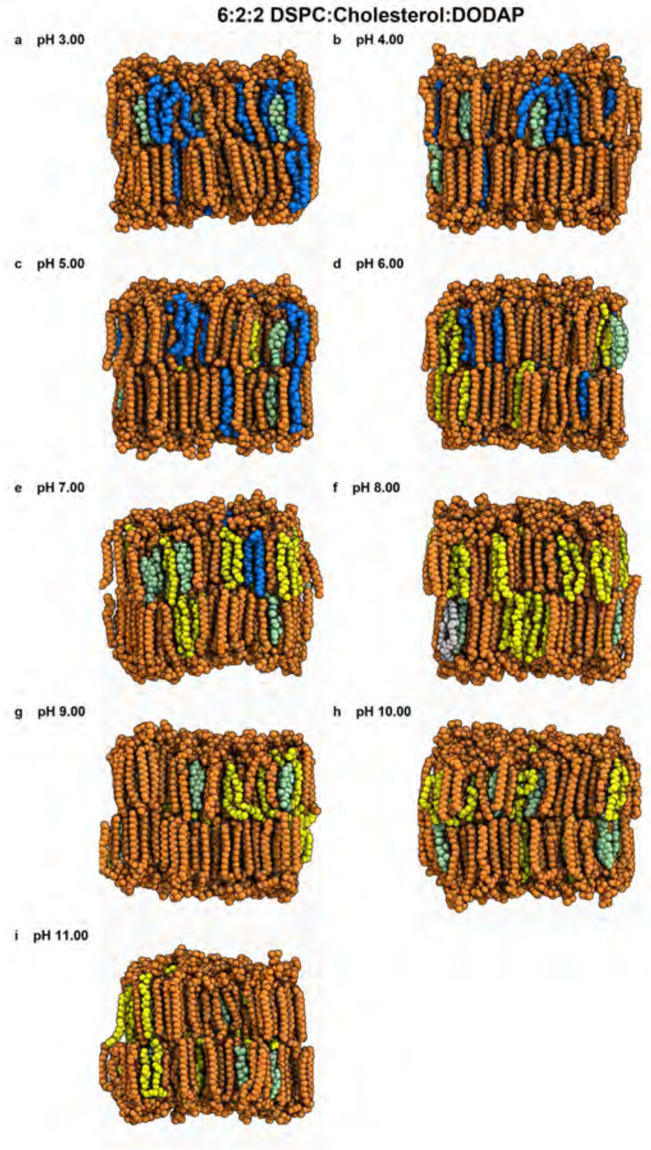


**Fig. 26** Snapshots of the membrane in the final frame of the simulation at the corresponding pH values. The helper lipid is shown in orange, cholesterol in green, the protonated aminolipid (net charge +1) in blue, and the deprotonated aminolipid (net charge 0) in yellow.

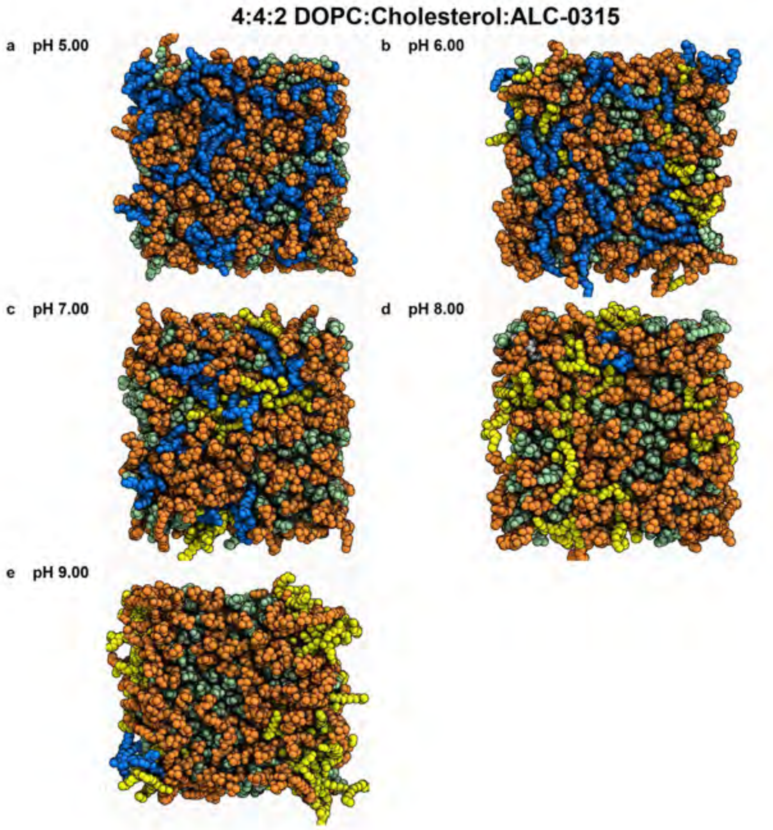


**Fig. 27** Snapshots of the membrane in the final frame of the simulation at the corresponding pH values. The helper lipid is shown in orange, cholesterol in green, the protonated aminolipid (net charge +1) in blue, and the deprotonated aminolipid (net charge 0) in yellow.

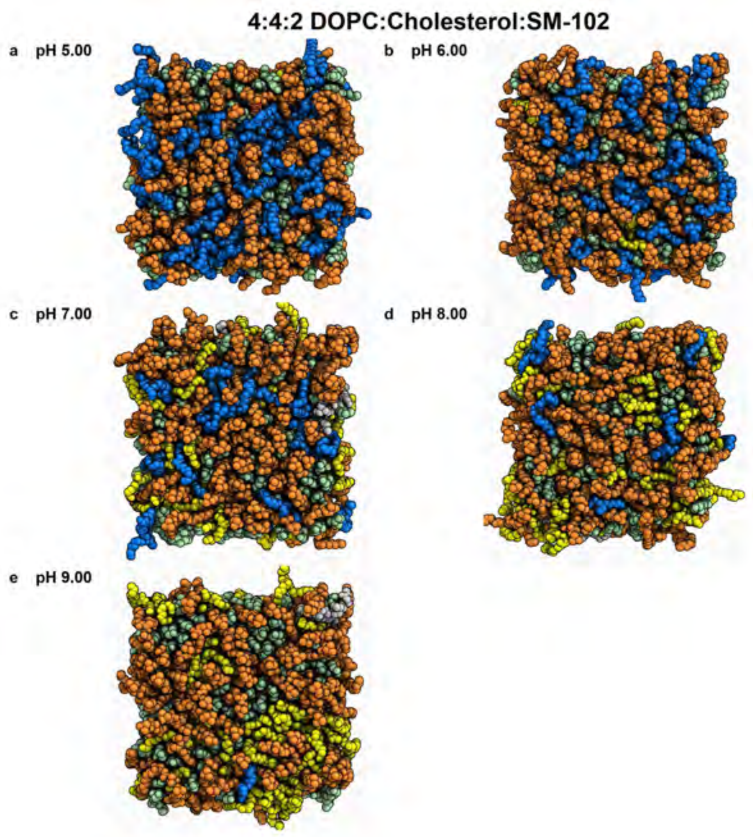


**Fig. 28** Snapshots of the membrane in the final frame of the simulation at the corresponding pH values. The helper lipid is shown in orange, cholesterol in green, the protonated aminolipid (net charge +1) in blue, and the deprotonated aminolipid (net charge 0) in yellow.

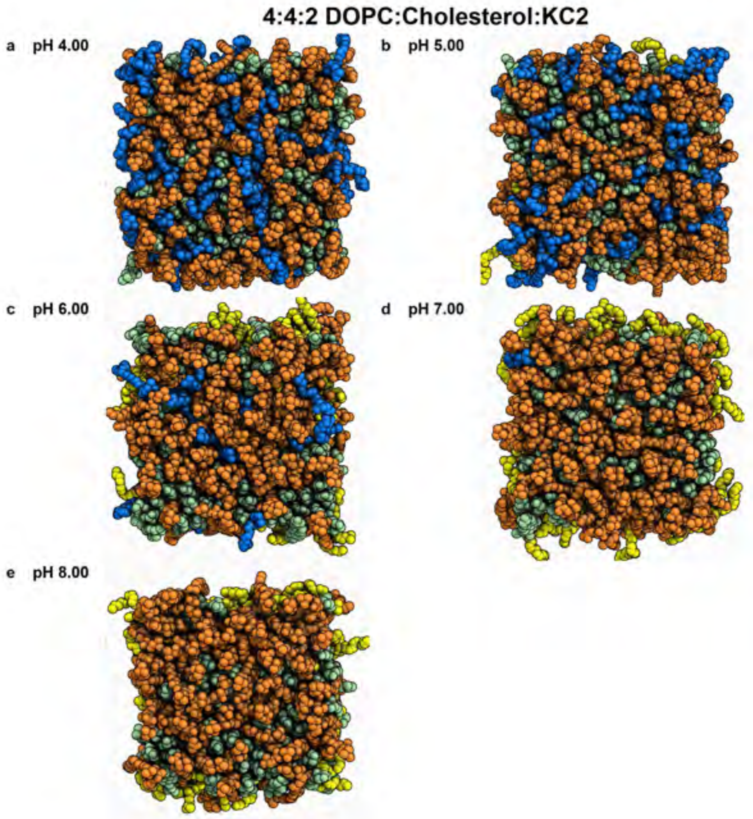


**Fig. 29** Snapshots of the membrane in the final frame of the simulation at the corresponding pH values. The helper lipid is shown in orange, cholesterol in green, the protonated aminolipid (net charge +1) in blue, and the deprotonated aminolipid (net charge 0) in yellow.

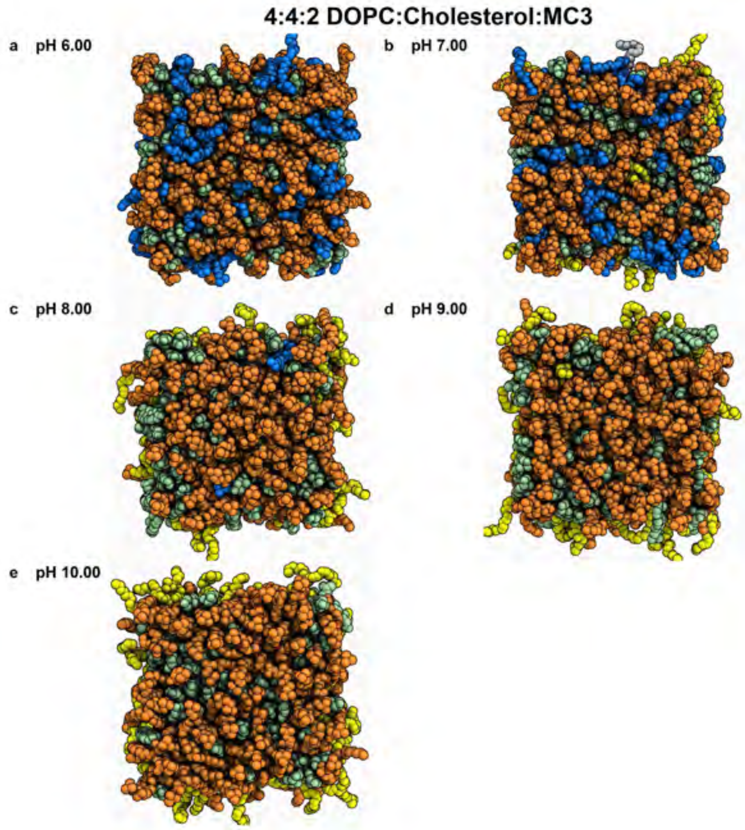


**Fig. 30** Snapshots of the membrane in the final frame of the simulation at the corresponding pH values. The helper lipid is shown in orange, cholesterol in green, the protonated aminolipid (net charge +1) in blue, and the deprotonated aminolipid (net charge 0) in yellow.

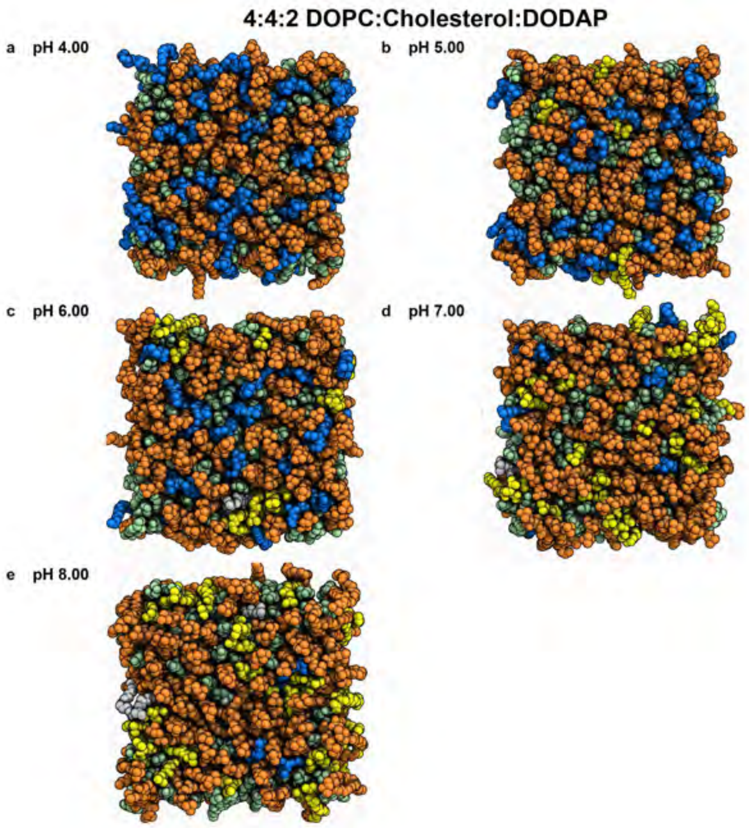


**Fig. 31** Snapshots of the membrane in the final frame of the simulation at the corresponding pH values. The helper lipid is shown in orange, cholesterol in green, the protonated aminolipid (net charge +1) in blue, and the deprotonated aminolipid (net charge 0) in yellow.

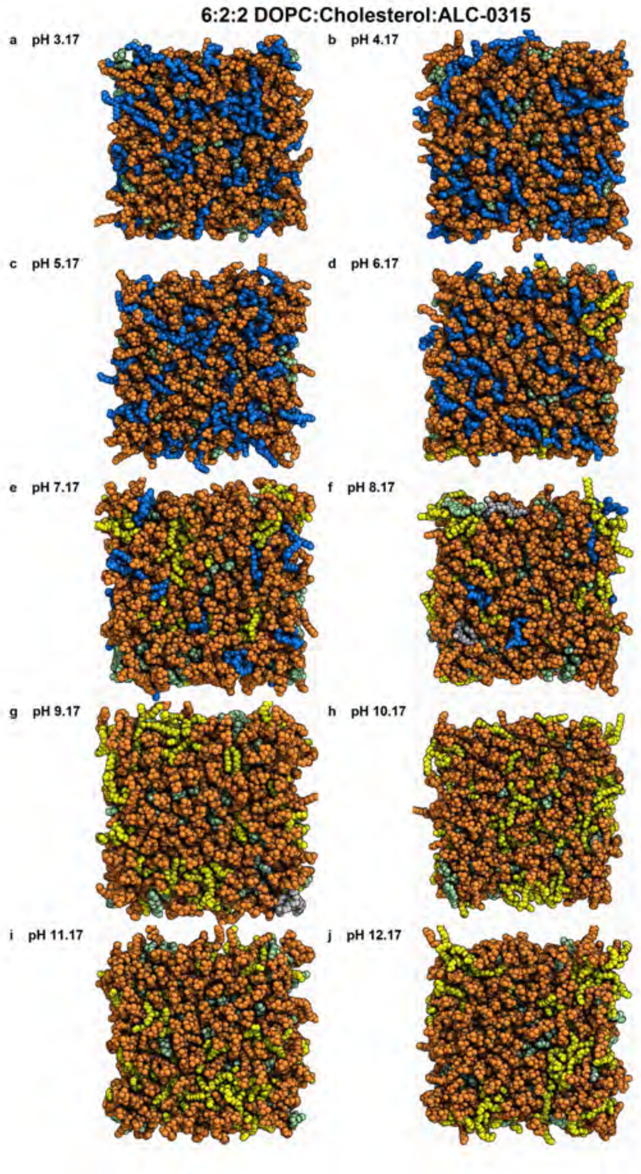


**Fig. 32** Snapshots of the membrane in the final frame of the simulation at the corresponding pH values. The helper lipid is shown in orange, cholesterol in green, the protonated aminolipid (net charge +1) in blue, and the deprotonated aminolipid (net charge 0) in yellow.

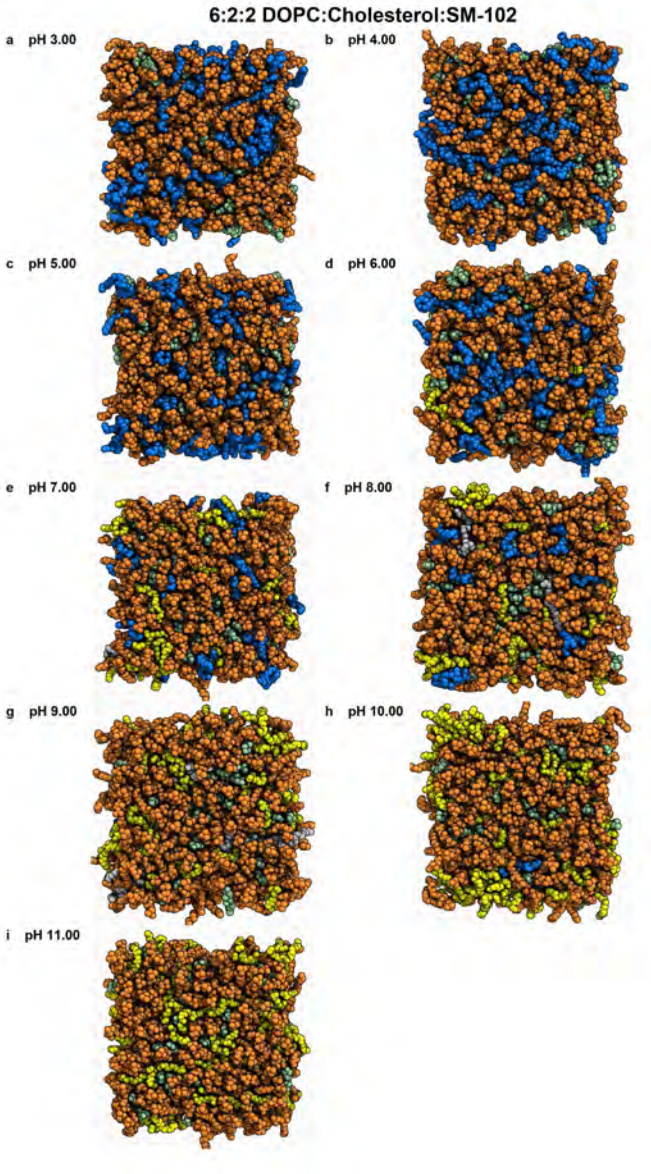


**Fig. 33** Snapshots of the membrane in the final frame of the simulation at the corresponding pH values. The helper lipid is shown in orange, cholesterol in green, the protonated aminolipid (net charge +1) in blue, and the deprotonated aminolipid (net charge 0) in yellow.

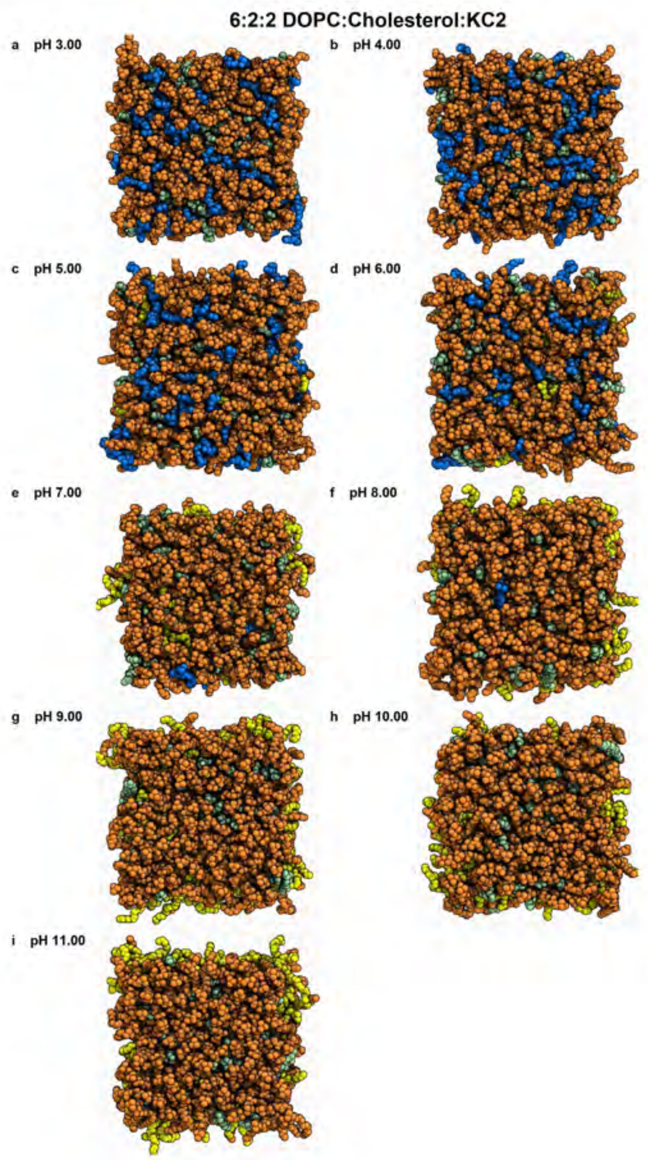


**Fig. 34** Snapshots of the membrane in the final frame of the simulation at the corresponding pH values. The helper lipid is shown in orange, cholesterol in green, the protonated aminolipid (net charge +1) in blue, and the deprotonated aminolipid (net charge 0) in yellow.

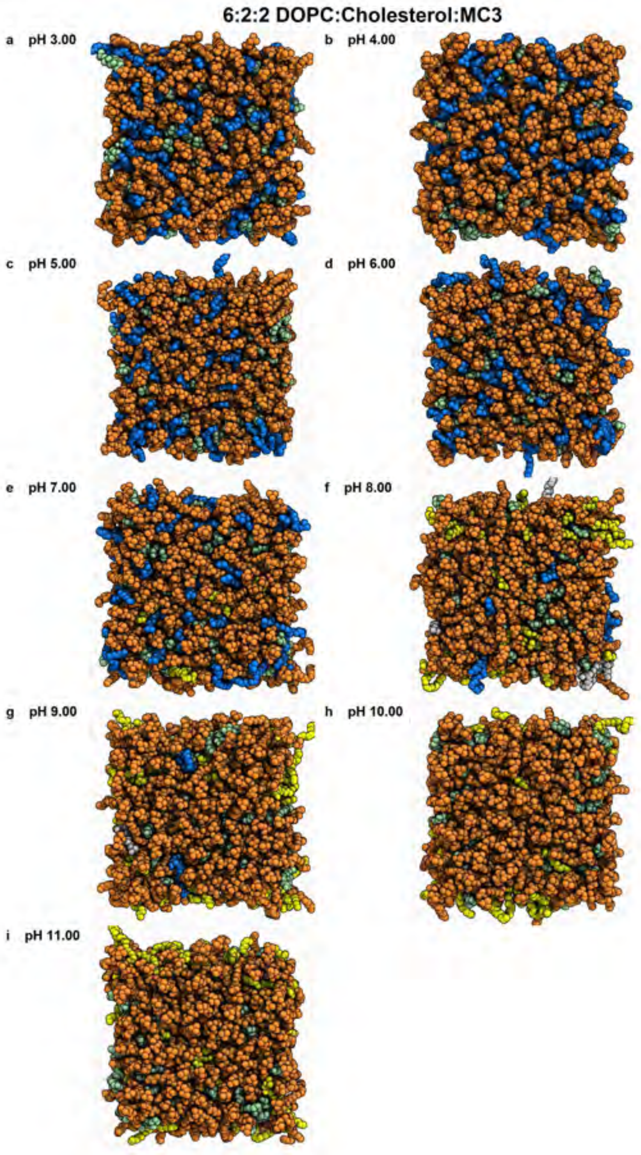


**Fig. 35** Snapshots of the membrane in the final frame of the simulation at the corresponding pH values. The helper lipid is shown in orange, cholesterol in green, the protonated aminolipid (net charge +1) in blue, and the deprotonated aminolipid (net charge 0) in yellow.

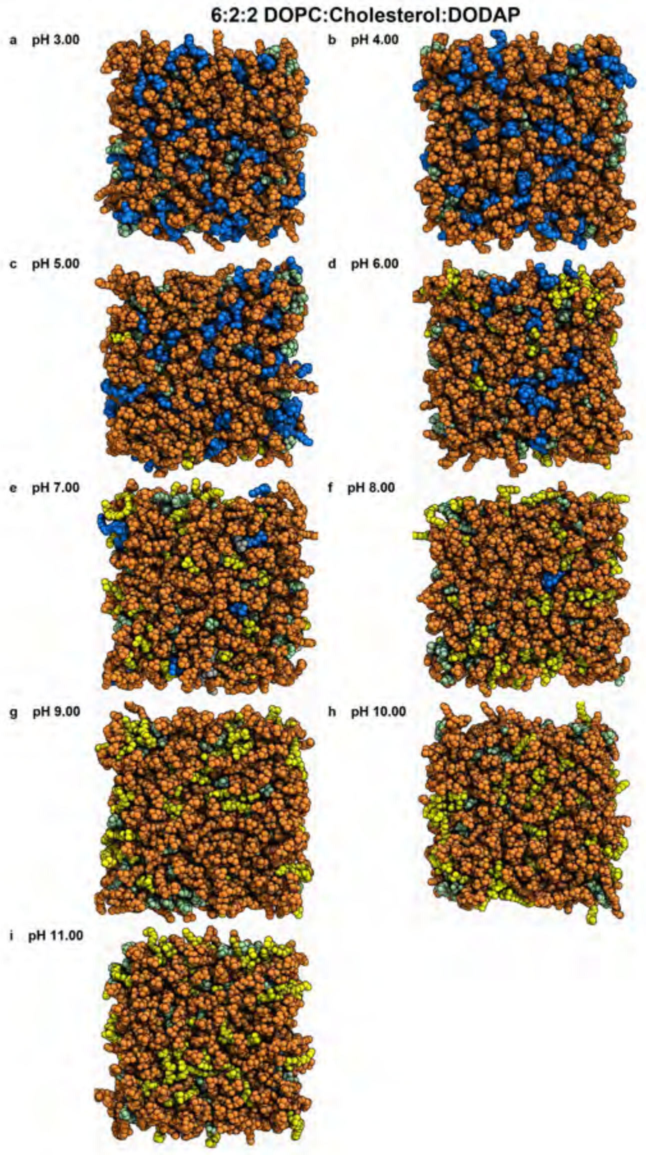


**Fig. 36** Snapshots of the membrane in the final frame of the simulation at the corresponding pH values. The helper lipid is shown in orange, cholesterol in green, the protonated aminolipid (net charge +1) in blue, and the deprotonated aminolipid (net charge 0) in yellow.

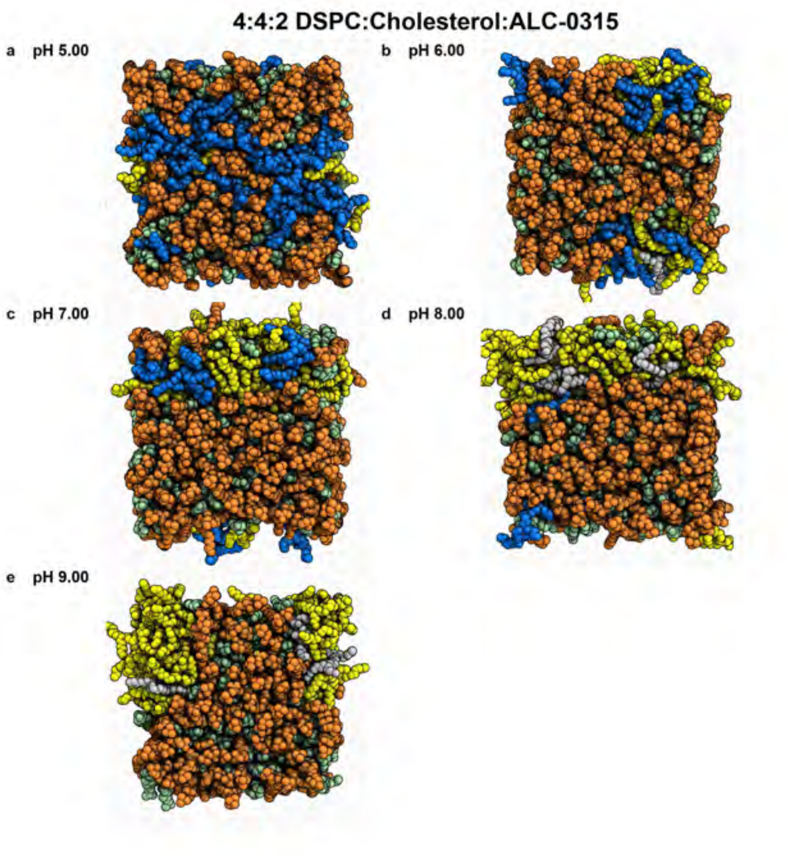


**Fig. 37** Snapshots of the membrane in the final frame of the simulation at the corresponding pH values. The helper lipid is shown in orange, cholesterol in green, the protonated aminolipid (net charge +1) in blue, and the deprotonated aminolipid (net charge 0) in yellow.

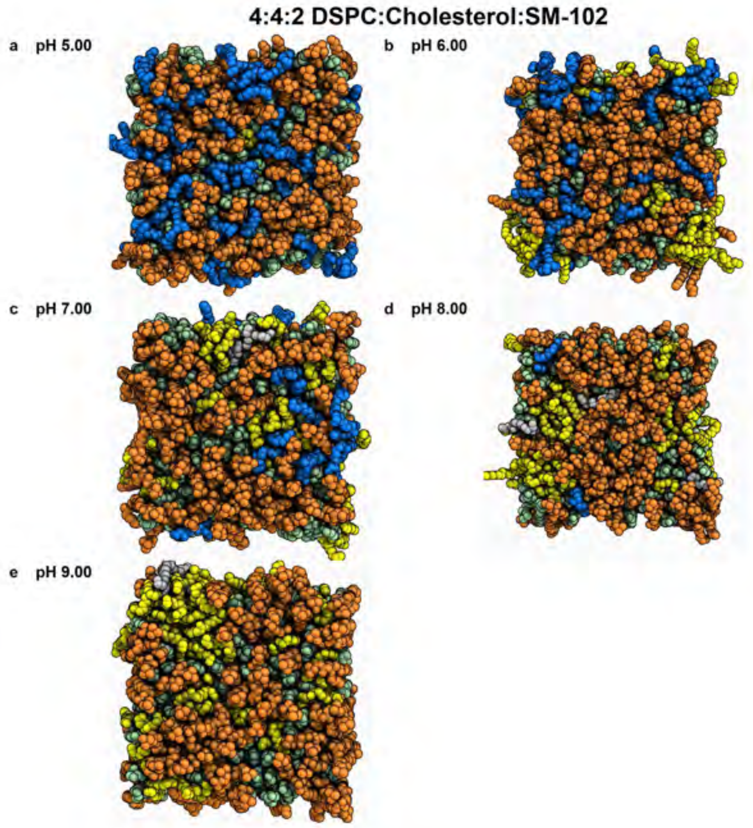


**Fig. 38** Snapshots of the membrane in the final frame of the simulation at the corresponding pH values. The helper lipid is shown in orange, cholesterol in green, the protonated aminolipid (net charge +1) in blue, and the deprotonated aminolipid (net charge 0) in yellow.

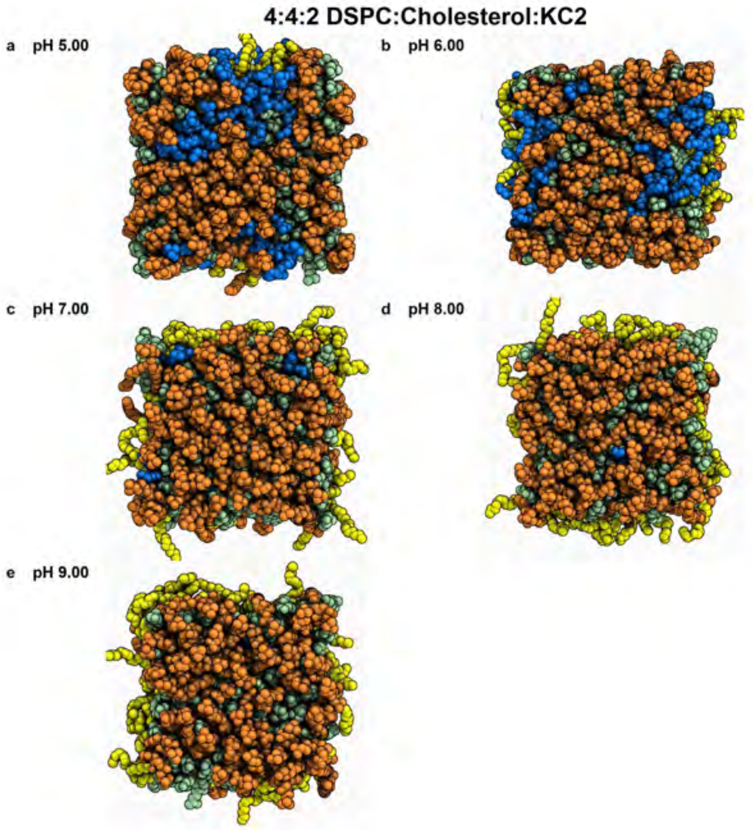


**Fig. 39** Snapshots of the membrane in the final frame of the simulation at the corresponding pH values. The helper lipid is shown in orange, cholesterol in green, the protonated aminolipid (net charge +1) in blue, and the deprotonated aminolipid (net charge 0) in yellow.

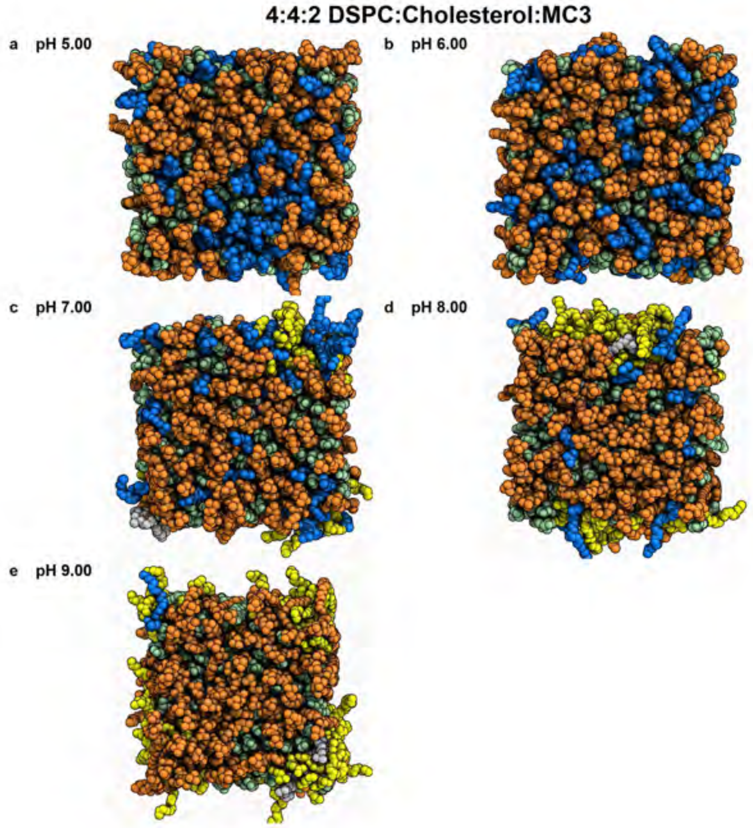
4:4:2 DSPC:Cholesterol:MC3
a pH 5.00
b pH 6.00
c pH 7.00
d pH 8.00
e pH 9.00

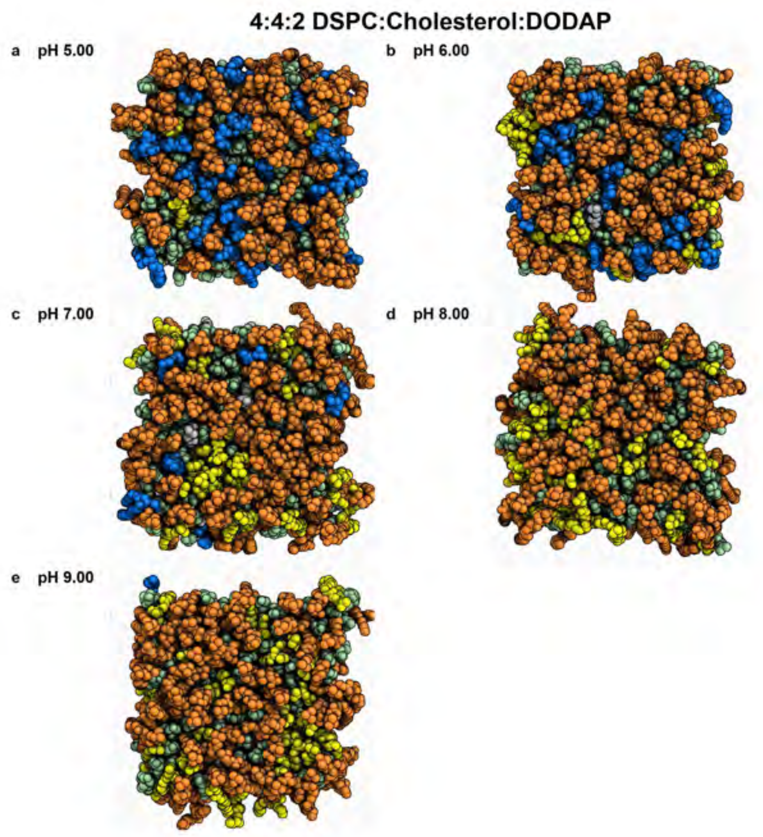


**Fig. 40** Snapshots of the membrane in the final frame of the simulation at the corresponding pH values. The helper lipid is shown in orange, cholesterol in green, the protonated aminolipid (net charge +1) in blue, and the deprotonated aminolipid (net charge 0) in yellow.

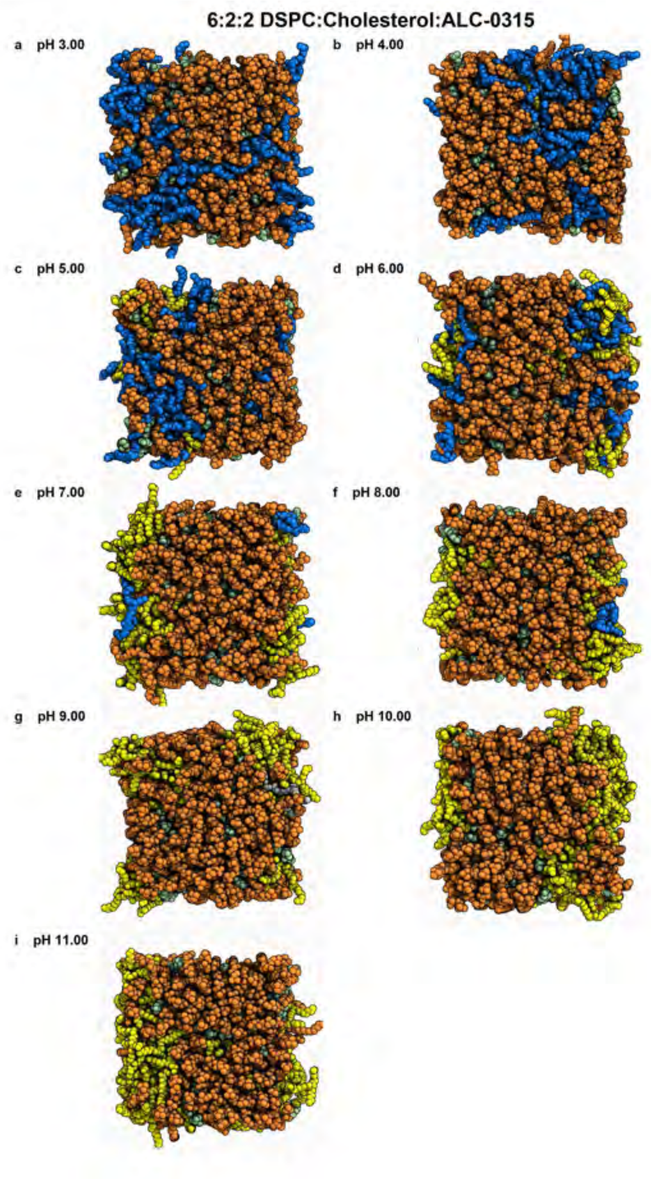


**Fig. 41** Snapshots of the membrane in the final frame of the simulation at the corresponding pH values. The helper lipid is shown in orange, cholesterol in green, the protonated aminolipid (net charge +1) in blue, and the deprotonated aminolipid (net charge 0) in yellow.

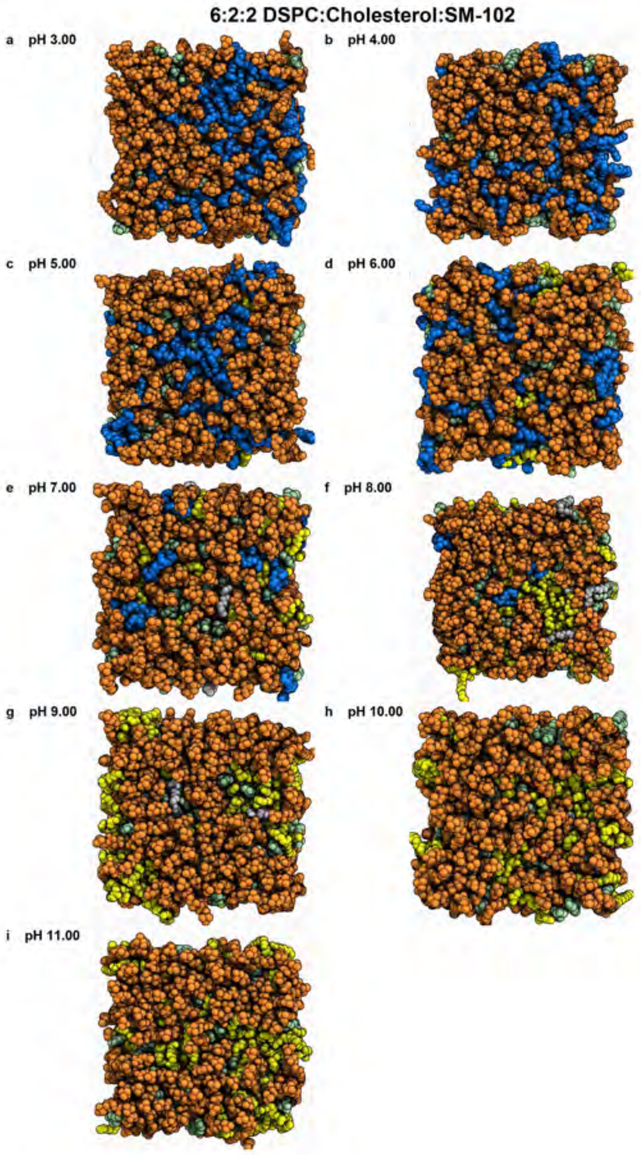


**Fig. 42** Snapshots of the membrane in the final frame of the simulation at the corresponding pH values. The helper lipid is shown in orange, cholesterol in green, the protonated aminolipid (net charge +1) in blue, and the deprotonated aminolipid (net charge 0) in yellow.

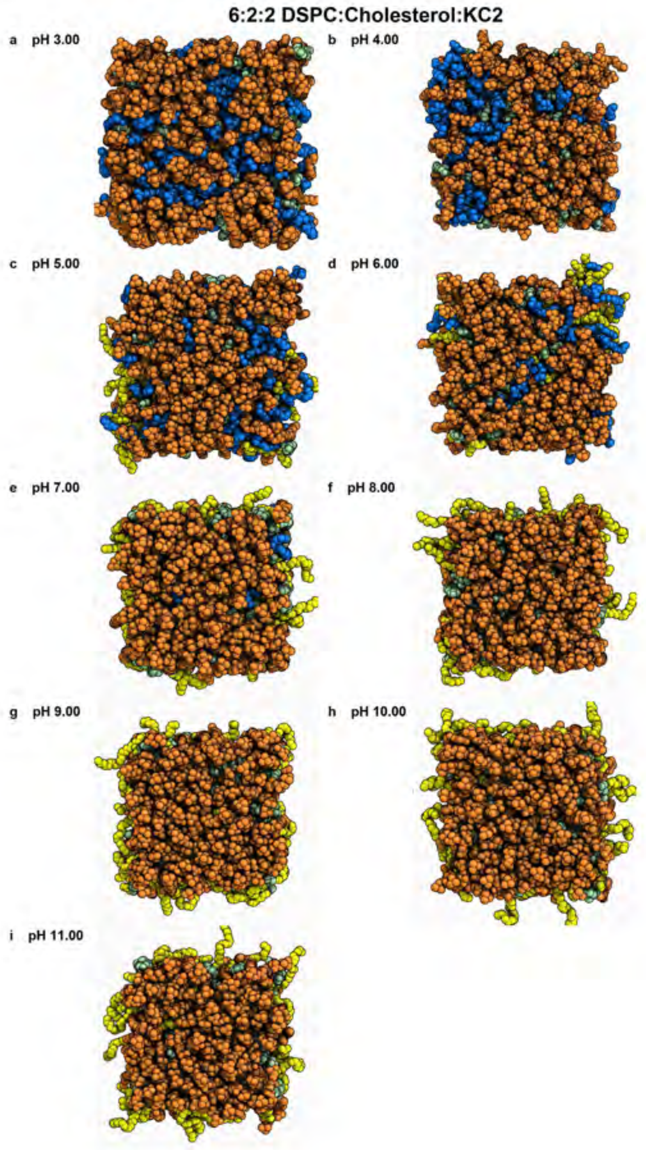


**Fig. 43** Snapshots of the membrane in the final frame of the simulation at the corresponding pH values. The helper lipid is shown in orange, cholesterol in green, the protonated aminolipid (net charge +1) in blue, and the deprotonated aminolipid (net charge 0) in yellow.

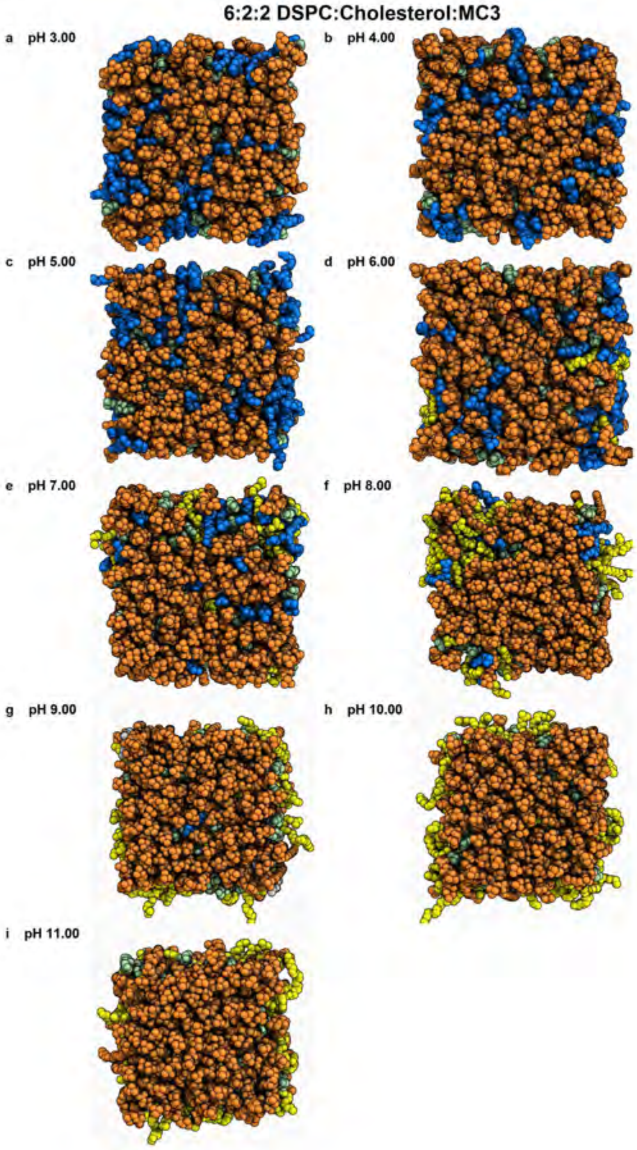
6:2:2 DSPC:Cholesterol:MC3
a pH 3.00
b pH 4.00
c pH 5.00
d pH 6.00
e pH 7.00
f pH 8.00
g pH 9.00
h pH 10.00
i pH 11.00

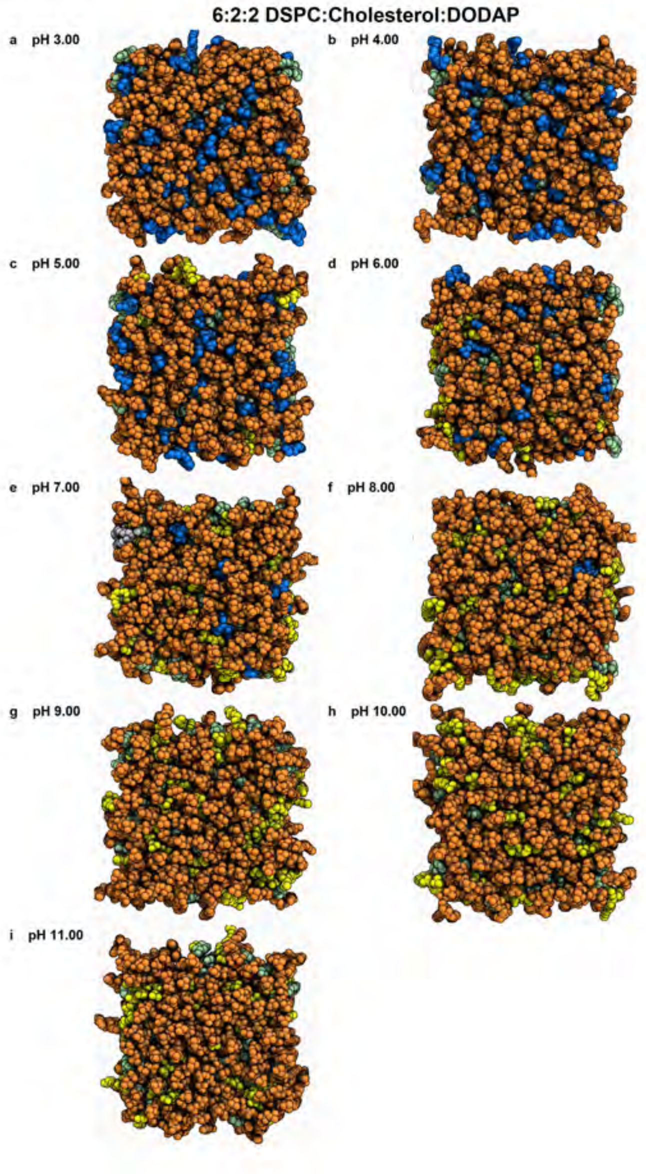


**Fig. 44** Snapshots of the membrane in the final frame of the simulation at the corresponding pH values. The helper lipid is shown in orange, cholesterol in green, the protonated aminolipid (net charge +1) in blue, and the deprotonated aminolipid (net charge 0) in yellow.

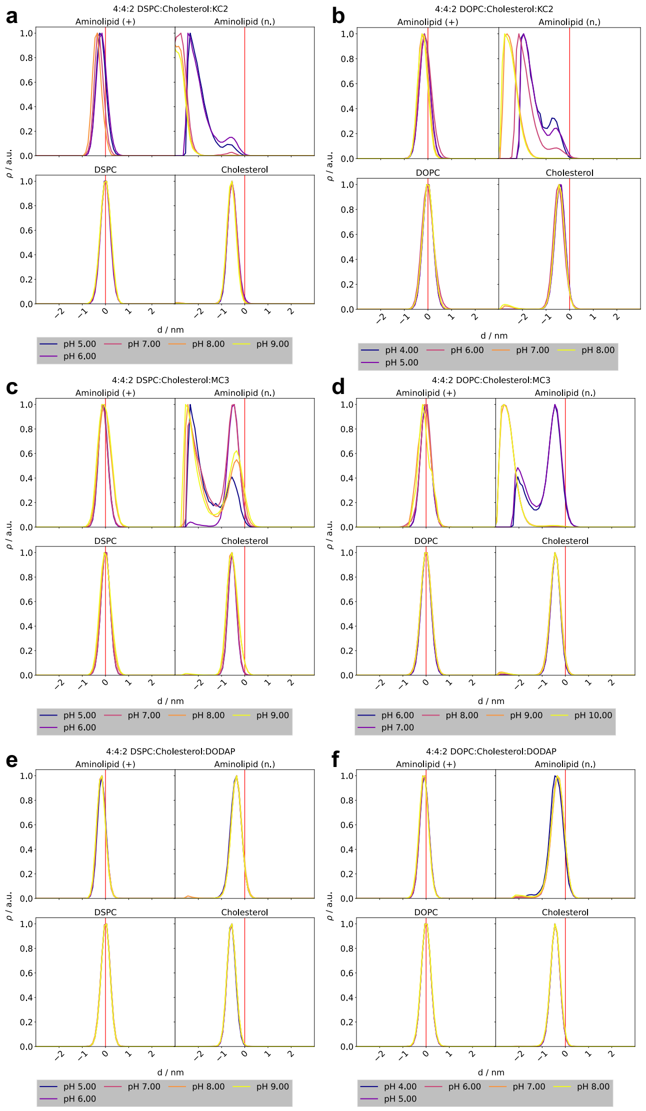


**Fig. 45** Lipid density profiles along the z-axis for membranes with a 4:4:2 helper lipid:cholesterol:aminolipid composition containing DLin-KC2-DMA, DLin-MC3-DMA, and DODAP. Densities were calculated using one representative atom per lipid (oxygen for cholesterol, nitrogen for the aminolipid, and phosphorus for the helper lipid). Positions are reported relative to the membrane surface, defined as the average z-position of the phosphorus atoms in the helper lipid headgroups of each leaflet. Negative distances (d) correspond to the membrane core, whereas positive distances correspond to the solvent phase. Results were averaged over both leaflets.

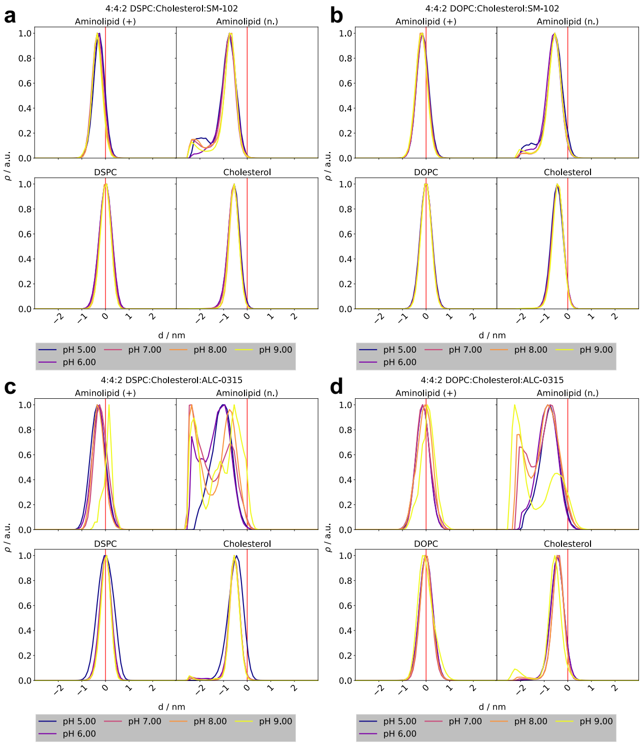


**Fig. 46** Lipid density profiles along the z-axis for membranes with a 4:4:2 helper lipid:cholesterol:aminolipid composition containing SM-102 and ALC-0315. Densities were calculated using one representative atom per lipid (oxygen for cholesterol, nitrogen for the aminolipid, and phosphorus for the helper lipid). Positions are reported relative to the membrane surface, defined as the average z-position of the phosphorus atoms in the helper lipid headgroups of each leaflet. Negative distances (d) correspond to the membrane core, whereas positive distances correspond to the solvent phase. Results were averaged over both leaflets.

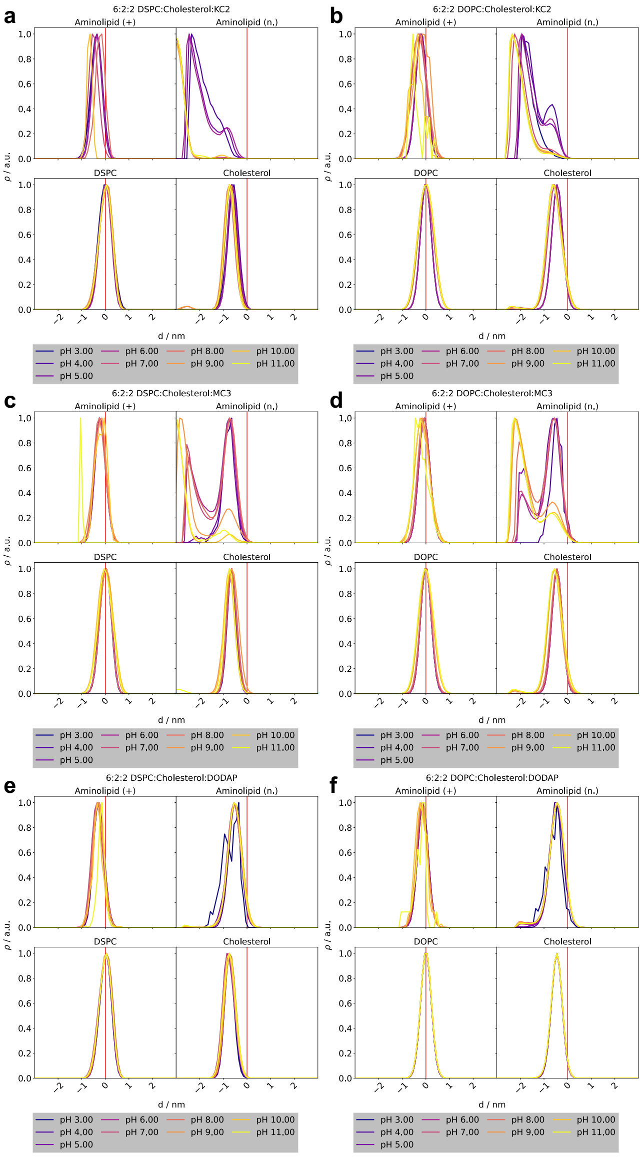


**Fig. 47** Lipid density profiles along the z-axis for membranes with a 6:2:2 helper lipid:cholesterol:aminolipid composition containing DLin-KC2-DMA, DLin-MC3-DMA, and DODAP. Densities were calculated using one representative atom per lipid (oxygen for cholesterol, nitrogen for the aminolipid, and phosphorus for the helper lipid). Positions are reported relative to the membrane surface, defined as the average z-position of the phosphorus atoms in the helper lipid headgroups of each leaflet. Negative distances (d) correspond to the membrane core, whereas positive distances correspond to the solvent phase. Results were averaged over both leaflets.

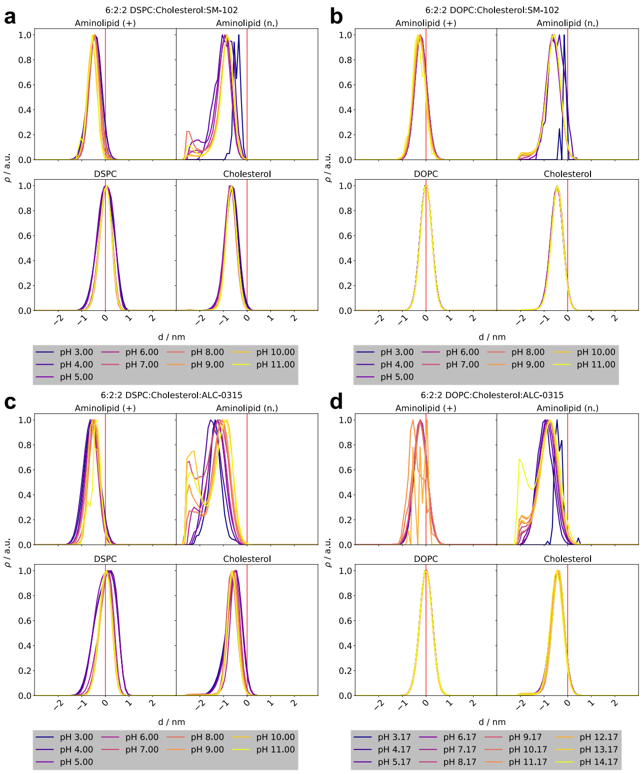


**Fig. 48** Lipid density profiles along the z-axis for membranes with a 6:2:2 helper lipid:cholesterol:aminolipid composition containing SM-102 and ALC-0315. Densities were calculated using one representative atom per lipid (oxygen for cholesterol, nitrogen for the aminolipid, and phosphorus for the helper lipid). Positions are reported relative to the membrane surface, defined as the average z-position of the phosphorus atoms in the helper lipid headgroups of each leaflet. Negative distances (d) correspond to the membrane core, whereas positive distances correspond to the solvent phase. Results were averaged over both leaflets.

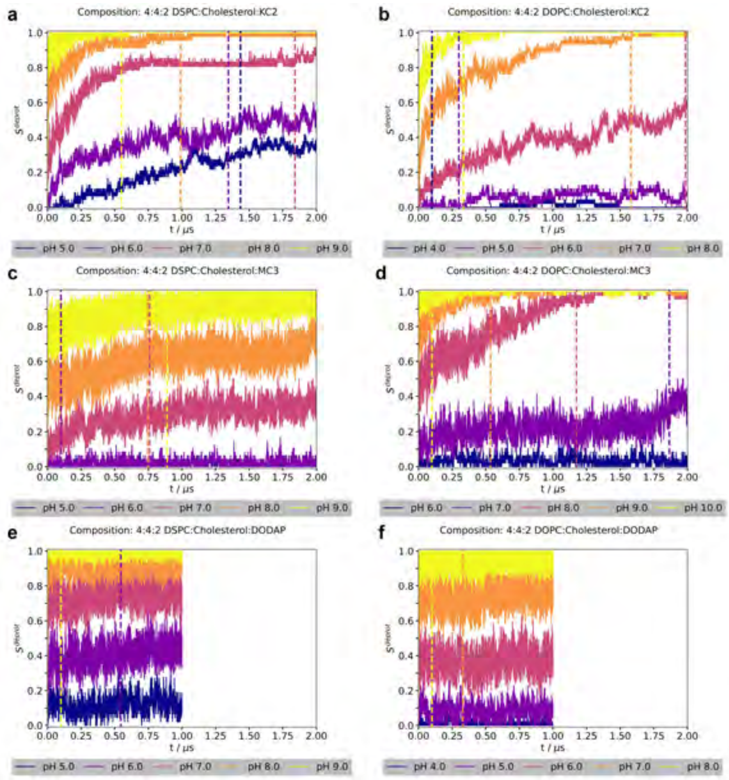


**Fig. 49** Fraction of deprotonated aminolipids ($S^{deprot}$) per frame for membranes with a 4:4:2 composition (DOPC/DSPC:cholesterol:KC2/MC3/DODAP) over the full trajectory length. Vertical dashed lines indicate the equilibration time determined for each pH value.

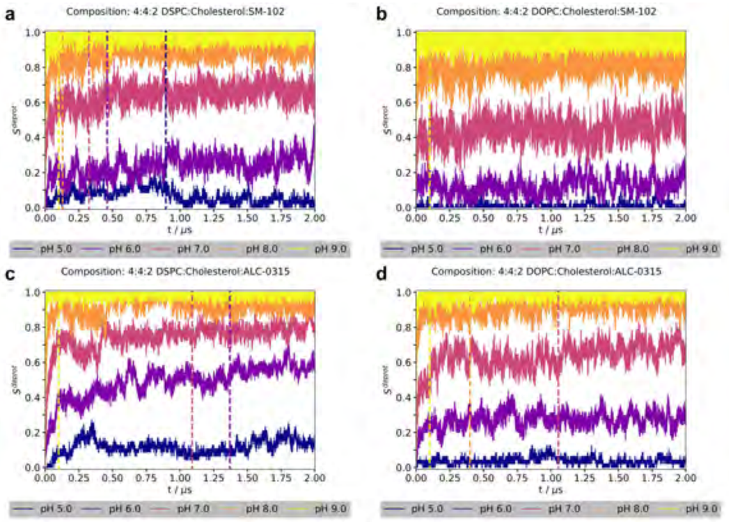


**Fig. 50** Fraction of deprotonated aminolipids ($S^{deprot}$) per frame for membranes with a 4:4:2 composition (DOPC/DSPC:cholesterol:SM-102/ALC-0315 over the full trajectory length. Vertical dashed lines indicate the equilibration time determined for each pH value.

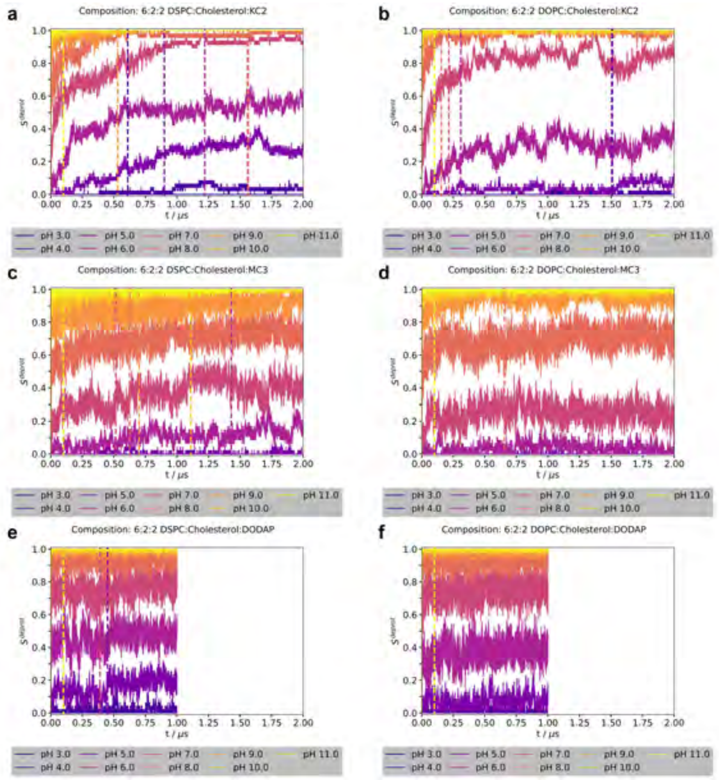


**Fig. 51** Fraction of deprotonated aminolipids ($S^{deprot}$) per frame for membranes with a 6:2:2 composition (DOPC/DSPC:cholesterol:KC2/MC3/DODAP) over the full trajectory length. Vertical dashed lines indicate the equilibration time determined for each pH value.

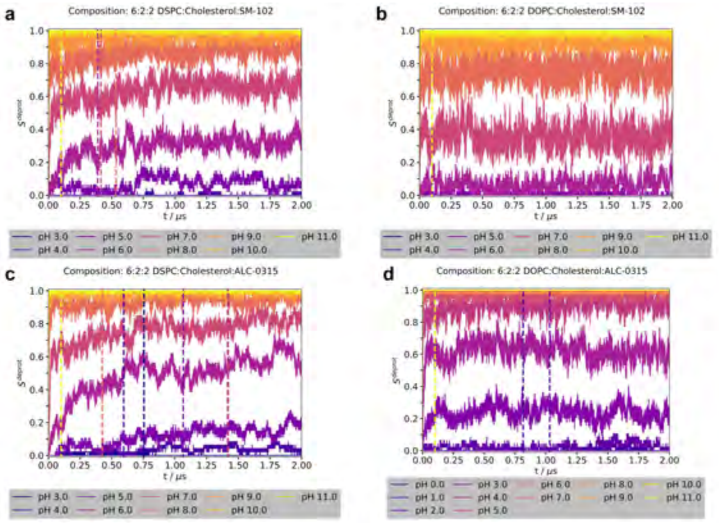


**Fig. 52** Fraction of deprotonated aminolipids ($S^{deprot}$) per frame for membranes with a 6:2:2 composition (DOPC/DSPC:cholesterol:SM-102/ALC-0315) over the full trajectory length. Vertical dashed lines indicate the equilibration time determined for each pH value.

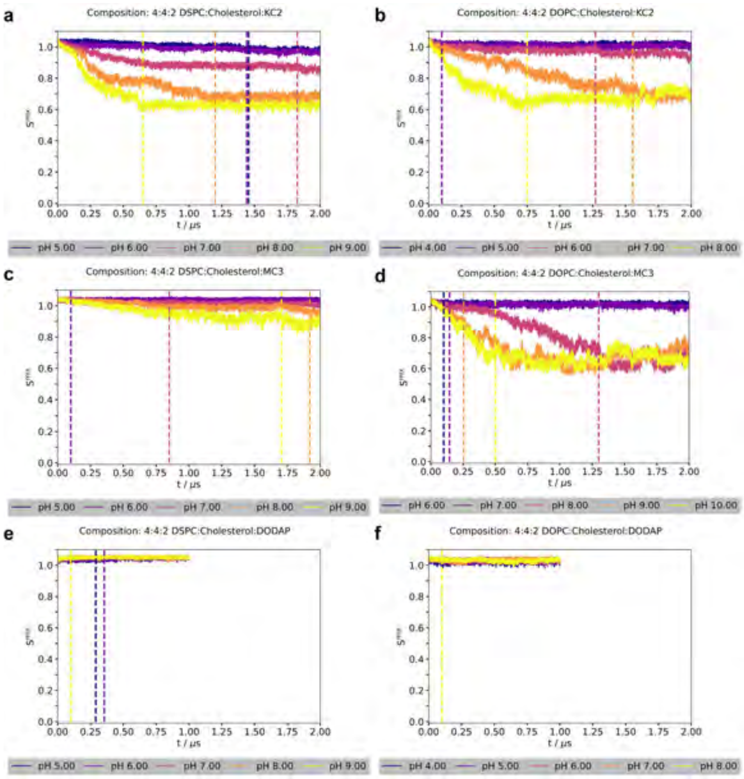


**Fig. 53** Conditional mixing entropy ($S^{mix}$) per frame for membranes with a 4:4:2 composition (DOPC/DSPC:cholesterol:KC2/MC3/DODAP) over the full trajectory length. Vertical dashed lines indicate the equilibration time determined for each pH value. Conditional mixing entropies were calculated following Brandani *et al.* [1], using a 1 nm cutoff to define neighboring lipids. Contacts were defined based on the distance between reference atoms in the headgroups of the target lipids: the titratable nitrogen for aminolipids, the oxygen atom for cholesterol, and the sn-2 carbon atom for phospholipids.

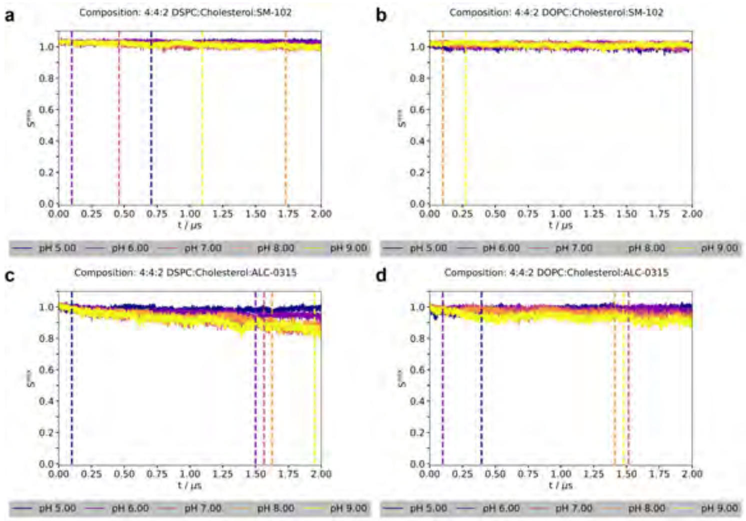


**Fig. 54** Conditional mixing entropy ($S^{mix}$) per frame for membranes with a 4:4:2 composition (DOPC/DSPC:cholesterol:SM-102/ALC-0315) over the full trajectory length. Vertical dashed lines indicate the equilibration time determined for each pH value. Conditional mixing entropies were calculated following Brandani *et al.* [1], using a 1 nm cutoff to define neighboring lipids. Contacts were defined based on the distance between reference atoms in the headgroups of the target lipids: the titratable nitrogen for aminolipids, the oxygen atom for cholesterol, and the sn-2 carbon atom for phospholipids.

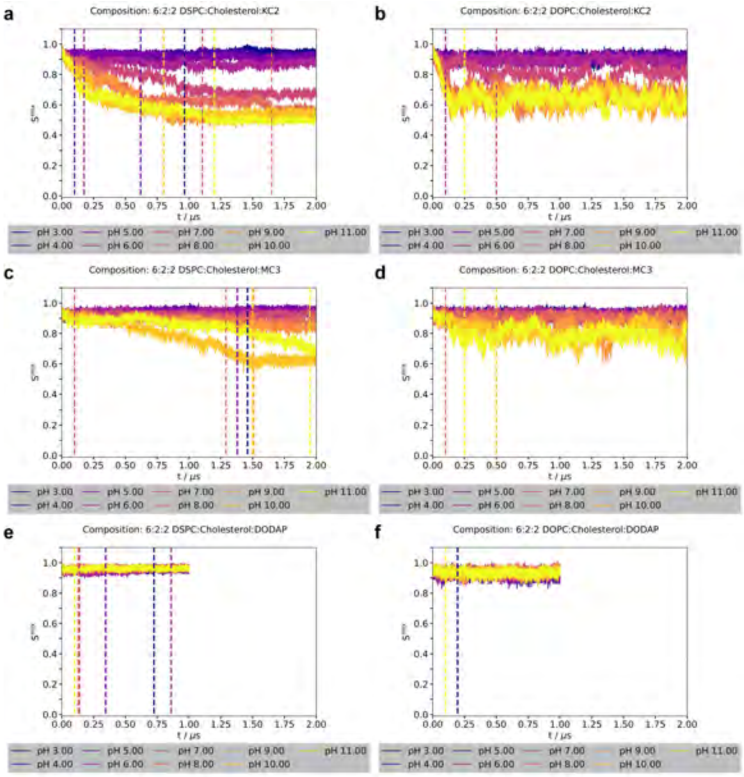


**Fig. 55** Conditional mixing entropy ($S^{mix}$) per frame for membranes with a 6:2:2 composition (DOPC/DSPC:cholesterol:KC2/MC3/DODAP) over the full trajectory length. Vertical dashed lines indicate the equilibration time determined for each pH value. Conditional mixing entropies were calculated following Brandani *et al.* [1], using a 1 nm cutoff to define neighboring lipids. Contacts were defined based on the distance between reference atoms in the headgroups of the target lipids: the titratable nitrogen for aminolipids, the oxygen atom for cholesterol, and the sn-2 carbon atom for phospholipids.

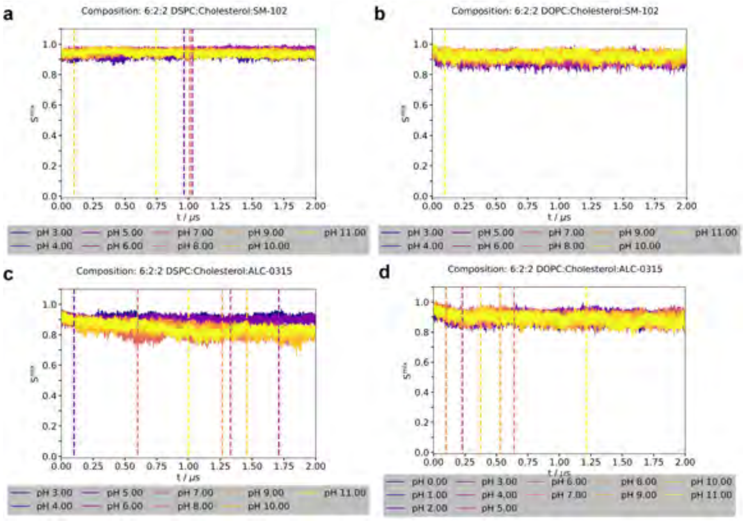


**Fig. 56** Conditional mixing entropy ($S^{mix}$) per frame for membranes with a 6:2:2 composition (DOPC/DSPC:cholesterol:SM-102/ALC-0315) over the full trajectory length. Vertical dashed lines indicate the equilibration time determined for each pH value. Conditional mixing entropies were calculated following Brandani *et al.* [1], using a 1 nm cutoff to define neighboring lipids. Contacts were defined based on the distance between reference atoms in the headgroups of the target lipids: the titratable nitrogen for aminolipids, the oxygen atom for cholesterol, and the sn-2 carbon atom for phospholipids.

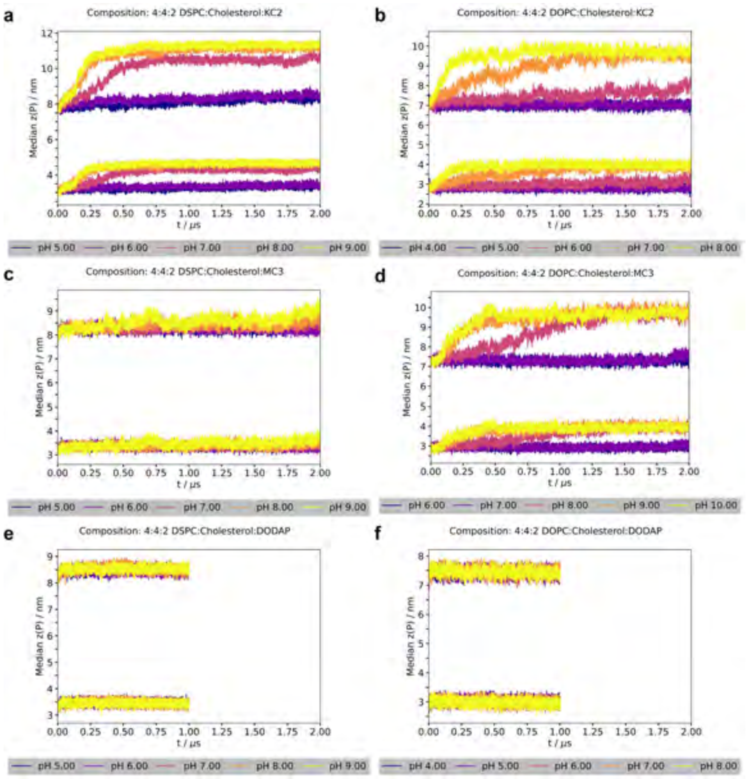


**Fig. 57** Average z-position of the phosphorus atoms in the upper and lower leaflets per frame for systems with a 4:4:2 composition (DOPC/DSPC:cholesterol:KC2/MC3/DODAP) over the full trajectory length. Average z-positions were computed using a periodic trigonometric approach (see Methods) to mitigate artifacts arising when the membrane is split across periodic boundary conditions.

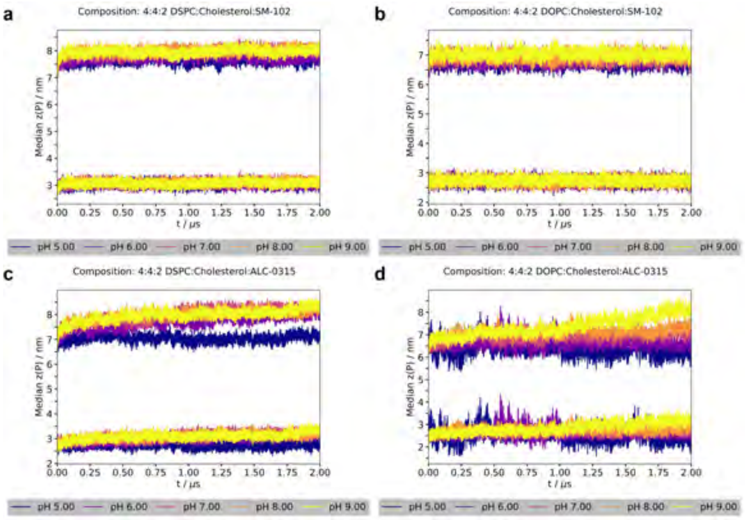


**Fig. 58** Average z-position of the phosphorus atoms in the upper and lower leaflets per frame for systems with a 4:4:2 composition (DOPC/DSPC:cholesterol:SM-102/ALC-0315) over the full trajectory length. Average z-positions were computed using a periodic trigonometric approach (see Methods) to mitigate artifacts arising when the membrane is split across periodic boundary conditions.

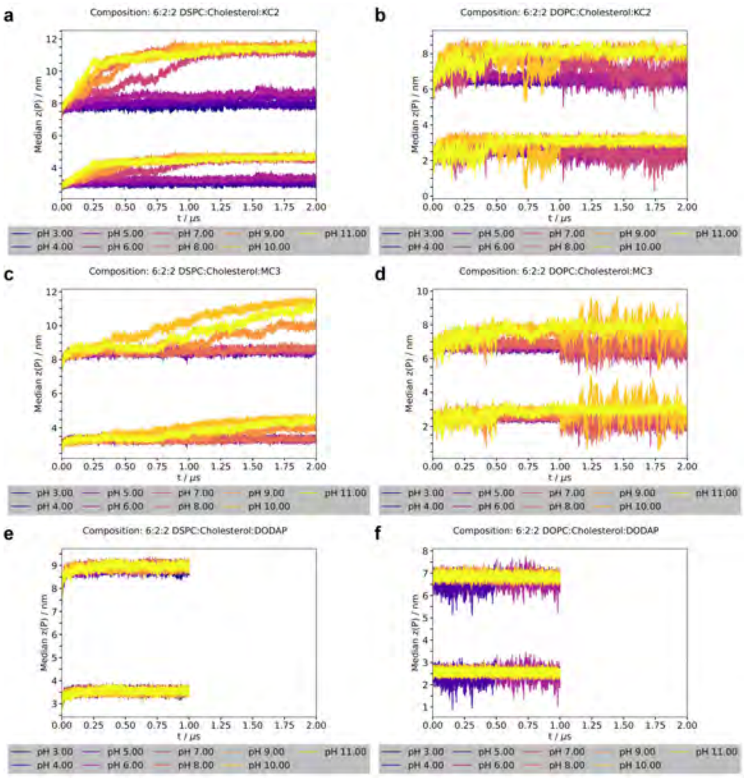


**Fig. 59** Average z-position of the phosphorus atoms in the upper and lower leaflets per frame for systems with a 6:2:2 composition (DOPC/DSPC:cholesterol:KC2/MC3/DODAP) over the full trajectory length. Average z-positions were computed using a periodic trigonometric approach (see Methods) to mitigate artifacts arising when the membrane is split across periodic boundary conditions.

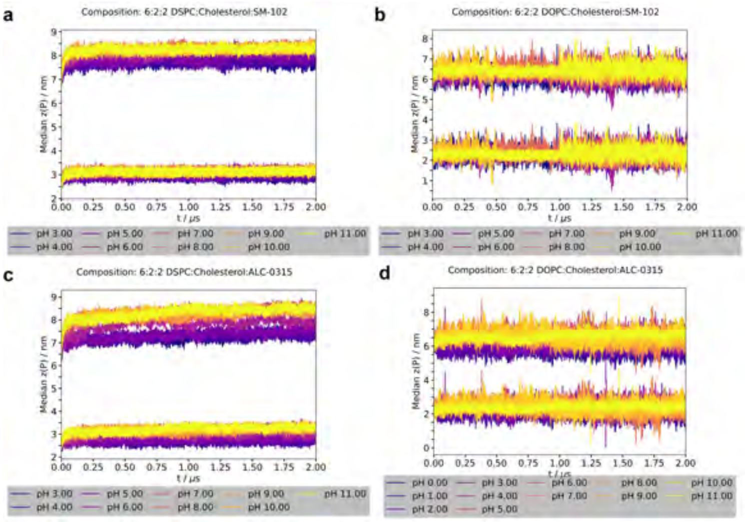


**Fig. 60** Average z-position of the phosphorus atoms in the upper and lower leaflets per frame for systems with a 6:2:2 composition (DOPC/DSPC:cholesterol:SM-102/ALC-0315) over the full trajectory length. Average z-positions were computed using a periodic trigonometric approach (see Methods) to mitigate artifacts arising when the membrane is split across periodic boundary conditions.

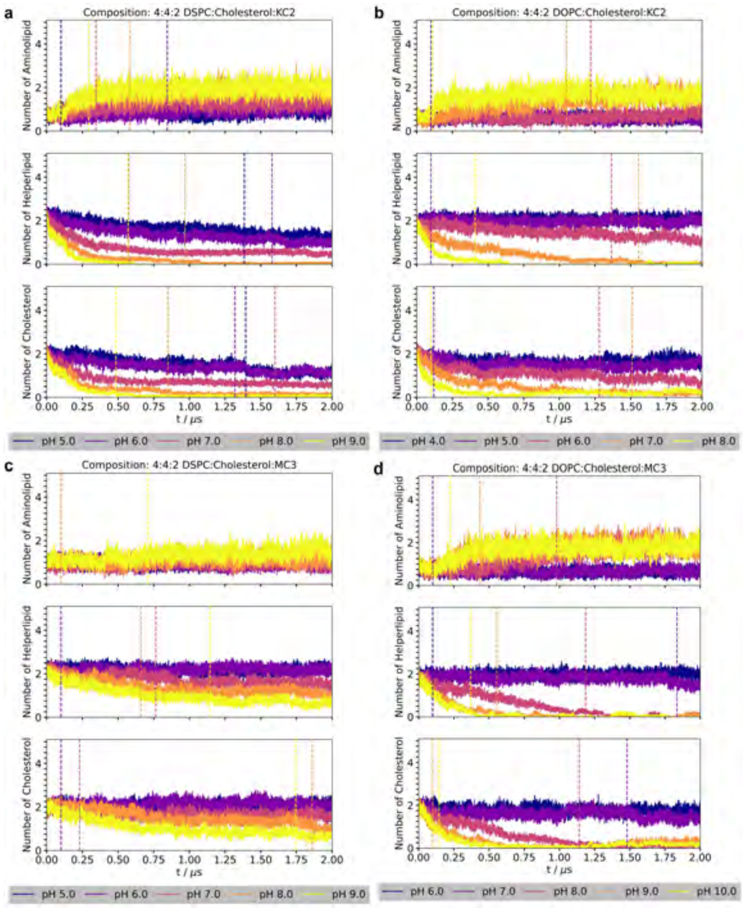


**Fig. 61** Number of neighboring lipids per aminolipid per frame for membranes with a 4:4:2 composition (DOPC/DSPC:cholesterol:KC2/MC3) over the full trajectory length. Vertical dashed lines indicate the equilibration time determined for each pH value. A cutoff distance of 1 nm was used to define neighboring lipids. Contacts were defined based on the distance between reference atoms in the lipid headgroups: the titratable nitrogen atom for aminolipids, the oxygen atom for cholesterol, and the sn-2 carbon atom for phospholipids.

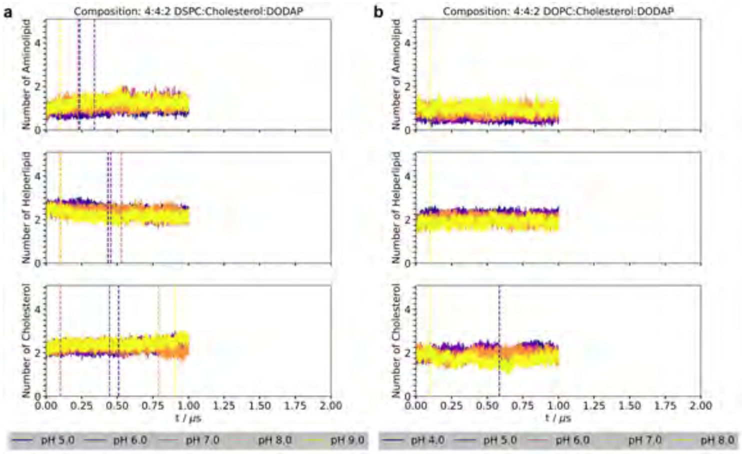


**Fig. 62** Number of neighboring lipids per aminolipid per frame for membranes with a 4:4:2 composition (DOPC/DSPC:cholesterol:DODAP) over the full trajectory length. Vertical dashed lines indicate the equilibration time determined for each pH value. A cutoff distance of 1 nm was used to define neighboring lipids. Contacts were defined based on the distance between reference atoms in the lipid headgroups: the titratable nitrogen atom for aminolipids, the oxygen atom for cholesterol, and the sn-2 carbon atom for phospholipids.

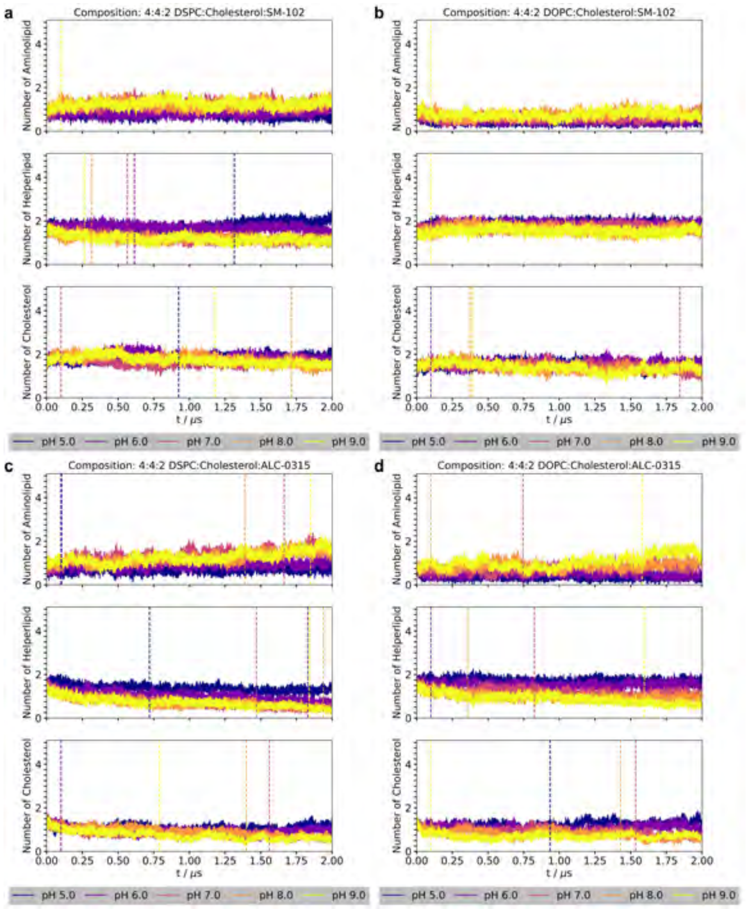


**Fig. 63** Number of neighboring lipids per aminolipid per frame for membranes with a 4:4:2 composition (DOPC/DSPC:cholesterol:SM-102/ALC-0315) over the full trajectory length. Vertical dashed lines indicate the equilibration time determined for each pH value. A cutoff distance of 1 nm was used to define neighboring lipids. Contacts were defined based on the distance between reference atoms in the lipid headgroups: the titratable nitrogen atom for aminolipids, the oxygen atom for cholesterol, and the sn-2 carbon atom for phospholipids.

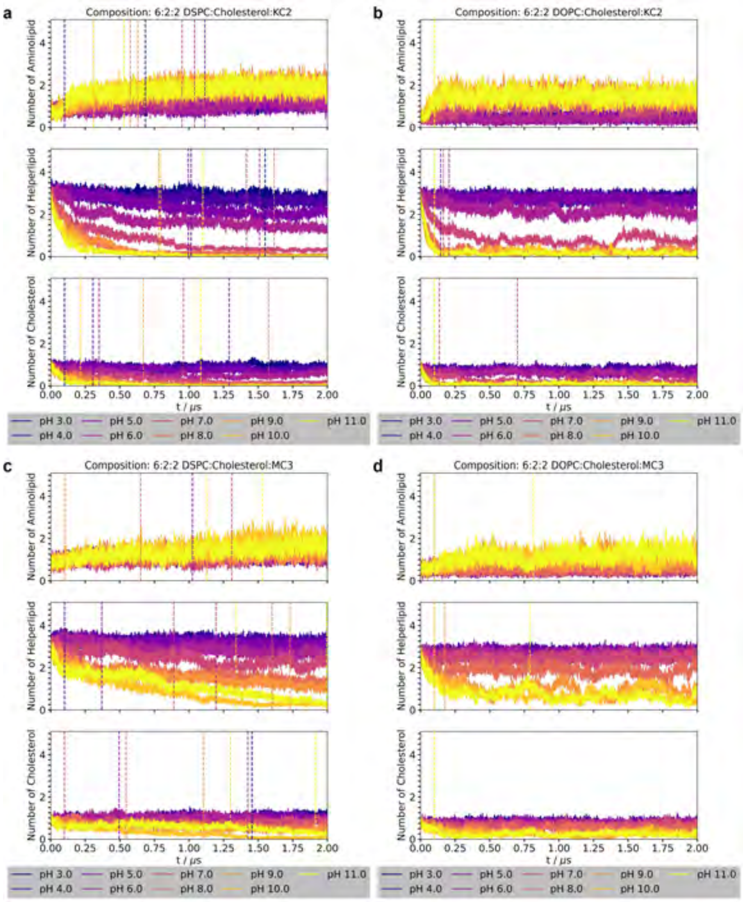


**Fig. 64** Number of neighboring lipids per aminolipid per frame for membranes with a 6:2:2 composition (DOPC/DSPC:cholesterol:KC2/MC3) over the full trajectory length. Vertical dashed lines indicate the equilibration time determined for each pH value. A cutoff distance of 1 nm was used to define neighboring lipids. Contacts were defined based on the distance between reference atoms in the lipid headgroups: the titratable nitrogen atom for aminolipids, the oxygen atom for cholesterol, and the sn-2 carbon atom for phospholipids.

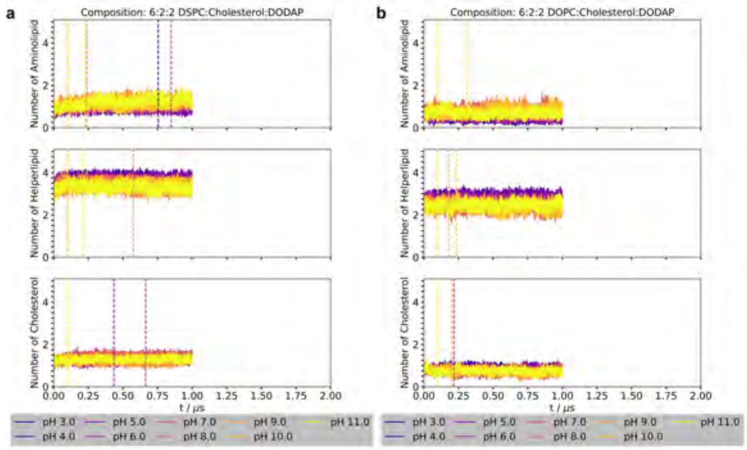


**Fig. 65** Number of neighboring lipids per aminolipid per frame for membranes with a 6:2:2 composition (DOPC/DSPC:cholesterol:DODAP) over the full trajectory length. Vertical dashed lines indicate the equilibration time determined for each pH value. A cutoff distance of 1 nm was used to define neighboring lipids. Contacts were defined based on the distance between reference atoms in the lipid headgroups: the titratable nitrogen atom for aminolipids, the oxygen atom for cholesterol, and the sn-2 carbon atom for phospholipids.

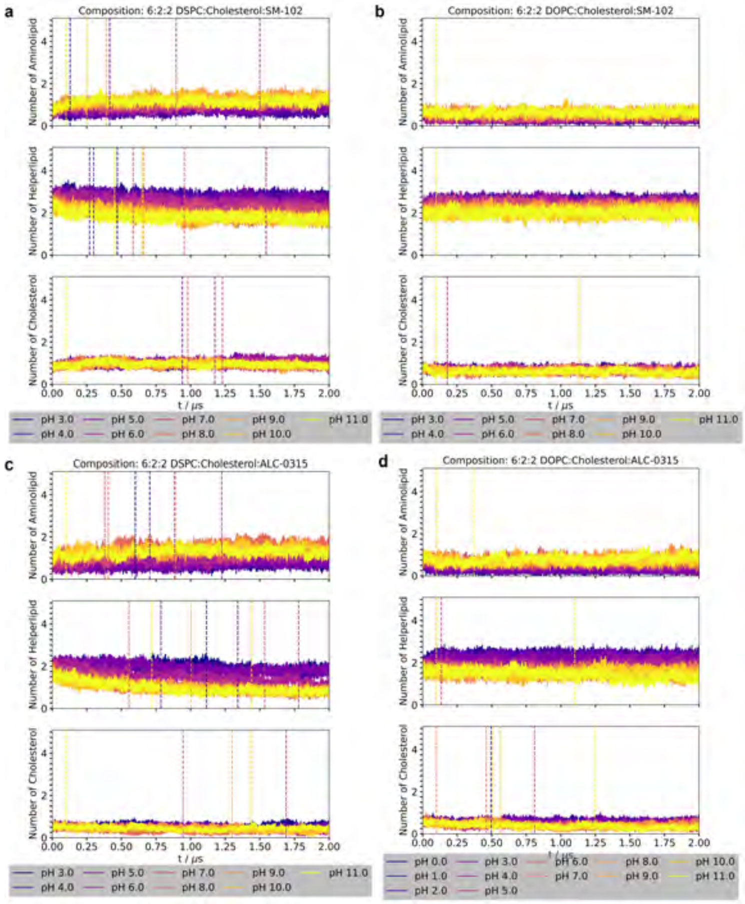


**Fig. 66** Number of neighboring lipids per aminolipid per frame for membranes with a 6:2:2 composition (DOPC/DSPC:cholesterol:SM-102/ALC-0315) over the full trajectory length. Vertical dashed lines indicate the equilibration time determined for each pH value. A cutoff distance of 1 nm was used to define neighboring lipids. Contacts were defined based on the distance between reference atoms in the lipid headgroups: the titratable nitrogen atom for aminolipids, the oxygen atom for cholesterol, and the sn-2 carbon atom for phospholipids.

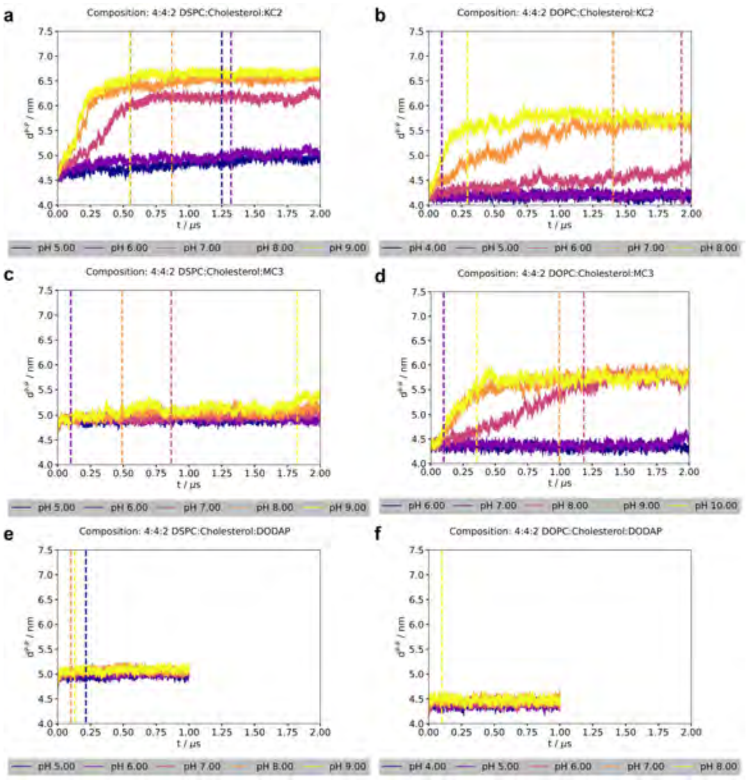


**Fig. 67** Membrane thickness per frame for systems with a 4:4:2 composition (DOPC/DSPC:cholesterol:KC2/MC3/DODAP) over the full trajectory length. Thickness is defined as the distance between the average z-positions of the phosphorus atoms in the upper and lower leaflets. Vertical dashed lines indicate the equilibration time determined for each pH value. Average z-positions were computed using a periodic trigonometric approach (see Methods) to mitigate artifacts arising when the membrane is split across periodic boundary conditions.

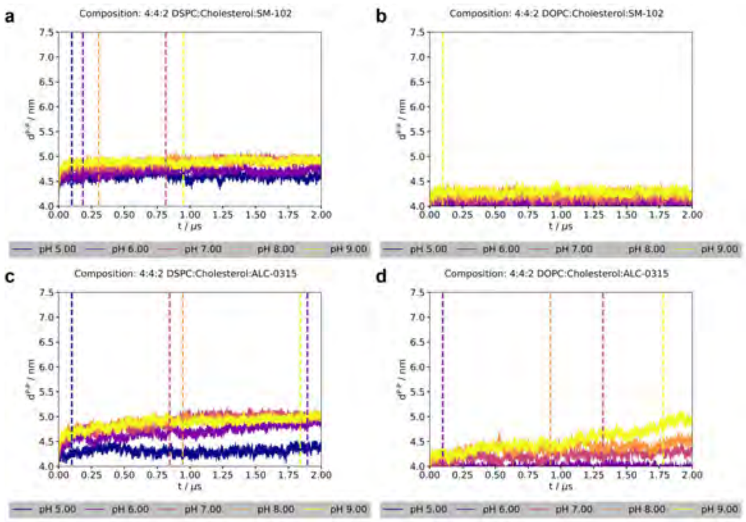


**Fig. 68** Membrane thickness per frame for systems with a 4:4:2 composition (DOPC/DSPC:cholesterol:SM-102/ALC-0315) over the full trajectory length. Thickness is defined as the distance between the average z-positions of the phosphorus atoms in the upper and lower leaflets. Vertical dashed lines indicate the equilibration time determined for each pH value. Average z-positions were computed using a periodic trigonometric approach (see Methods) to mitigate artifacts arising when the membrane is split across periodic boundary conditions.

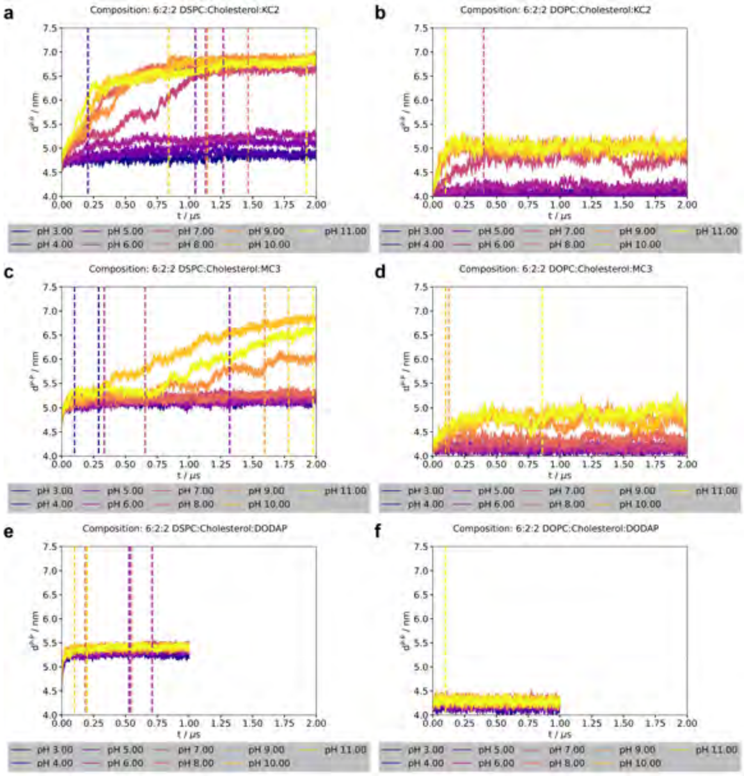


**Fig. 69** Membrane thickness per frame for systems with a 6:2:2 composition (DOPC/DSPC:cholesterol:KC2/MC3/DODAP) over the full trajectory length. Thickness is defined as the distance between the average z-positions of the phosphorus atoms in the upper and lower leaflets. Vertical dashed lines indicate the equilibration time determined for each pH value. Average z-positions were computed using a periodic trigonometric approach (see Methods) to mitigate artifacts arising when the membrane is split across periodic boundary conditions.

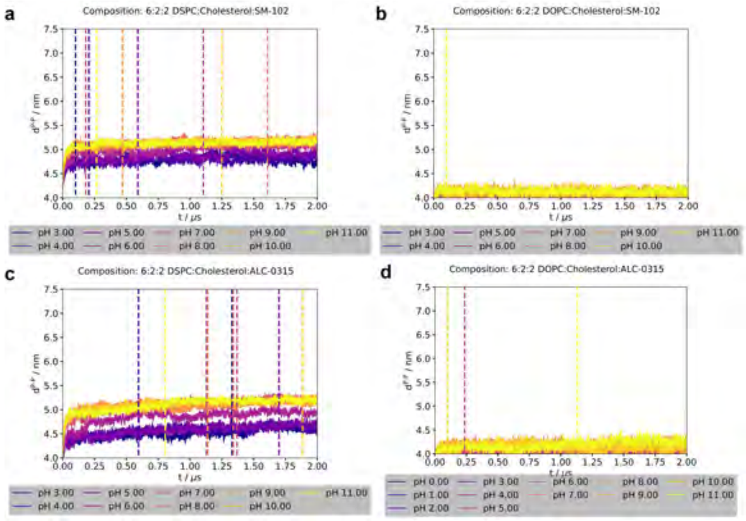


**Fig. 70** Membrane thickness per frame for systems with a 6:2:2 composition (DOPC/DSPC:cholesterol:SM-102/ALC-0315) over the full trajectory length. Thickness is defined as the distance between the average z-positions of the phosphorus atoms in the upper and lower leaflets. Vertical dashed lines indicate the equilibration time determined for each pH value. Average z-positions were computed using a periodic trigonometric approach (see Methods) to mitigate artifacts arising when the membrane is split across periodic boundary conditions.

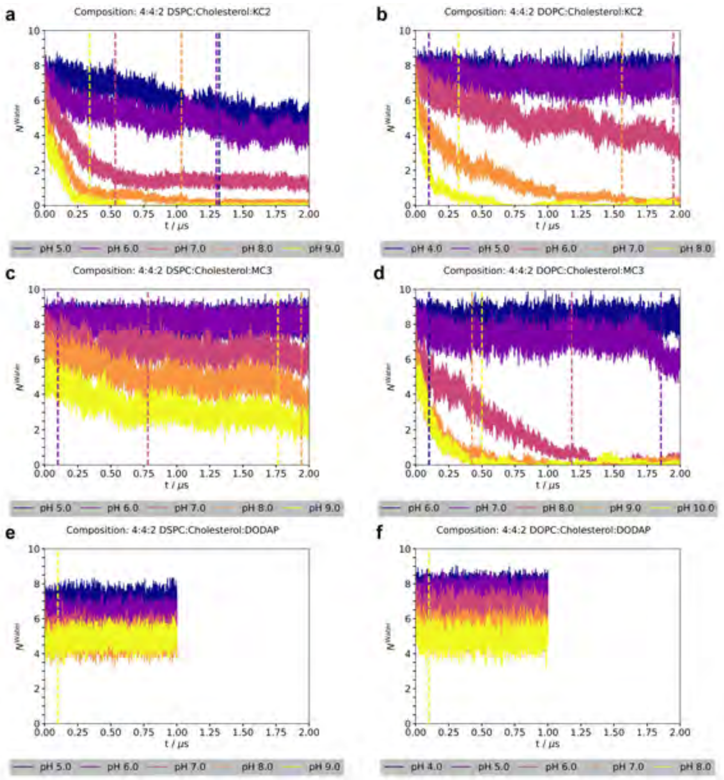


**Fig. 71** Average number of water molecules within 0.5 nm of the titratable headgroups ($\rho_n^{\mathrm{Water}}$) per frame for membranes with a 4:4:2 composition (DOPC/DSPC:cholesterol:KC2/MC3/DODAP) over the full trajectory length. Vertical dashed lines indicate the equilibration time determined for each pH value. The reference atom of the aminolipid was the hydrogen atom of the titratable headgroup, whereas the oxygen atom was used as the reference for water.

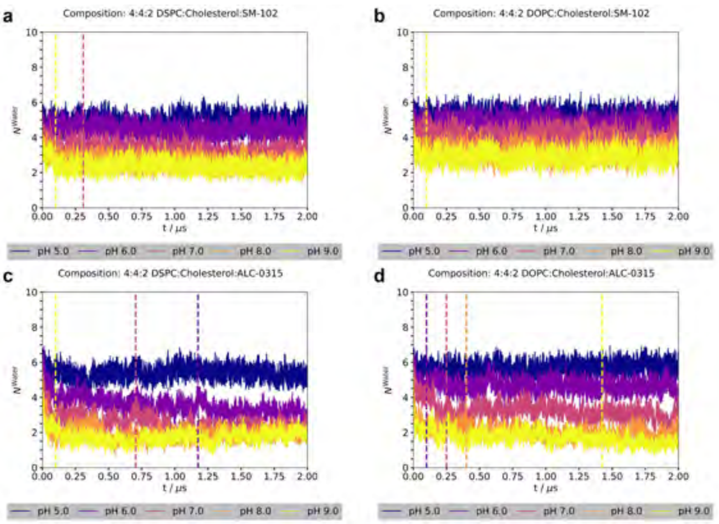


**Fig. 72** Average number of water molecules within 0.5 nm of the titratable headgroups ($\rho_n^{\mathrm{Water}}$) per frame for membranes with a 4:4:2 composition (DOPC/DSPC:cholesterol:SM-102/ALC-0315) over the full trajectory length. Vertical dashed lines indicate the equilibration time determined for each pH value. The reference atom of the aminolipid was the hydrogen atom of the titratable headgroup, whereas the oxygen atom was used as the reference for water.

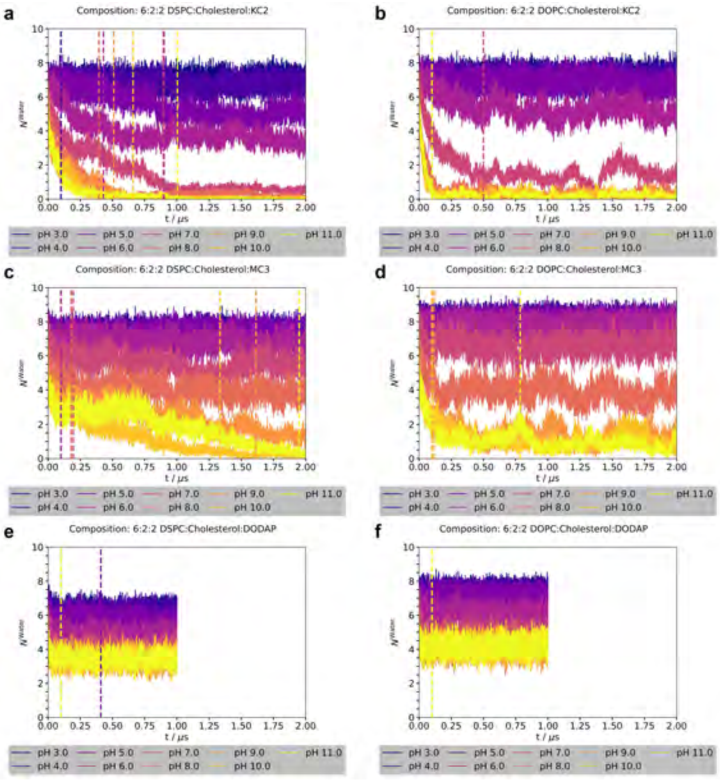


**Fig. 73** Average number of water molecules within 0.5 nm of the titratable headgroups ($\rho_n^{\mathrm{Water}}$) per frame for membranes with a 6:2:2 composition (DOPC/DSPC:cholesterol:KC2/MC3/DODAP) over the full trajectory length. Vertical dashed lines indicate the equilibration time determined for each pH value. The reference atom of the aminolipid was the hydrogen atom of the titratable headgroup, whereas the oxygen atom was used as the reference for water.

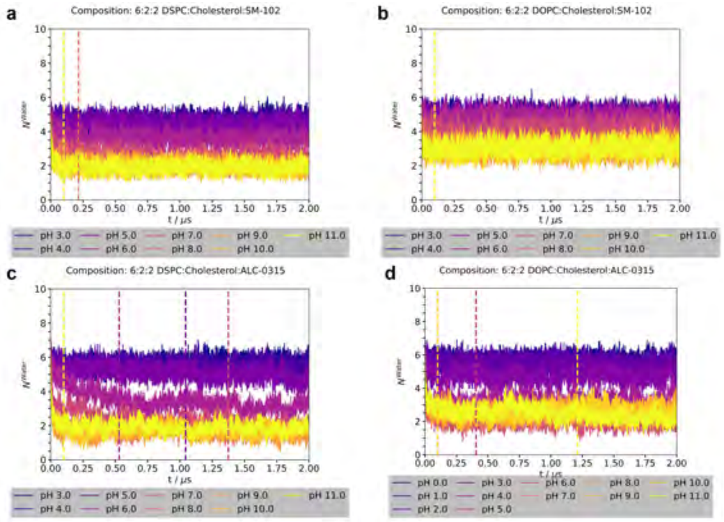


**Fig. 74** Average number of water molecules within 0.5 nm of the titratable headgroups ($\rho_n^{\text{Water}}$) per frame for membranes with a 6:2:2 composition (DOPC/DSPC:cholesterol:SM-102/ALC-0315) over the full trajectory length. Vertical dashed lines indicate the equilibration time determined for each pH value. The reference atom of the aminolipid was the hydrogen atom of the titratable headgroup, whereas the oxygen atom was used as the reference for water.

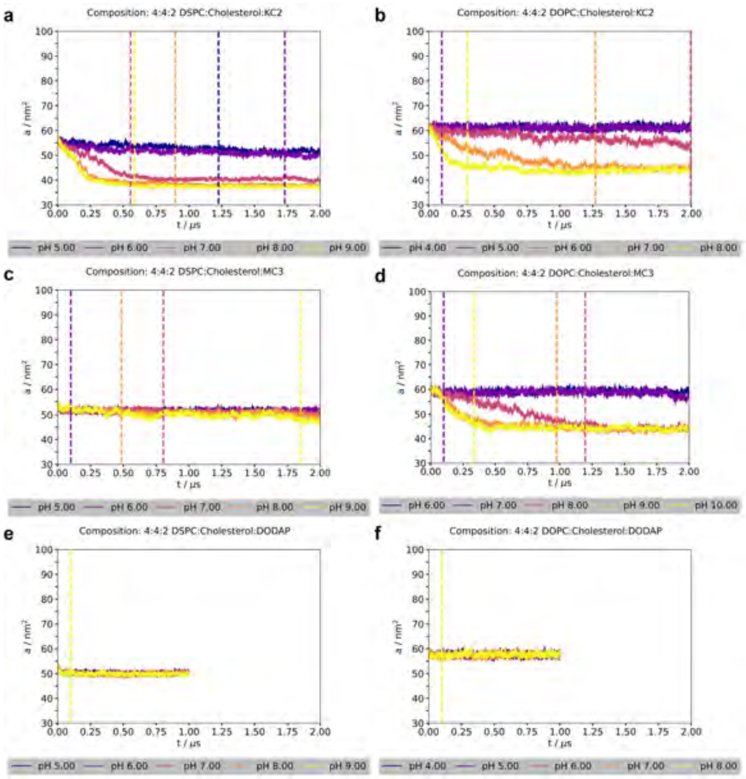


**Fig. 75** Area per frame for membranes with a 4:4:2 composition (DOPC/D-SPC:cholesterol:KC2/MC3/DODAP) over the full trajectory length. Vertical dashed lines indicate the equilibration time determined for each pH value.

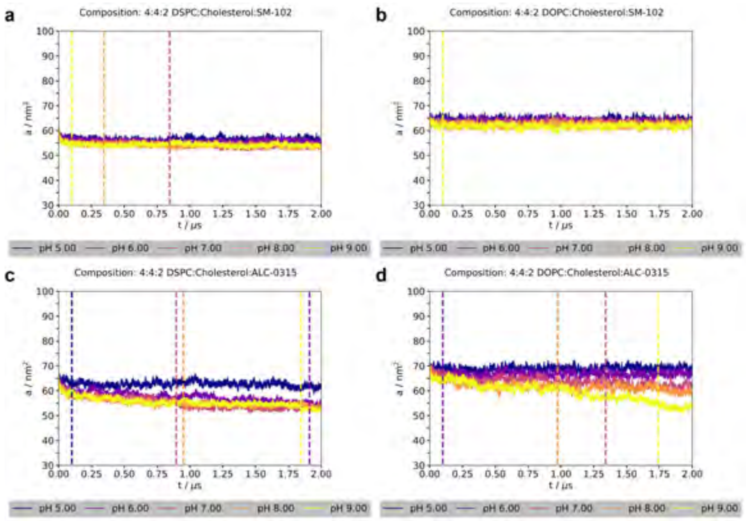


**Fig. 76** Area per frame for membranes with a 4:4:2 composition (DOPC/DSPC:cholesterol:SM-102/ALC-0315) over the full trajectory length. Vertical dashed lines indicate the equilibration time determined for each pH value.

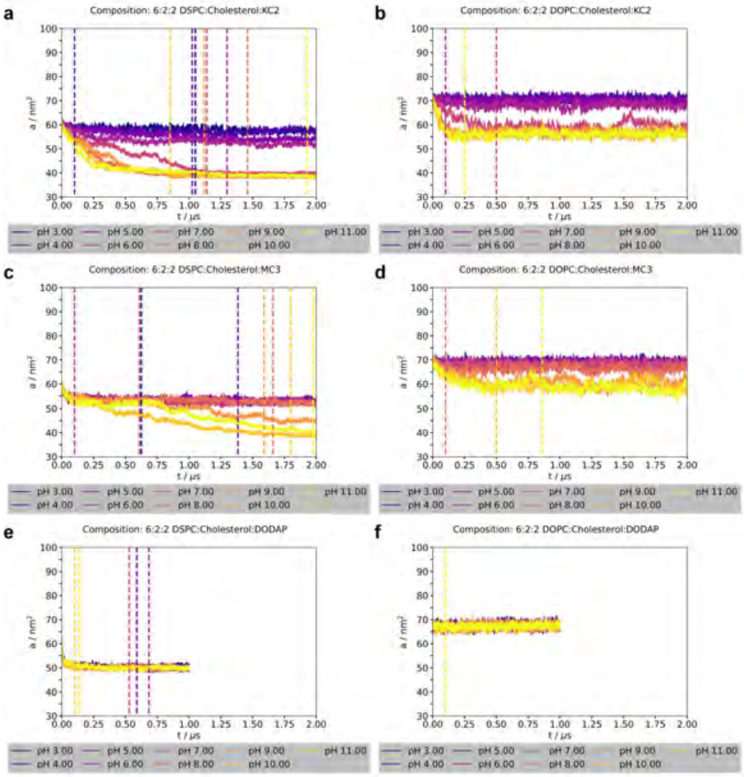


**Fig. 77** Area per frame for membranes with a 6:2:2 composition (DOPC/DSPC:cholesterol:KC2/MC3/DODAP) over the full trajectory length. Vertical dashed lines indicate the equilibration time determined for each pH value.

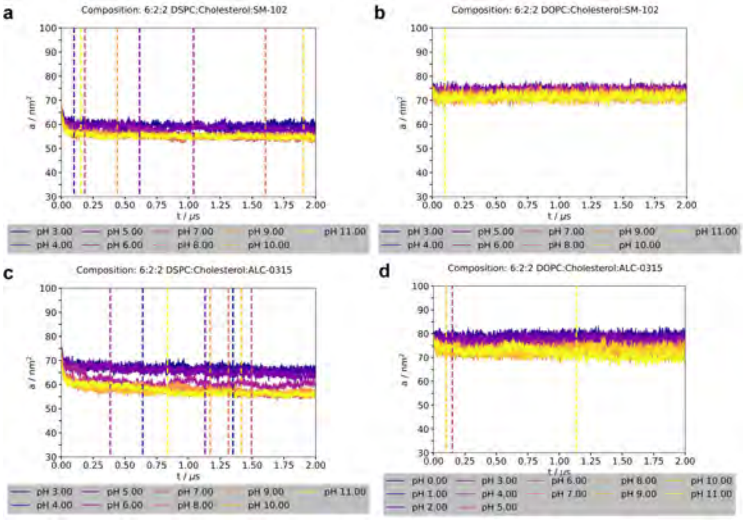


**Fig. 78** Area per frame for membranes with a 6:2:2 composition (DOPC/DSPC:cholesterol:SM-102/ALC-0315) over the full trajectory length. Vertical dashed lines indicate the equilibration time determined for each pH value.

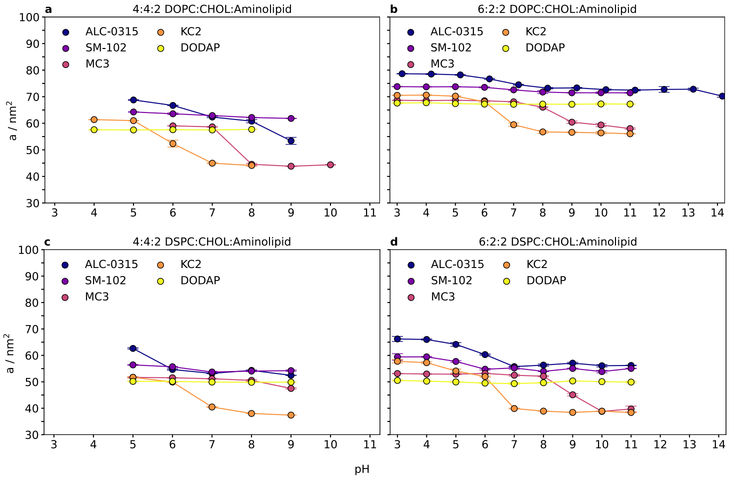


**Fig. 79** Average membrane surface area per leaflet as a function of pH. Mean values of the surface area are shown for DOPC-based (top) and DSPC-based (bottom) membranes containing **a,c** 40 mol% cholesterol and **b,d** 20 mol% cholesterol.
